# Genetics of 38 blood and urine biomarkers in the UK Biobank

**DOI:** 10.1101/660506

**Authors:** Nasa Sinnott-Armstrong, Yosuke Tanigawa, David Amar, Nina J. Mars, Matthew Aguirre, Guhan Ram Venkataraman, Michael Wainberg, Hanna M. Ollila, James P. Pirruccello, Junyang Qian, Anna Shcherbina, FinnGen, Fatima Rodriguez, Themistocles L. Assimes, Vineeta Agarwala, Robert Tibshirani, Trevor Hastie, Samuli Ripatti, Jonathan K. Pritchard, Mark J. Daly, Manuel A. Rivas

## Abstract

Clinical laboratory tests are a critical component of the continuum of care and provide a means for rapid diagnosis and monitoring of chronic disease. In this study, we systematically evaluated the genetic basis of 38 blood and urine laboratory tests measured in 358,072 participants in the UK Biobank and identified 1,857 independent loci associated with at least one laboratory test, including 488 large-effect protein truncating, missense, and copy-number variants. We tested these loci for enrichment in specific single cell types in kidney, liver, and pancreas relevant to disease aetiology. We then causally linked the biomarkers to medically relevant phenotypes through genetic correlation and Mendelian randomization. Finally, we developed polygenic risk scores (PRS) for each biomarker and built multi-PRS models using all 38 PRSs simultaneously. We found substantially improved prediction of incidence in FinnGen (n=135,500) with the multi-PRS relative to single-disease PRSs for renal failure, myocardial infarction, liver fat percentage, and alcoholic cirrhosis. Together, our results show the genetic basis of these biomarkers, which tissues contribute to the biomarker function, the causal influences of the biomarkers, and how we can use this to predict disease.

## Introduction

Serum and urine biomarker measurements are the primary clinical tools for diagnosing adverse health conditions and monitoring treatment response. As such, understanding the genetic predisposition to particular phenotype measurements, and the factors that confound them, has implications for disease treatment. While the genetics of some biomarkers have been extensively studied, most notably lipids^1,2,3^, glycaemic traits ^4–6^, and measurements of kidney function^7–9^, most biomarkers have only had limited measured genetic contribution.

To this end, UK Biobank (UKB) has performed lab testing of >30 proteins, metabolites, and protein modifications in serum and urine on a cohort of >480,000 individuals with extensive prior phenotype and genome-wide genotype data including ∼360,000 unrelated individuals (**Supplementary Figure 1**).

Here, we 1) performed a systematic characterization of genetic architecture in ∼360,000 individuals including protein-altering, protein-truncating, non-coding, human leukocyte antigen (HLA), and copy number variants; 2) built phenome-wide associations for implicated genetic variants; 3) evaluated causal relationships with 56 diseases and 90 clinically relevant phenotypes; and 4) constructed prediction models from genome data.

## Results

### Biomarker phenotype distributions

We aimed to examine the consistency of the measurements themselves. Despite extensive quality control and significant effort to avoid batch effects, even subtle deviations from expectation can be highly significant in biobank-scale datasets^10^. First, we assessed the impact of medication status for individuals that were on lipid-lowering medications at baseline (2006-2010) by estimating changes in biomarker measurements between visits (a subset of 20,000 individuals that returned for a repeated assessment, **Methods**). Overall, the estimated adjustments typically agreed with literature estimates^3^ (**Supplementary Table 1**). After adjusting for medication status, we fit a regression model for 127 covariates including age, sex, urine and serum processing metrics, fasting time, and estimated sample dilution factor, along with numerous relevant interactions between covariates (see **Methods**). These covariates explained a range of 1% (Rheumatoid factor) to 81% (Testosterone) of the phenotypic variance observed (**Supplementary Figures 2A-C**). Because oestradiol, microalbumin in urine, and rheumatoid factor had a high proportion of values below the lower reportable range (80%, 68%, and 91%, respectively), which is to be expected given the age range and health status of the UK Biobank population, we considered these values as ‘naturally low’ rather than missing, and treated these phenotypes as binary if they were above certain levels (higher than 212 pmol/L for oestradiol^11^, higher than 40mg/L for microalbumin in urine, and higher than 16 IU/mL for rheumatoid factor)^12^. Furthermore, we derived the urine albumin-to-creatinine ratio (UACR), which is indicative of chronic kidney disease when higher than 30 mg/g^13^. Taking all the 38 lab phenotypes together, we recover previously estimated phenotype correlations (**Supplementary Table 2, Supplementary Figure 3**)^14,15^.

### Comparison of self-reported, diagnosed, medication, and lab-derived disease status

When comparing between studies, disease status is often evaluated in different ways and it is a challenge to reconcile these differences. In addition to biomarker measurements, UK Biobank has extensive self-reported disease and diagnosis, nurse interviews with participants, and inpatient and soon-to-be-released primary care diagnosis and medication codes. This affords a unique opportunity to evaluate the overlap between such measures in the definition of complex traits.

To examine the validity of our phenotypes and biomarkers, we used type 2 diabetes (T2D) and hemoglobin A1c (HbA1c) levels as our clinical outcome and biomarker, respectively. Type 2 diabetes is characterized by progressive loss of insulin sensitivity and is commonly diagnosed through hemoglobin A1c (HbA1c), a modification to red blood cells induced by long term exposure to high serum glucose. We compared self-reported and nurse-collected diagnosis, medication (sulfonylureas, metformin, and other oral antidiabetic drugs), and serum glucose and HbA1c as measures of diabetes to the previously published definition that was validated against a subset of individuals with available primary care data (Eastwood et al. 2016)^16^. As expected, HbA1c levels, regardless of residualization or adjustment for statins (**Methods**), were well correlated, and thresholded HbA1c (>48 mmol/mol or 6.5%, the clinical threshold for type 2 diabetes) was also similar (Pearson r = 0.72 with residualized, statin adjusted HbA1c). Glucose levels were not as predictive of T2D status as HbA1c, and medication status was similar to T2D itself, consistent with their use in the Eastwood et al. T2D definition (Pearson r = 0.79). Diagnosed T2D (defined by a nurse survey of participant diagnoses) was not similar to either the Eastwood et al. or biomarker measurements (maximum r = 0.48 with Eastwood et al. defined T2D, **Supplementary Figure 4, Supplementary Table 3**).

### Genetics of biomarkers

We performed association analysis between autosomal genetic variants and 38 biomarkers in 318,984 unrelated self-identified-White-British individuals of European ancestry and stratified the association into three bins: 1) protein-truncating (27,816), 2) protein-altering (87,407), and 3) non-coding (MAF > 1%, 1000 Genomes Phase 1 variants also present in Haplotype Reference Consortium [HRC], 9,444,561^17^) (**Figure 2A**). Comparison of effect sizes estimated across 42 comparison studies with 25 of the biomarkers show overall high agreement (correlation greater than 0.5 for 33 comparisons, **Supplementary Figure 5, Supplementary Table 4** for comparison) across previous studies of lipids^1,2,18,19^, glycaemic^20,21^, kidney function ^22,23^, liver function^20^, and other biomarkers measurements^24,25^.

**Figure 1.**
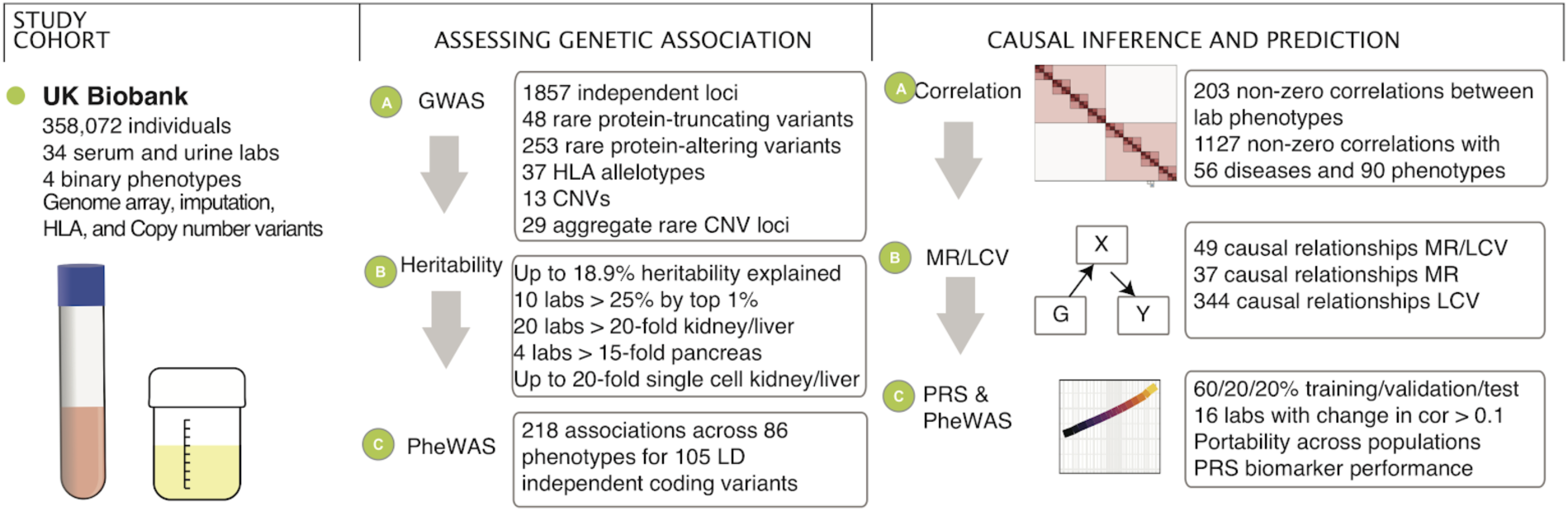
Schematic overview of the study. We prepared a dataset of 38 serum and urine biomarkers from 358,072 individuals in UK Biobank. From these data, we analyzed their genetic basis, assessed their relationship to disease outcomes and medically relevant phenotypes, and generated predictive models from genome data.

**Figure 2.**
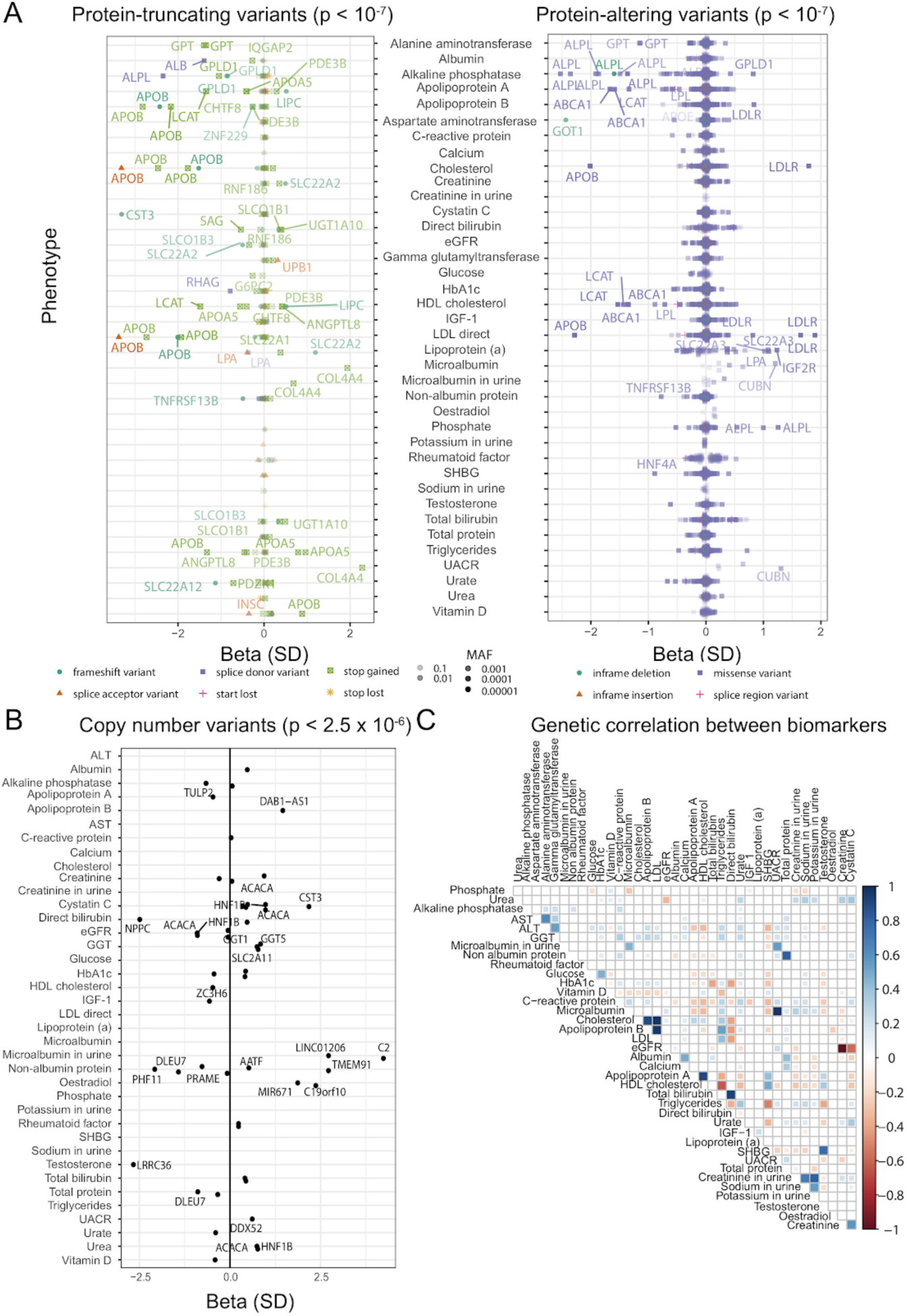
Genetics of biomarkers. **(A)** Summary of large-effect protein-truncating (abs(Beta) >= 0.25) and protein-altering variants (abs(Beta) >= 0.75). Every point plotted is independent and significantly associated with the given trait, and most variants have small effect, particularly for protein-altering variants; those with large effect are labeled. All variants are directly genotyped on the genotyping array, and effect betas are the number of standard deviations changed in the phenotype per alternative allele. Each set of variants is separated by trait but overlap of individual variants in the same gene between traits is present -- a more detailed table of individual hits and cascade plots are in Supplementary Tables 5-7 and Supplementary Figures 6-8. **(B)** Summary of large-effect burden of rare CNVs overlapping a gene and the tested biomarkers. Large effect variants (abs(Beta) >= 0.5) are labeled. **(C)** Correlation of genetic effects plot between the 38 lab phenotypes.

We adjusted the nominal association p values separately for each annotation bin using Bonferroni correction for multiple hypothesis testing and identified over 7,000 significant associations (p < 1 × 10^−7^ for coding variants [including protein-truncating and protein-altering], p < 5 × 10^−8^ for non-coding, posterior probability > 0.8 for model selection of associating with at least one biomarker phenotype for HLA alleles, **Supplementary Figures 6-9, Supplementary Tables 5-8**). LD Score intercepts for single-variant association results were between 0.929 and 1.33 for all 38 phenotypes, consistent with anthropometric traits in UKB and suggesting that population structure in our analysis is well-controlled^26^.

#### Biomarker associated variants prioritize therapeutic targets

Protein-altering variants that alter disease risk provide *in vivo* validation of therapeutic targets^27,28^. For example, recent studies suggest that molecular therapies targeting *ANGPTL3* and its encoded protein angiopoietin-like protein 3 have clinical potential comparable to therapies targeting *PCSK9* and its encoded protein proprotein convertase subtilisin/kexin type 9. By mainly affecting triglyceride-rich lipoproteins, ANGPTL3 inhibition might prove complementary to LDL cholesterol lowering with PCSK9 blockade^29,30^. These therapeutic hypotheses were catalyzed by the discovery of protein-altering variants in *ANGPTL3* and *PCSK9* that affect biomarker levels in humans^27,31–34^. In this study, we evaluated the relationship of predicted protein-truncating variants (PTVs) and protein-altering variants across the 38 biomarkers. We found 123 (48 rare, minor allele frequency [MAF] < 0.01) PTVs and 2,737 (253 rare) protein-altering alleles associations outside the major histocompatibility complex (MHC) (chr6:25,477,797-36,448,354; Bonferroni p < 1 × 10^−7^). In this study, we found 33 non-MHC PTVs (30 rare [MAF < 0.01]) with large estimated lowering effects (>0.1 sd) and 27 (22 rare) with large estimated raising effects (>0.1 sd, **Supplementary Table 5**) across at least one biomarker phenotype (**Figure 2A, Supplementary Figure 6**). Similarly, there were 264 (154 rare) and 202 (109 rare) non-MHC protein-altering variants with large estimated lowering and raising effects (>0.1 sd) across at least one biomarker phenotype, respectively (**Figure 2A, Supplementary Figure 7, Supplementary Table 6**).

For the eight cardiovascular biomarkers, we identified (**Supplementary Table 2**) three PTVs in *APOB* with documented protection against coronary artery disease and a range of strong effects on LDL cholesterol (1.9-3.4 sd), Apolipoprotein B (2.2-2.8 sd), and triglycerides (1.3 sd) in our dataset^35^; two PTVs in *LPA* with strong lowering effects on Lipoprotein A levels (0.36, 0.39 sd) and an estimated decreased risk of peripheral vascular disease (p = 0.0018; OR = 0.81 [0.70, 0.92]); a 0.2% MAF missense allele impacting the enzyme Acetyl-CoA carboxylase 2 (*ACACB*) with LDL, triglyceride, ApoB, and alkaline phosphatase-lowering effects with a trending protective effect on coronary artery disease (OR = 0.801, p = 0.12)^36^; two independent missense alleles in *PAL2G12A* with strong raising effects on triglycerides, SHBG, testosterone, and strong lowering effects on HDL cholesterol, ApoA, and HbA1c levels (**Supplementary Table 6**); a splice region variant in *CPT1A*, which encodes carnitine palmitoylransferase IA, with lowering effects on triglycerides; and a missense variant in *PCSK6*, with strong ApoB and LDL lowering effects.

For the liver biomarkers, we found a 0.05% MAF inframe deletion in *GOT1* with a 2.4 standard deviation lowering effect on aspartate aminotransferase and a strong increased risk of pancreatitis (p = 8.0 × 10^−6^; OR = 4.55 [2.34, 8.84]); four missense alleles in *GPT*, the alanine aminotransferase 1 gene with strong alanine aminotransferase lowering effects and an increased risk of primary biliary cirrhosis (p = 0.0018, OR = 2.37 [1.38, 4.07]); and two missense alleles in *DGKD*, which encodes diaglycerose kinase delta, an enzyme that phosphorylates diacylglycerol to produce phosphatidic acid, with raising and lowering effects on direct and total bilirubin.

For the renal biomarkers, we found a PTV in *COL4A4* with effects on microalbumin in serum and urine (1.95 and 0.68 sd), and urine albumin to creatinine ratio (2.29 sd), and an estimated increased risk of kidney problems (p = 6.7 × 10^−13^, OR = 6.9 [4.06, 11.60]) and gout (p = 0.0004, OR = 2.3 [1.44, 3.57]) in UK Biobank; a PTV in *SLC22A2* with strong lowering effects on eGFR (0.34 sd) and increasing effect on creatinine (0.50 sd), and an estimated increased risk of kidney problems in UK Biobank (p = 0.00077; OR = 2.77, 95% CI: [1.53, 5.01]); a PTV in *SLC22A11* with raising effects on urate (0.14 sd), and increased risk for gout (p = 0.00015; OR = 1.50 [1.22, 1.85]); a 0.1% rare missense allele in *SLC34A3* with strong eGFR lowering, serum creatinine and Cystatin C raising effect, and increased risk to kidney stones (p = 0.00066; OR = 2.52 [1.48, 4.29])^37^; *SLC47A1 (*also known as MATE1) with multiple independent coding alleles with increasing effect on serum creatinine and increased risk to renal failure requiring dialysis (p = 0.00031; OR = 3.07 [1.67, 5.66]); a missense allele in *SLC6A19, LRP2, ALDOB, SLC7A9*, and *SLC25A45*, all with high expression in kidney with creatinine lowering and eGFR raising effects, among other examples (**Supplementary Table 6**). Together, these results suggest that the genetics of biomarkers will aid in prioritizing disease associated variants.

#### CNVs influencing biomarkers

Copy number variations (CNV) constitute a significant fraction of the genetic differences by affected base pairs between individuals. We found 17 associations from 13 contributing CNVs (Bonferroni p < 5 × 10^−6^, **Supplementary Table 9, Supplementary Figure 10A**). We perform aggregate rare-variant burden tests, pooled by gene for 23,598 genes. We found a total of 29 unique 300kb windows containing genetic associations (Bonferroni p < 2.5 × 10^−6^; **Figure 2B, Supplementary Table 10**) including a burden of rare CNVs overlapping *HNF1B* associate with Urea, eGFR, Creatinine, and Cystatin-C (p < 8.8 × 10^−13^) and are estimated to have large effects on these biomarkers (Beta = 0.77, −0.90, 0.93, 0.98 s.d, respectively; **Supplementary Figure 10B**). *HNF1B* is a membrane bound transcription factor part of the family of hepatocyte nuclear factors, believed to play a role in renal and pancreatic development. Previous studies have associated mutations in *HNF1B* with maturity onset diabetes of the young (MODY) and altered kidney function^38^. Consistent with its developmental role and clinical associations, the rare CNVs overlapping *HNF1B* associate with renal failure in UK Biobank (p = 1 × 10^−7^; OR = 4.94, SE = 0.30; **Supplementary Figure 10B**)^39,40^. These results highlight the value of high-resolution analysis of copy number variation with potentially large effects on lab measurements.

#### Global and local heritability of biomarkers

To characterize the heritability of the 38 biomarkers we first applied LD-score regression^41^. We further applied the Heritability Estimator from Summary Statistics (HESS), an approach for estimating the phenotype variances explained by all typed SNPs at a single locus in the genome while accounting for LD among the SNPs^42,43^. We found that both LD-score regression and HESS find that common SNPs explain a large fraction of the heritability (5.3% [Potassium in urine] to 34.3% [HDL cholesterol] across the studied continuous phenotypes in HESS, **Supplementary Table 11A**). We compare the polygenicity of all 38 biomarkers by computing the fraction of total SNP heritability attributable to loci by the top 1% of SNPs. We found that 10 phenotypes have more than 25% of the SNP heritability explained by the top 1% of genetic associations (Lipoprotein (a) 44%, Total and Direct bilirubin 31 and 30%) and the remaining 28 phenotypes show patterns of high polygenicity (**Supplementary Table 11B, Supplementary Figure 11**).

### Cell type decomposition of genetic effects

We used tissue and cell type expression data to assess whether SNPs within epigenetic annotations of a given tissue/cell type are enriched for heritability^44^. Overall, we found that 20 of the 38 biomarkers are at least 20-fold enriched (p < 1 × 10^−5^) in either kidney, pancreas, or liver, highlighting the primary role of these tissues in the phenotypes we studied (**Figure 3A, Supplementary Figure 12, Supplementary Table 11C**); and observed broadly consistent enrichments in individual annotations that comprised the pancreas, liver, and kidney ChIP-seq experiments (**Supplementary Figure 13**).

**Figure 3.**
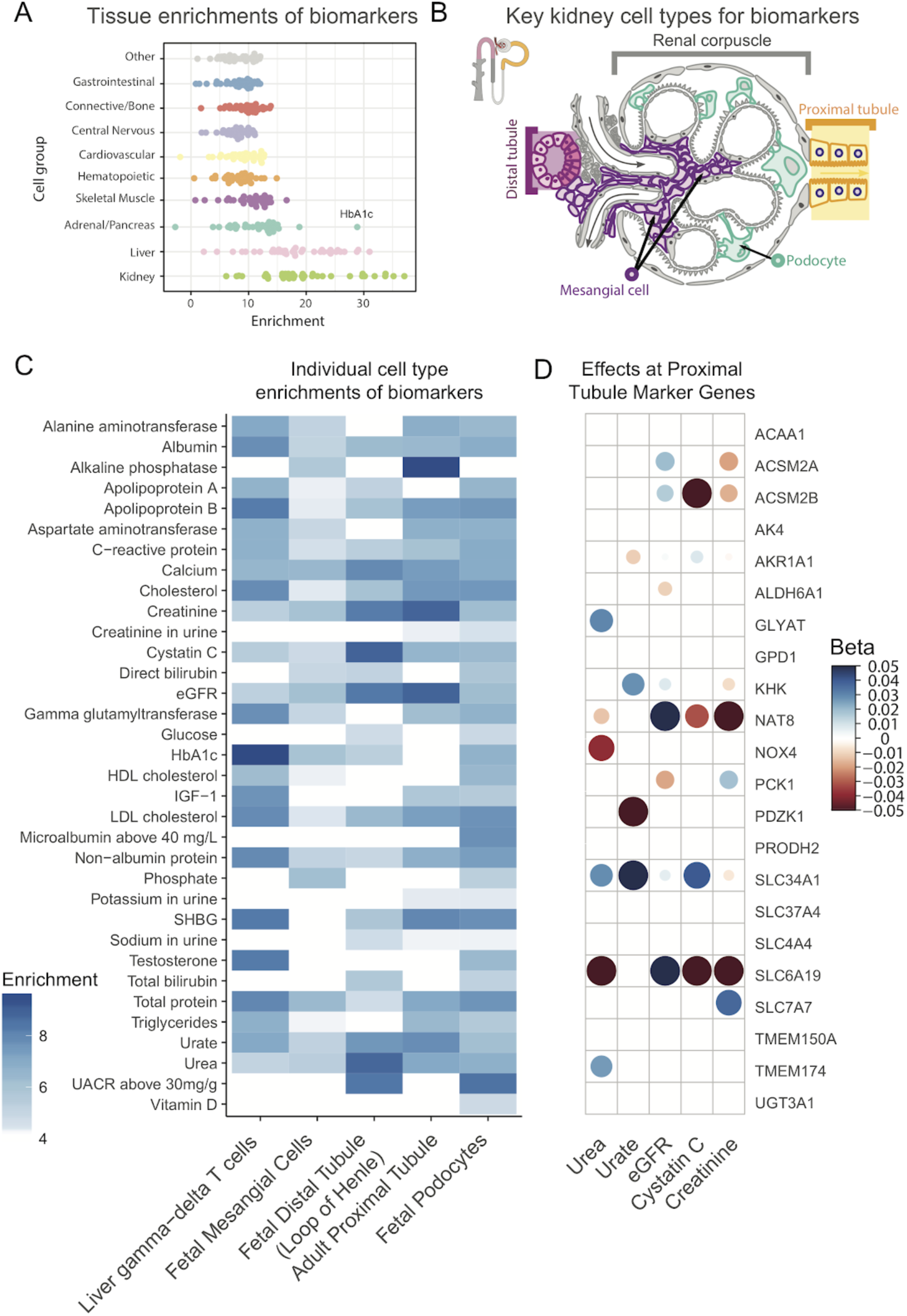
Cell type decomposition of genetic effects. **(A)** (x-axis) Fold enrichment for each biomarker across 10 tissues (y-axis). **(B)** Structure of the renal corpuscle, the blood-filtering component of the nephron of the kidney. **(C)** Single cell heritability enrichment across the 38 biomarkers (y-axis) for single cell types with large numbers of enriched traits in kidney, liver, and pancreas (x-axis). Cells with *P*-value < 10-5 in cell types for which more than 20 biomarkers are enriched are shown. **(D)** Genetic variants (cell coloring and size denotes effect direction and size) associated to renal failure biomarkers (y-axis) in gene markers of single cell gene expression shared between adult mouse kidney proximal tubule and adult and fetal human kidney proximal tubule (x-axis).

We hypothesized that further integration with single cell data may help refine the enrichment across these bulk tissues^45^. As such, we downloaded marker genes of clusters from single-cell RNA-seq of liver (adult human), kidney (adult mouse, fetal human, and adult human), and pancreas (adult human) and ran LD Score regression to test for enrichment of single cell types^46^. Consistent with previous reports^47,48^, we found that numerous traits were significantly enriched in podocytes, proximal and distal tubule, and hepatocytes, with our large sample sizes capturing a novel enrichment of variation near marker genes of mesangial cells across many traits and γδ T cells in HbA1c, possibly due to their role in inflammation during diet induced obesity (**Figure 3B-D, Supplementary Table 11D**)^49^.

### Targeted phenome-wide association study

We performed a phenome-wide association analysis (PheWAS) to detect whether the variants we implicated might impact other diseases or clinically relevant phenotypes (**Supplementary Table 12, Supplementary Figures 14,15**). We find a total of 218 associations across 86 phenotypes for 25 protein-truncating and 80 LD-independent protein-altering variants that were also associated with increased risk for 68 disease outcomes, and lower risk of 39 disease outcomes (p < 1 × 10^−5^). Overall, these results demonstrate that variants with effects on biomarkers have pleiotropic effects across diverse phenotypes.

### Correlation of genetic effects between biomarkers, diseases, and medically relevant phenotypes

Given the widespread polygenicity and pleiotropy observed in the GWAS and PheWAS analysis, we then estimated global genetic correlation patterns between the biomarkers, diseases, and medically relevant phenotypes. We applied LD-score regression to estimate genetic correlations^50^ among the 38 biomarkers and an additional 146 summary statistics including 56 diseases and 90 previously published medically relevant phenotypes (**Figure 2C, 4A, Supplementary Figures 16-18, Supplementary Table 13**). Overall, we found that between the 38 biomarkers and the 146 other phenotypes, there exist 1,127 significant non-zero correlations of genetic effects (**Figure 4A**, p < 1 × 10^−3^).

**Figure 4.**
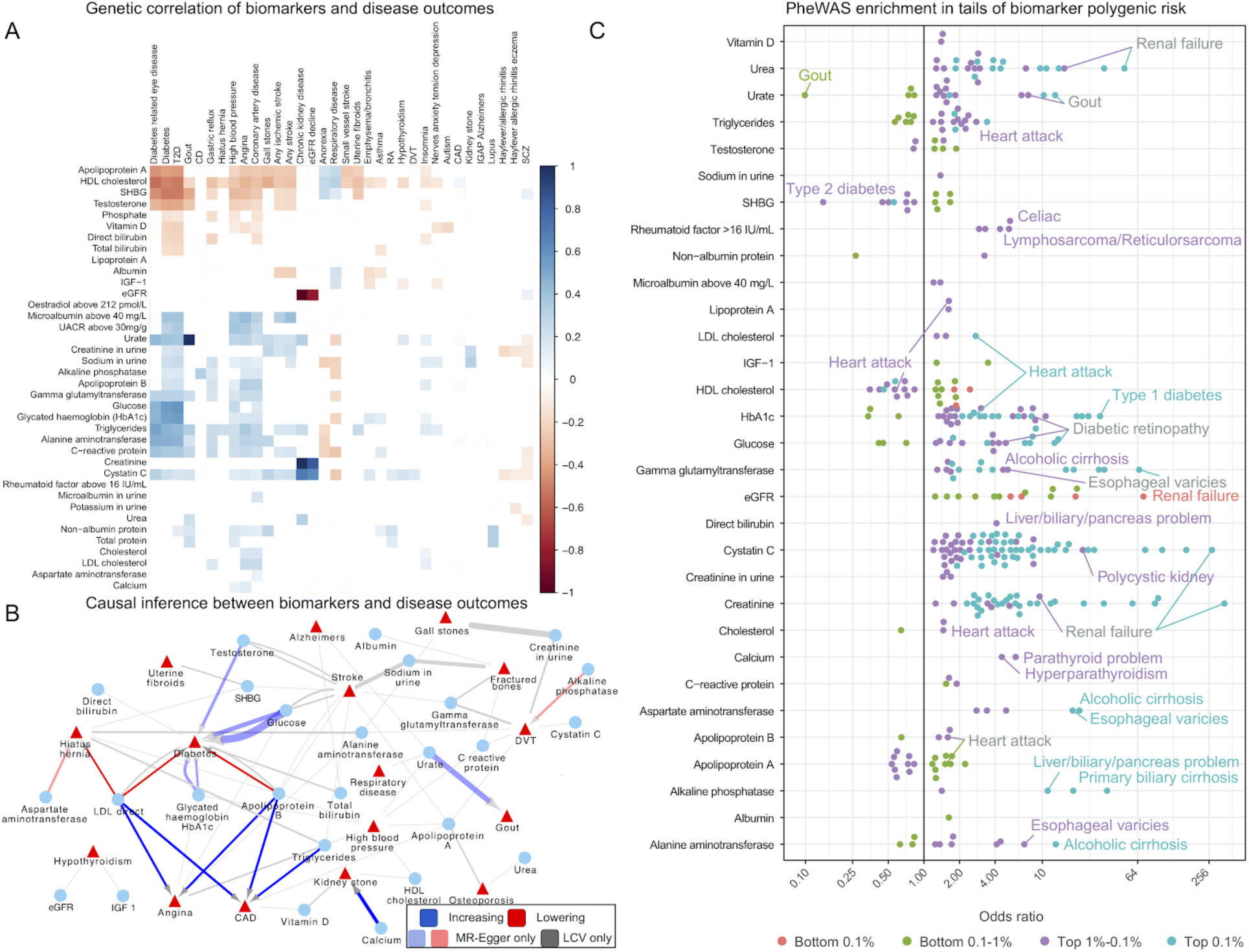
Correlation of genetic effects, causal inference, and complex trait association in polygenic risk tails. **(A)** Correlation of genetic effects plot between the 38 biomarkers and 123 complex traits using LD-score regression. Cells with p < 0.001 are highlighted, and traits (n=26) with no associations are not shown. **(B)** MR-Egger and LCV predict causal links between biomarkers (blue nodes) and selected complex traits (red nodes). Association arrows are drawn based on MR-Egger (red, blue), LCV (gray), or both (dark), and multiple arrows indicate support from multiple studies. MR-Egger and LCV were jointly adjusted for FDR 10% cutoff across all tests. Triangles are used for binary and circles for continuous summary statistics. Edge width is proportional to the absolute causal effect size, estimated by MR Egger. A complete listing of discovered associations is provided as a table (Supplementary Table 14). **(C)** (x-axis) Biomarker polygenic risk scores for the top 0.1%, top 1%, bottom 0.1%, and bottom 1% of individuals and their association to different diseases in UK Biobank, represented as the odds ratio of the disease in this group relative to the middle 40-60% of individuals.

### Causal inference

The patterns of significant correlation of genetic effects between the 38 biomarkers and 123 diseases and medically relevant phenotypes raised the possibility that some of these associations might be causally relevant.

First, to estimate causal effects we used *MR-Egger* to perform Mendelian Randomization, treating the genome-wide significant variants for each trait as instrumental variables^51^ (**Methods**). Using MR-Egger we find 86 causal relationships at an FDR of 10%, many of which are causal relationships to disease outcomes (**Supplementary Table 14**).

It has been noted that Mendelian Randomization can be confounded by genetic correlations reflecting shared etiology, and sometimes instruments with pleiotropic effects may introduce limitations in evaluating causal relationships. Hence, to distinguish between genetic correlation from causation we used the O’Connor and Price Latent Causal Variable (LCV) Model^52^. We found that 49 of the 86 causal relationships inferred by Mendelian Randomization are recovered by LCV (**Figure 4B**). 344 additional causal relationships are unique to LCV, highlighting potentially novel causal associations (**Supplementary Table 14**). Many of these are well-described -- such as that of LDL cholesterol on coronary artery disease and angina, which we estimate using MR-Egger at 0.34 log odds change per standard deviation (LCV causal percent 0.75) and 0.313 LOR/SD (LCV genetic causal percent [GCP] 0.8) respectively, or the effect of urate on gout (MR-Egger slope 1.2 LOR/SD, LCV n.s.). Our large sample size provide unique genetic evidence supporting the effect of calcium levels on kidney stones (0.625 LOR/SD and GCP 0.48), and testosterone on diabetes (0.49 LOR/SD, LCV n.s.) consistent with existing epidemiological reports^53,54^. We also discovered novel associations, such as that of AST levels on hernia (−0.319 LOR/SD, GCP 0.14), that warrant further study.

### Polygenic prediction of biomarkers within and across populations

The vast size of our cohort affords the opportunity to build predictive polygenic risk models of biomarkers from genotype data alone^55^. We constructed PRS for all 38 biomarkers using batch screening iterative lasso (BASIL) implemented in R snpnet package^56,57^(**Methods**). To do this, we split the self-identified White British sample set into 60% training, 20% validation, and 20% test sets, and evaluated the trained models using goodness-of-fit (*R*) for the quantitative phenotypes or the area under the receiver operator curve (AUC) for the binary phenotypes (**Supplementary Figure 19A, Supplementary Table 15**). The inclusion of the lasso coefficients for all these phenotypes is shown in the form of “lake” plots showing shrinkage and variable selection (**Supplementary Figure 20**). We observed strong concordance between predicted and observed phenotypes (**Supplementary Figure 19 B-D**). We found that predicted Lipoprotein (a) level stratified individuals into two groups, which reflects the huge contribution of the *LPA* gene (**Supplementary Figure 19B**). To assess whether this prediction performance could translate to other populations, we compared predicted genetic scores for the 38 biomarkers to their true measured or derived values in 24,131 self-identified Non-British White, 6,951 South Asian, 1,950 East Asian, and 6,056 African individuals in the UK Biobank. Overall, these 4 populations experienced median reductions in prediction performance of 6.4%, 30.0%, 43.2%, and 72.0% respectively, suggesting these polygenic models have somewhat limited generalizability across populations (**Supplementary Figure 19A, E, Supplementary Tables 15,16**)^58^.

### Multiple regression with PRSs for biomarkers improves prediction of traits and diseases

We then tested the hypothesis that the 38 biomarker PRSs may improve the prediction of higher-level traits and diseases in combination with the PRS for the trait or disease itself. To this end, we constructed multi-PRS models for renal failure, alcoholic cirrhosis, myocardial infarction, kidney cancer, liver cancer, and liver fat percentage by using multiple regression to predict the trait or disease from a) its own PRS, b) the PRSs for each of the 38 biomarkers, and c) relevant covariates such as age and sex **(Methods)**.

We selected these multi-PRSs by considering the enrichment of disease prevalence in the UK Biobank at the tails of the distribution of predicted values (**Figure 4C**). We calculated the top and bottom 0.1% and 0.1-1% bins of self-identified-White British individuals by polygenic score and compared them to the 40-60% center of the distribution with a Fisher exact test. Traits with low case numbers and multiple enriched biomarkers, which we reasoned would benefit most from the combination of multiple biomarker PRSs, were selected for further analysis.

We further tested multiple disease outcomes and liver fat percentage (LFP), a quantitative measure derived from costly MRI images of the liver obtained approximately 8 years after the initial assessment visit where the serum and urine samples used for our polygenic score calculation were collected^59^. Liver fat is driven by a combination of alcohol use and metabolic disorder^60^. Only 4,617 individuals thus far have quantified LFP in UK Biobank, and we reasoned that the substantial power increase from the full cohort might help with prediction of this important physiological parameter.

First, we ran ordinary least squares, predicting LFP from covariates (including alcohol and interactions; see **Methods**). Covariates were moderately effective at predicting liver fat percentage (adjusted r-squared 0.024, **Supplementary Table 17**), and adding our snpnet-derived polygenic risk score increased the predictive power substantially (**Figure 5A, Supplementary Tables 17,18, Supplementary Figure 21a**, adjusted r-squared 0.050, F test p < 1 × 10^−10^). Adding the 38 biomarker PRSs to the regression improved predictive capacity further (0.087, F test vs LFP PRS alone p < 1 × 10^−10^, **Supplementary Tables 17,18**). The PRS for Alanine aminotransferase, sodium in urine, urate, SHBG, and triglycerides all had significant coefficients (**Supplementary Tables 19**), supporting the previously described notion that complex interplay between organ systems, or multiple underlying disease states, might contribute to LFP^61^. Meanwhile, the myocardial infarction snpnet PRS was equally stratifying to the multi-PRS, with both explaining a substantial portion of trait heritability (R^2^ = 0.15-0.16; Figure 5B). This suggests that the genetic basis of MI, as previously reported, is largely driven by lipids (**Supplementary Figure 21b, Supplementary Table 21**). In contrast, for renal failure, the multi-PRS resulted in the highest risk individuals having a nearly 30% prevalence (compared to 5% with the snpnet PRS alone, **Figure 5B, Supplementary Table 21**). This effect was replicated, albeit with a smaller effect, in self-identified non-British White individuals (**Supplementary Figure 22, Supplementary Table 21**).

**Figure 5.**
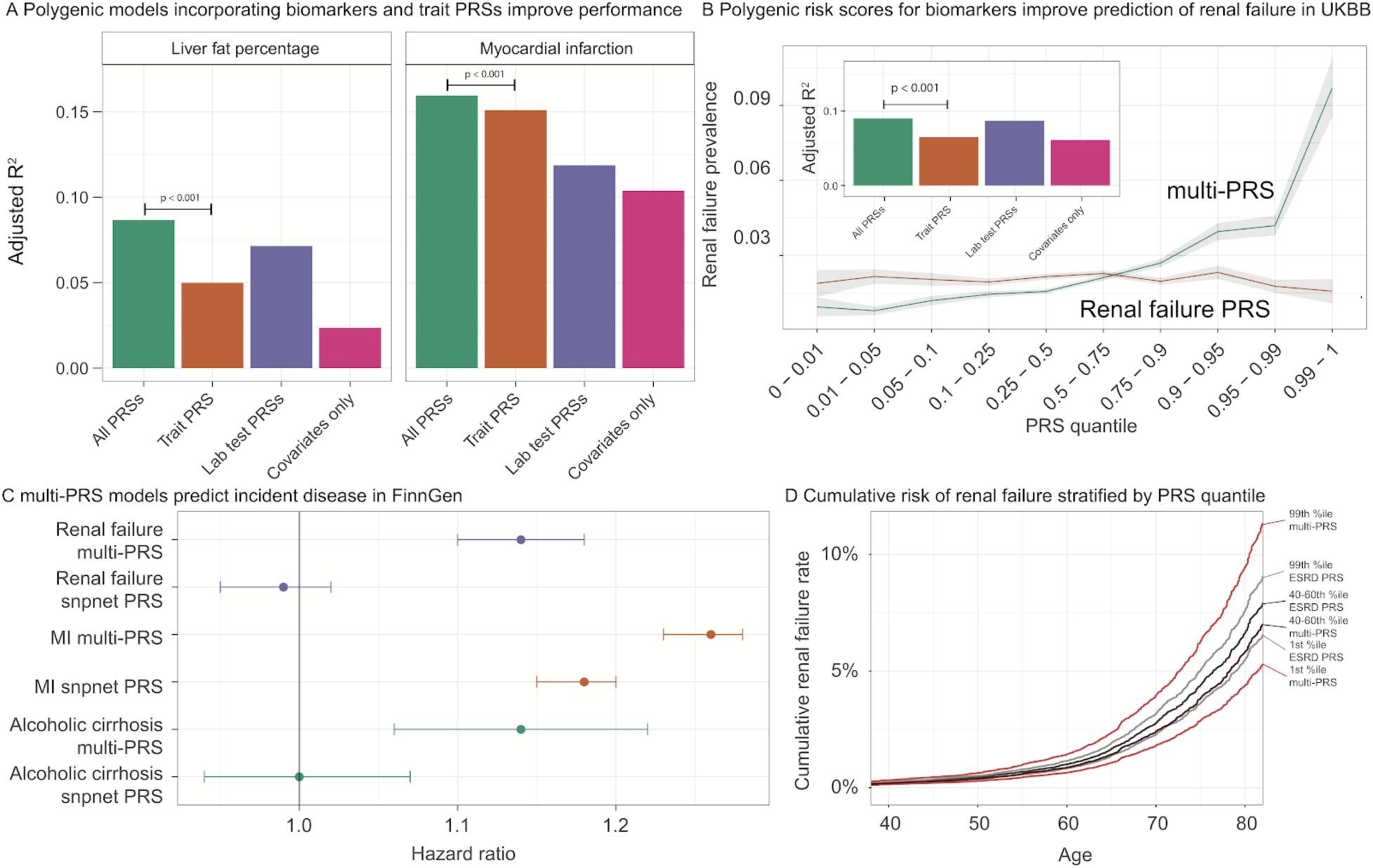
Multi-Polygenic Risk Scores improve disease incidence and prevalence prediction. **(A)** Improvement in prediction accuracy of traits when including biomarker polygenic risk scores. Each trait was tested against a model with just the covariates (for liver fat, including alcohol and interactions; for all traits, including age, sex, Townsend Deprivation Index, principal components, and interactions); with the covariates and the polygenic score for the trait (trained using snpnet), termed “Trait PRS”; with the covariates and all the polygenic scores for biomarkers, termed “Biomarker PRSs”; and with covariates and all PRSs, termed “All PRSs.” In each case, the trait PRS only was outperformed by including all the biomarker PRSs as well. See F-test results and regression terms in Supplementary Tables 18-21. (**B)** (x axis) quantiles of polygenic risk score, spaced to linearly represent the mean of the corresponding bin of scores. (y axis) Prevalence of renal failure (defined by verbal questionnaire and hospital in-patient record ICD code data) within each quantile bin of the polygenic risk score. Error bars represent the standard error around each measurement. **(C)** Hazard ratios for incidence of renal failure (n=3,058), myocardial infarction (n=7,913), and cirrhosis (n=845) in FinnGen using the standard PRS trained on UK Biobank using snpnet versus the multi-PRS including both biomarker PRSs and the trait PRS. Error bars represent 95% confidence intervals. **(D)** Cumulative renal failure rate in quantiles of Finnish individuals in FinnGen. Individual estimates were predicted from existing polygenic scores and the bins of individuals in both the snpnet PRS for renal failure and the multi-PRS are shown. The central dark color corresponds to the 40-60th percentile of renal failure (grey; ‘ESRD PRS’) or multi-PRS (red, ‘multi-PRS’) bins, and the lighter color corresponds to the top and bottom 1 percentiles.

Encouraged by these findings, we evaluated the potential of these improved polygenic scores in cases of incident disease. We examined myocardial infarction (MI), renal failure, alcoholic cirrhosis, kidney and liver cancer, all of which had improved predictive power in the presence of the biomarker PRSs (**Figure 5A, Supplementary Tables 20,21**), but which varied in total predictive power and in improvement from inclusion of the biomarker PRSs. To assess whether the improvement in predictive performance was transferable, we applied both trait specific PRS and combined PRS in FinnGen (n = 135,500, **Supplementary Tables 22**). Here, we found strong evidence that the combination of PRS increased the effect size in myocardial infarction (hazard ratio = 1.18 per SD increment for snpnet PRS and 1.26 for multi-PRS), renal failure (hazard ratio = 0.99, p = 0.46 for snpnet PRS and hazard ratio = 1.14, p = 9.3 × 10^−13^ for multi-PRS, **Figure 5C,D, Supplementary Tables 23,24**), and alcoholic cirrhosis (hazard ratio = 1.00, p = 0.98 for snpnet PRS and hazard ratio = 1.14, p = 1.8 × 10^−4^ for multi-PRS, **Supplementary Tables 23,24**) and increased lifetime prevalence and cumulative disease rates over time at the tails of polygenic risk (**Supplementary Figures 23,24**). This suggests that multiple regression of polygenic risk for biomarkers might capture multiple underlying disease states and/or underlying causes, similar to what has recently been reported for neuropsychiatric disease^62^, and that these multiple states are predictive of incident disease.

## Discussion

Using data from 38 biomarkers in ∼360,000 UK samples, we provide a systematic assessment of genetic associations, as well as their disease relevance and predictive performance.

Detailed analysis of HLA alleles, and copy number, protein-altering, and protein-truncating variants highlight potential drug targets. In contrast, in non-coding variants, we found strong enrichment of polygenic heritability in specific single-cell clusters from kidney and liver tissue, including podocytes, stellate cells, and the descending loop of Henle. These results are corroborated by mouse knockout data^63^. By expanding single-cell gene expression and epigenetic resources, particularly under stress and disease conditions, we will likely reveal additional novel cellular enrichment signatures to inform the mechanism underlying our novel large-effect variants.

The genome-wide resource made available with this study provides a starting point for cataloging variants affecting the 38 biomarkers, and larger datasets across under-represented populations may improve their generalizability. These results highlight the benefits of direct measurements of biomarkers for interpretation of genetic variation.

To assess the translatability of our findings, we built predictive models aggregating trait PRS with those of the biomarkers, improving the predictive accuracy of multiple disease outcomes both overall and especially at the extremes of genetic risk. Methodologies such as this will improve resolution in dissecting personalized drivers of disease. Integration with independent population biobanks and prospective cohorts will help elucidate the extent to which these combined risk models can be transferred. Together, we anticipate this will inform clinical practice in the coming years as a larger fraction of the population is genotyped and sequenced.

## Methods

### Genotype and phenotype data

We used genotype data from the UK Biobank dataset release version 2 and the hg19 human genome reference for all analyses in the study^64^. To minimize the variability due to population structure in our dataset, we restricted our analyses to unrelated individuals based on the following four criteria reported by the UK Biobank in the file “ukb_sqc_v2.txt”:

1. used to compute principal components (“used_in_pca_calculation” column)
2. not marked as outliers for heterozygosity and missing rates (“het_missing_outliers” column)
3. do not show putative sex chromosome aneuploidy (“putative_sex_chromosome_aneuploidy” column)
4. have at most 10 putative third-degree relatives (“excess_relatives” column).

We used a combination of self reported ethnicity and principal component analysis (UK Biobank field ID: 21000) and analyzed 5 subpopulations in the study : self-identified White British (n = 337,151 individuals), African (6,497), East Asian (2,061), South Asian (7,363), and self-identified non-British White (26,471). Using the imputed common variants on chromosome 1 (MAF >=10%, 443,757 variants remained), we applied principal component analysis (PCA) implemented in PLINK v2.00aLM (17 Jul 2017) with --pca approx 10 and characterized the top 10 components. We defined threshold on PC2 as shown below and defined European, African, East Asian, and South Asian individuals (**Supplementary Figure 1**).

- PC2 <= −0.011 indicates East Asian ancestry
- PC2 in range [−0.007, −0.004] indicates South Asian ancestry
- PC2 in range [−0.002,0.002] indicates European ancestry
- PC2 >= 0.003 indicates African ancestry

These cutoffs were selected by overlaying self-reported ancestry from UK Biobank field 21000, which revealed distinct ancestry clusters. Individuals between the boundary regions (i.e. no confident ancestry grouping) were excluded from the analysis. For European individuals, we identified self-reported White British individuals and self-identified non-British White Europeans. We subsequently focused on a subset of individuals with non-missing values for covariates and biomarkers as described below.

We annotated variants using the VEP LOFTEE plugin (https://github.com/konradjk/loftee) and variant quality control by comparing allele frequencies in the UK Biobank and gnomAD (gnomad.exomes.r2.0.1.sites.vcf.gz) as previously described^28^. We focused on variants outside of major histocompatibility complex (MHC) region (chr6:25477797-36448354) and performed LD-pruning using PLINK with “--indep 50 5 2” as previously described^28,65^. The LD-pruned sets are used for targeted PheWAS analysis described below.

We focused on 32 biomarkers (UKBB field ID column in **Supplementary Table 2**) and also defined two derived phenotypes, estimated glomerular filtration rate (eGFR) and non-albumin proteins. The eGFR measure is an indicator of renal function and is defined by the CKD-EPI equation^66^. We defined non-albumin protein as the difference between the total protein and albumin. Given the dominance of signals below the detection limit for some biomarkers, we additionally defined four binary phenotypes:

- Urine albumin to creatinine ratio >30 mg/g
- Microalbumin >40 mg/L
- Rheumatoid factor >16 IU/mL
- Oestradiol >212 pmol/L

We treated individuals beyond the detection limit for those biomarkers as cases in those four binary phenotypes, and below the detection limit as controls, as reported by the corresponding reportability fields.

#### Statin identification and LDL adjustment

We reviewed the medications taken by one or more participants in the UK Biobank and identified 13 medication codes corresponding to statins (1141146234, atorvastatin; 1141192414, crestor 10mg tablet; 1140910632, eptastatin; 1140888594, fluvastatin; 1140864592, lescol 20mg capsule; 1141146138, lipitor 10mg tablet; 1140861970, lipostat 10mg tablet; 1140888648, pravastatin; 1141192410, rosuvastatin; 1141188146, simvador 10mg tablet; 1140861958, simvastatin; 1140881748, zocor 10mg tablet; 1141200040, zocor heart-pro 10mg tablet). We then identified participants (n ∼ 1,427) with biomarker measurements who were not taking a statin upon enrollment (years 2006-2010), but who were taking a statin at the time of the first repeat assessment visit (years 2012-2013). For each participant, we divided their on-statin biomarker measurement by their pre-statin biomarker measurement. The mean of this value was considered to be the statin correction factor within the UK Biobank. For all individuals who were taking statins upon enrollment, we divided their on-statin measurement by the correction factor to yield an adjusted biomarker measurement value. For all traits, we calculated a p-value from a wilcoxon signed rank test for paired samples comparing whether the pre- and on-statin values were significantly different, and only traits with a significant non-zero effect were evaluated with adjustment for statins. The following list of statins were identified in the UK Biobank for the purposes of adjusting by the estimated factor: 1140861958, simvastatin; 1140888594, fluvastatin; 1140888648, pravastatin; 1141146234, atorvastatin; 1141192410, rosuvastatin; 1140861922, lipid-lowering drug; 1141146138, lipitor 10mg tablet.

#### Covariate correction

Raw UK Biobank measurements for all reported individuals (excluding out of range and QC failed measurements) were fit with linear regression against the 127 covariates. These included demographics (age, sex, age*sex, age^2^), population structure (the top 40 principal components and indicators for each of the assessment centers in the UK Biobank), temporal variation (indicators for each month of participation, with the exception that all of 2006 and August through October of 2010 were considered single months), socioeconomic status indicators (townsend deprivation indices and interactions with age and sex), the genotyping array used, and technical confounders (blood draw time and it’s square and interactions with age and sex; urine sample time and its square and interactions with age and sex; sample dilution factor; fasting time, its square, and interactions with age and sex; and interactions of blood draw time and urine sample time with dilution factor). The residual from this regression was inverse normal transformed using the Blom transform and then used as the tested outcome. For sensitivity analysis, we also applied covariate transformation within just the self-identified White British individuals and obtained similar estimates.

### Genome-wide association analysis

We performed association analyses using imputed 1000 Genomes Phase I variants (for non-coding variants), directly genotyped variants on array (for protein-truncating and protein-altering variants), imputed HLA alleles, and copy number variations (CNVs).

#### GWAS of imputed 1000 Genomes Phase 1 variants

We employed a GWAS without covariates of the residuals computed above. This was run using plink v2.00aLM with the following parameters:

> --glm cols=chrom,pos,ref,alt,altfreq,firth,test,nobs,orbeta,se,ci,t,p hide-covar --pgen <imputed PGEN> --remove <non-White-British individuals> --keep <all individuals, males, or females> --geno 0.1 --hwe 1e-50 midp;

For binary traits, a logistic glm was fit directly with covariates to age, sex, genotyping platform, 10 PCs, age^2^, and fasting time, with the --vif 999 parameter to avoid the collinearity of age and age^2^.

#### Derivation of independent loci

Once we ran the GWAS, full summary statistics were clumped to r^2^ > 0.1 using the following clump command:

> plink1.9 --bfile <1000G Phase 3 European plink file> --clump <summary statistics> --clump-p1 1e-6 --clump-p2 1e-4 --clump-r2 0.1 --clump-kb 10000 --clump-field P --clump-snp-field ID

Then, to avoid calling very large signals as multiple associations, these were further filtered such that any SNPs within 0.1cM of each other (as annotated by 1000 Genomes) were considered part of the same association signal, with the cM annotation derived from the 1000G Phase 3 European samples (n = 489)^41^ -- variants within 0.1cM were chose to only have the minimum p-value. For the final results, all lead variants with p < 5 × 10^−8^ were kept for the mendelian randomization analyses.

In order to report independent signals, we ran the following plink command:

> plink1.9 --bfile <1000G Phase 3 European plink file> --extract <all unique hit SNPs, n = 6269> --indep 50 5 2

And counted the number of independent SNPs it reported.

#### GWAS on coding variants on genotyping array

Univariate association analyses for single variants were applied to the 38 phenotypes independently using PLINK v2.00aLM (2 April 2019). For binary phenotypes, we performed Firth-fallback logistic regression as previously described^28^. For the residuals of the quantitative phenotypes after adjusting the 127 covariates, we applied generalized linear model association analysis.

#### Cascade plot visualization of coding and non-coding variants

We visualized the minor allele frequency and effect size estimates in a series of cascade plots. For protein-truncating and protein-altering variants, we focused on genome-wide significant associations with p < 1 × 10^−7^ and annotated the corresponding gene symbols for variants with absolute value of betas greater than 0.1 (outliers). For non-coding variant associations characterized on the imputed 1000 Genomes Phase 1 variants, we focused on the clumped set of associations (described above in “Derivation of independent loci” section above) and applied the following procedure to determine and highlight the outliers:

- Fit a univariate linear regression model with absolute value of effect size estimate (BETA or log odds ratio) as the response and the log of minor allele frequency as the predictor.
- Find the residuals from the regression model and find the mean and standard deviation of the residuals.
- We defined association is an outlier on cascade plot if and only if the residuals from the regression model above is outside of the mean plus or minus 1 SD range.

#### Association and Bayesian model averaging analyses for HLA alleles

The HLA data from the UK Biobank contains all HLA loci (one line per person) in a specific order (A, B, C, DRB5, DRB4, DRB3, DRB1, DQB1, DQA1, DPB1, DPA1). We downloaded these values, which were imputed via the HLA:IMP*2 program (Resource 182); the UK Biobank reports one value per imputed allele, and only the best-guess alleles are reported. Out of the 362 alleles reported in UKB, we used 175 alleles that were present in >0.1% of the population surveyed.

We performed association analysis for our 38 phenotypes and 175 HLA alleles using PLINK v2.00aLM (2 April 2019). We included only self-identified White British individuals (n = 337,151). We used generalized linear models for quantitative traits and generalized linear models with a Firth-fallback method for binary traits.

To identify the HLA alleles that were not simply associated to a particular phenotype due to LD, we used Bayesian Model Averaging (BMA), implemented in the ‘bma’ R package^67^. BMA is a model selection method that trains a variety of models, one on each possible subset of alleles. The posterior probability of each model being the correct one given the data is determined, and subsequently, a BIC per model is calculated. The degree to which an allele is included across models (posterior probability) is then deemed a measure of confidence in the association between allele and phenotype.

We first filtered the allele dosage file to those columns that were not sparse, making sure that each allele in the analysis had more than 5 entries. We then identified all of the allele-phenotype pairs that had BY-adjusted p-value <0.05 from PLINK. If there were >10 alleles below this threshold for a given phenotype, we used the 10 alleles with the lowest adjusted p-value in order to maintain computational tractability. If there were >2 such alleles for a given phenotype, we did not run BMA for that phenotype. These requirements filtered our testing base down to 33 phenotypes, with 56 alleles included in at least one analysis.

In order to maintain computational tractability, only models whose posterior model probability was within a factor of 1/5 of that of the best model were kept for the final analysis. We focused on alleles with posterior probabilities > 0.8 based on our BMA analysis. We ran BMA with a binomial error distribution and link function for binary traits and Gaussian ones for quantitative traits.

We used the in-built ‘imageplot.bma’ function to produce the plots of the model architectures, and report allele, phenotype, posterior mean effect size, standard deviation of said effect size, and the posterior probability that the effect is not equal to 0.

#### Copy number variants

CNVs were called by applying PennCNV v1.0.4 on raw signal intensity data from each array within each genotyping batch as previously described^68^, with the notable difference that here, all analyses are conducted within the self-identified-White-British unrelated cohort described above. Data for phenome-wide associations were derived from UK Biobank data fields corresponding to anthropometric measurements, laboratory tests, disease diagnoses, and medical procedures from medical records, as well as a questionnaire about lifestyle and medical history. We compute generalized linear models using the PLINK v2.00aLM (31 Mar 2018) --glm option. For burden tests, we add number and total length of CNV as covariates for both binary and quantitative traits. See the “GWAS on genetic variants on genotyping array” section for further description of PLINK’s implementation of these model specifications.

### Heritability estimates

#### LD Score regression

We used the default LD Scores from the 489 unrelated European individuals in 1000 Genomes as our reference. We converted our summary statistics to LDSC format using munge_sumstats, munging against the set of 1000 Genomes Phase 1 variants with calls of an ancestral allele in 1000 Genomes Phase 3. We ran ldsc.py with the following parameters:

> ldsc.py --h2 <trait summary statistics> --ref-ld-chr <ldsc/1000G.EUR.QC/> --w-ld-chr <ldsc/weights_hm3_no_hla/weights.>

#### HESS

We performed standard stage 1 fitting^42^, then removed all regions which contained no SNPs with MAF > 5% (5/∼1700 bins genome wide) and generated stage 2 estimates from the resulting matrices. We used the same munged sumstats described above, which were generated using a modified version of the munge_sumstats.py which also outputs chromosome and position. We confirmed heritability estimates of select associations using GCTA-GREML and genotyped array variants on a subset of individuals (data not shown) to ensure estimates were comparable to this model.

### Cell-type enrichment analysis

We ran stratified LD Score regression with the 53 baseline annotations and included all 10 tissue type annotations and the Roadmap control regions. The exact command was:

> ldsc.py --h2 <trait sumstats> --ref-ld-chr
>
> ldsc/1000G_EUR_Phase3_baseline/baseline.,ldsc/1000G_Phase3_cell_type_groups/cell_type_group.1.,ldsc/1000G_Phase3_cell_type_groups/cell_type_group.2.,ldsc/1000G_Phase3_cell_ty pe_groups/cell_type_group.3.,ldsc/1000G_Phase3_cell_type_groups/cell_type_group.4.,ldsc/10 00G_Phase3_cell_type_groups/cell_type_group.5.,ldsc/1000G_Phase3_cell_type_groups/c ell_type_group.6.,ldsc/1000G_Phase3_cell_type_groups/cell_type_group.7.,ldsc/1000G_Phase 3_cell_type_groups/cell_type_group.8.,ldsc/1000G_Phase3_cell_type_groups/cell_type_group.9.,ldsc/1000G_Phase3_cell_type_groups/cell_type_group.10.,ldsc /ldscores/Roadmap/Roadmap.control. --w-ld-chr ldsc/weights_hm3_no_hla/weights. --overlap-annot --frqfile-chr ldsc/1000G_frq/1000G.mac5eur.

We also ran each of the 394 Roadmap experiment annotations independently, including just the Roadmap control and baseline annotations as covariates^69^.

### Single-cell enrichment analysis

We ran stratified LD Score regression with the 53 baseline annotations and included all 10 tissue type annotations and the Roadmap control regions. To this, we extended the RefSeq gene region of each marker gene by 100kb and used these as a separate annotation, which we tested for enrichment. This was done separately for each of the single cell clusters for each experiment.

### Targeted Phenome-wide association analysis

We prioritized the following sets of variants for targeted phenome-wide association analysis.

1. PTVs with at least one significant associations (p < 1 × 10^−7^) with the 38 biomarkers
2. Protein-altering variants with at least one significant associations (p < 1 × 10^−7^) with the 38 biomarkers

For each set of variants, we used Global Biobank Engine (GBE) to query significant associations (p < 1 × 10^−5^) across previously reported binary phenotypes^28,65^. We visualized association results for both 38 biomarkers and other GBE traits as heatmaps and sorted variants and phenotypes with hierarchical clustering.

### Correlation of genetic effects across relevant phenotypes

We used LD Score regression in genetic correlation mode to estimate genetic correlation effects between biomarkers and other traits. The exact arguments were:

> ldsc.py --rg <traits> --ref-ld-chr ldsc/1000G.EUR.QC/ --w-ld-chr
>
> ldsc/weights_hm3_no_hla/weights.

### Causal inference

We ran LCV with a number of LDSC-formatted summary statistics files with the default parameters, using a customized script which takes in a list of summary statistics and applies LCV between the first and all others sequentially.

For MR, we used TwoSampleMR to calculate MR Egger regressions and perform trait munging^70^. To scale betas from the Neale lab binary trait outcomes, we considered the prevalence of the trait and raw mendelian randomization beta (with units of change of outcome per standard deviation change in the biomarker), and calculated the log odds ratio as log((prev + beta) / prev). In our simulations (data not shown), this is approximately equivalent to logistic regression across a range of prevalences and betas.

All MR and LCV results were jointly adjusted for multiple comparisons using the standard BH FDR algorithm (at 10%). Network visualization of the results was done using Cytoscape^71,72^. Scripts are provided on our github repository https://github.com/rivas-lab/public-resources/tree/master/uk_biobank/biomarkers.

### Polygenic prediction within and across populations

We applied the batch screening iterative lasso (BASIL) algorithm implemented in R snpnet package to find the exact lasso solution for ultra-high dimensional large dataset through an iterative procedure built on top of the R glmnet package^56,57,73^. For each trait, we randomly split self-identified-White-British individuals with non-missing values into 60% training, 20% validation, and 20% test sets and fit a multivariate Lasso regression model. With the 127 covariates described above, we used training and validation sets for training using R-snpnet package with the default parameters for phenotypes after the statin adjustments. For microalbumin in urine, we used a simpler model with a limited set of covariates -- age, sex, and the first 10 PCs -- given the smaller number of individuals. For two traits, Lipoprotein A and total bilirubin, we noticed that the snpnet package did not find the optimal lambda within the default maximum number of iterations (100). We took the model from the 100th iteration as the best model among the tested during the training phase.

Using the beta values for array-genotyped SNPs and covariates from multivariate Lasso regression, we computed polygenic risk score for each individual with PLINK2 --score subcommand and evaluated the goodness of fit using the test set. Specifically, we computed correlation coefficient R for continuous traits or ROC-AUC metric for binary traits for risk scores quantified from both genotype and covariates and covariates only, and quantified the difference of those two as the increment of predictive performance.

We applied the same evaluation procedure for the four non-White-British populations in the UK Biobank: self-identified non-British White, East Asian, South Asian, and African. We evaluated the transferability of our polygenic risk score within and across ethnic cohorts by comparing the increments of predictive performance between self-identified-White-British and the other four populations.

### Individual Extreme PRS-PheWAS

We started by enumerating all our high-confidence traits which were replicated between ICD codes and self reporting (n = 366), cancer outcomes (n = 46), family history traits (n = 10), and manually curated traits (n = 49)^28^. For each of the 38 biomarkers, we used R’s fisher.test implementation of the Fisher’s Exact test between the 40-60 percentile and the top and bottom 0.1% and 0.1-1%. We then corrected for multiple hypotheses using a Bonferroni-adjusted q-value less than 5% within each biomarker and report the enrichment as the odds ratio estimate from the Fisher’s exact test.

### Models for multi-PRS prediction of disease outcomes

We began by calculating polygenic scores from the snpnet predictions across all individuals in the dataset. For each outcome trait independently, we ran a covariate-only logistic regression with age, sex, 40 principal components of the genotyping matrix, genotyping array, age^2^, an age * sex interaction, and townsend deprivation index (and, for liver fat percentage, current alcohol consumption status at the second time-point and interactions of this with age and sex). Covariate-only models were augmented with the polygenic score for the corresponding trait (“Trait only”), for the polygenic score for the 38 biomarkers (“Biomarkers only”), or with both (“All PRSs”). We used R’s glm implementation and the McFadden’s adjusted PseudoR2 from DescTools (binary outcomes) or R’s lm implementation and reported adjusted r^2^ (continuous outcomes), along with relevant F tests with the anova command, to evaluate prediction.

In order to perform out-of-sample validation, we trained L1-regularized logistic or linear regression models with glmnet using just the 38 biomarker PRSs and the PRS for the trait of interest in the validation subset from the trait of interest. Results were evaluated in the test subset of the trait of interest and in all unrelated, self-identified non-British White individuals for which the corresponding phenotype was available (as used in the cross-population testing; see above).

We predicted the trait of interest in these test sets, calculated the quantiles of predicted trait, and averaged the phenotype within each non-overlapping quantile.

### Evaluation of multi-PRS prediction in an external cohort

The FinnGen Data Freeze 3 comprised 135,300 Finnish participants, with phenotypes derived from International Classification of Diseases (8^th^, 9^th^, and 10^th^ revision) diagnosis codes from national Finnish hospital discharge and cause-of-death registries as a part of the FinnGen project (**Supplementary Table 22**).

FinnGen samples were genotyped with Illumina and Affymetrix arrays (Thermo Fisher Scientific, Santa Clara, CA, USA). Genotype imputation was carried out by using the population-specific SISu v3 imputation reference panel with Beagle 4.1 (version 08Jun17.d8b, https://faculty.washington.edu/browning/beagle/b4_1.html) as described in the following protocol: dx.doi.org/10.17504/protocols.io.nmndc5e. Post-imputation quality control involved excluding variants with INFO score < 0.7.

We estimated a full weighting matrix for each SNP from the corresponding coefficients of the regression model, then applied the per-SNP weighted model to individuals in FinnGen. To assess the risk for incident first disease events, hazard ratios and 95% confidence intervals per SD increment and by PRS bins (<1%, 1-5%, 40-60%, 95-99%, and >99%) were estimated with Cox proportional hazards models after evaluation of the proportionality assumption. With age as the time scale, the survival models were stratified by sex and adjusted for batch, and the first ten principal components of ancestry calculated within Finns.

## Acknowledgments

This research has been conducted using the UK Biobank Resource under Application Number 24983, “Generating effective therapeutic hypotheses from genomic and hospital linkage data” (http://www.ukbiobank.ac.uk/wp-content/uploads/2017/06/24983-Dr-Manuel-Rivas.pdf). Statin adjustment analyses were further conducted via application 7089 using a protocol approved by the Partners HealthCare Institutional Review Board. We thank all the participants in the UK Biobank and FinnGen studies. We thank members of Pritchard and Bejerano labs for their feedback. This work was supported by National Human Genome Research Institute (NHGRI) of the National Institutes of Health (NIH) under awards R01HG010140 (M.A.R.), R01EB001988-21 (T.H.) and R01HG008140 (J.K.P.). The content is solely the responsibility of the authors and does not necessarily represent the official views of the National Institutes of Health. N.S.A. is supported by the Department of Defense through a National Defense Science and Engineering Grant and by a Stanford Graduate Fellowship. Figure 3B is modified version of Renal corpuscle structure © from Shypoetess 2019. From Wikimedia Commons. M.O is supported by Academy of Finland (#309643). Y.T. is supported by Funai Overseas Scholarship from Funai Foundation for Information Technology and the Stanford University School of Medicine. F.R. is supported by a National Heart, Lung, and Blood Institute grant (1K01HL144607). FinnGen is supported by Abbvie, Astra Zeneca, Biogen, Celgene, Genentech, GSK, Merck, Pfizer, and Sanofi. M.A.R. is supported by Stanford University and a National Institute of Health center for Multi- and Trans-ethnic Mapping of Mendelian and Complex Diseases grant (5U01 HG009080).

## Author information

### Author contributions

M.A.R. conceived and designed the study. N.S.A., Y.T., D.A., N.J.M, M.A., G.R.V., J.P.P., J.Q., and M.A.R. carried out the statistical and computational analyses with advice from S.R., V.A., R.T., J.K.P., M.J.D. and T.H. N.S.A., Y.T., D.A., G.V., M.A. and M.A.R. carried out quality control of the data. M.A.R. supervised computational and statistical aspects of the study. The manuscript was written by N.S.A., Y.T., D.A., G.V., M.A. and M.A.R.; and revised by all the co-authors. All co-authors have approved of the final version of the manuscript.

### Data availability

Data is displayed in the Global Biobank Engine (https://biobankengine.stanford.edu). Analysis scripts and notebooks are available on GitHub at https://github.com/rivas-lab/public-resources/tree/master/uk_biobank/biomarkers.

