## Supplementary Note for "Genetics of 38 blood and urine biomarkers in the UK Biobank"

**Population structure analysis** **2**
Phenotype distributions 3
Genetics of laboratory tests 12
Comparison of effect sizes with published studies 12
Biomarker associated variants prioritize therapeutic targets 14
CNVs influencing lab phenotypes 22
Global and local heritability of biomarkers 24
Cell type decomposition of genetic effects 25
Targeted phenome-wide association study 27
**Correlation of genetic effects between laboratory tests, diseases, and medically relevant phenotypes** **30**
**Causal inference** **33**
Mendelian randomization 33
Latent causal variables 33
**Polygenic prediction of biomarkers within and across populations** **34**
**Multiple regression with PRSs for laboratory tests improves prediction of traits and diseases** **39**
**Reference** **43**
FinnGen 45

### Population structure analysis

**(A)**

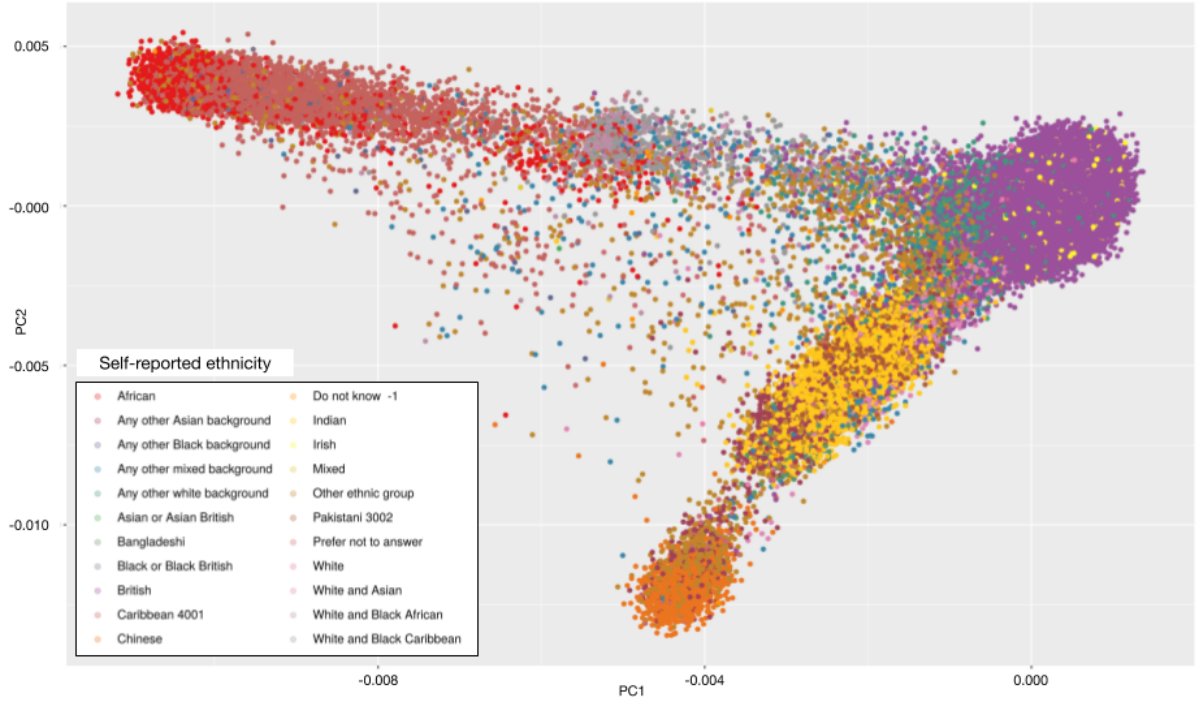


**(B)**

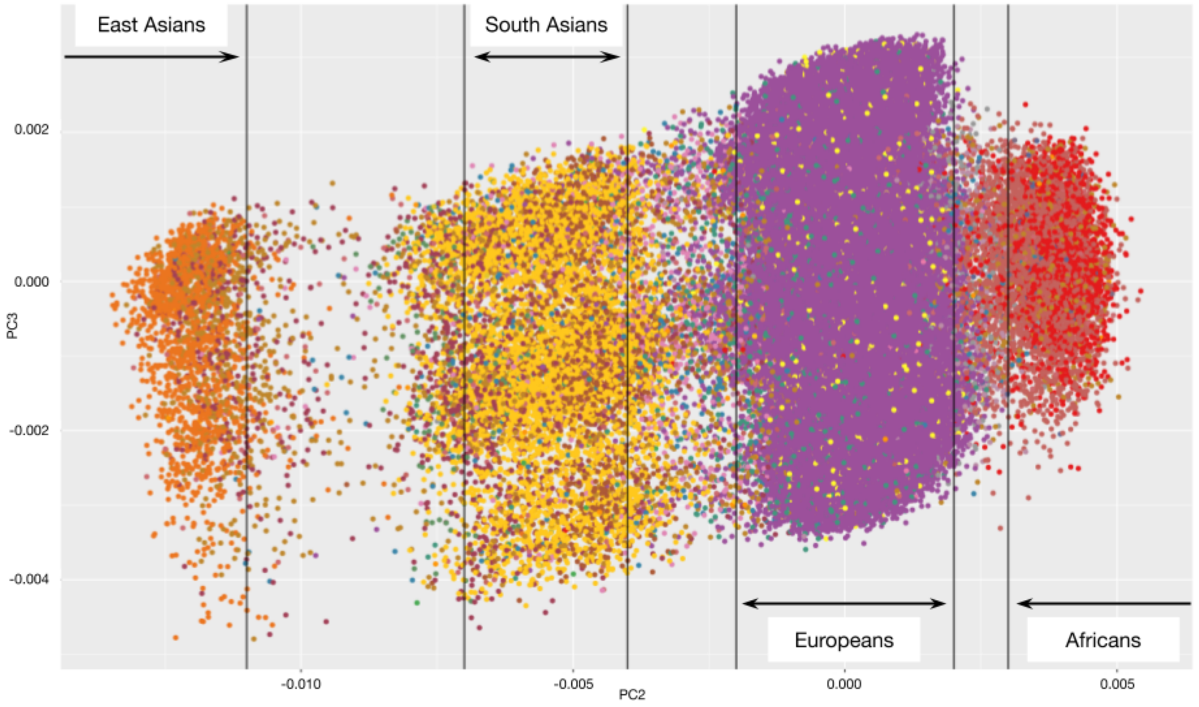


**Supplementary Figure 1. Population structure analysis for UK Biobank with a combination of principal component analysis and self-reported ethnicities.** The 487,409 subjects in the UK Biobank with the genotype data is projected to the principal components and shown on the PC1 vs. PC2 (A) and PC2 vs. PC3 (B) plots. Color represents the self-reported ethnicity (shown in panel A).

#### Phenotype distributions

Within the UK Biobank we estimated what the adjustments might be for statin treatment effect on LDL, TGs, and the rest of the lab phenotypes. As background, there are pre-existing estimates in the literature for LDL[1].
About 20,000 individuals returned for a repeated assessment. Of those, 1,705 either started or stopped a statin between enrollment and that second visit.

**N=1427** people were on statins at the second visit but had not been on them at enrollment. Their **LDL** direct, on average, came down by 1.37 mmol/L. Or, instead treating it as as a multiplier, it came down to 0.68x. So, it seems that a reasonable estimate would be to either add 1.37 or multiply by (1/0.68) for anyone on statins at baseline. This is close to the literature value of a 0.7 correction factor.

All lab phenotypes were assessed for adjustment based on statin usage:

**Supplementary Table 1. Estimated adjustment based on statin usage.** For serum lab phenotypes we estimated the additional constant effect (“Additive”) that statins seem to have on the trait once they are started; "Multiplicative" alternatively means the multiplier effect of statins; and the P-value is the Wilcoxon signed rank test for paired samples comparing whether the pre- and on-statin values seem to differ meaningfully.

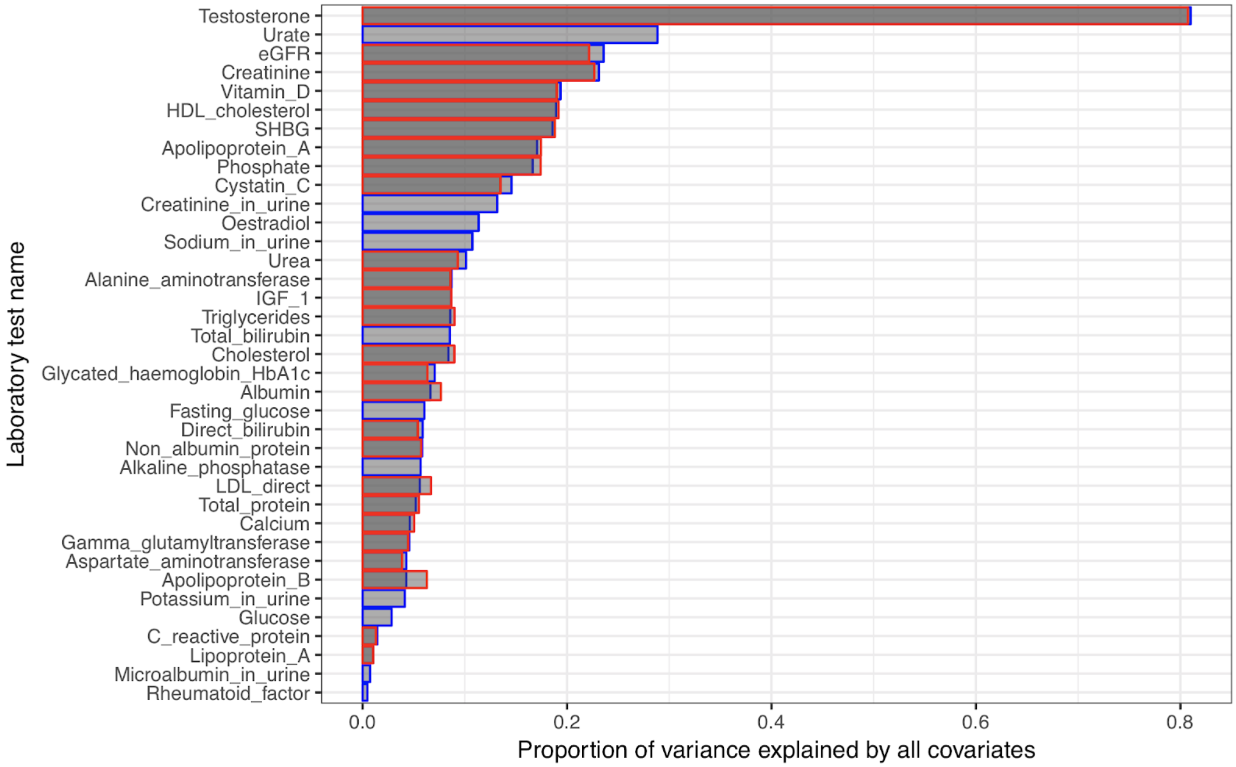


**Supplementary Figure 2A. Proportion of variance explained by all covariates across the 37 raw laboratory phenotypes.** (x-axis) Regression estimate of the proportion of variance explained by all 127 covariates in a linear model for 37 raw laboratory phenotypes including Fasting glucose defined if fasting time between 8 and 24 hours according to Data Field 74 in UK Biobank Data Showcase (y-axis). Blue bar plots indicate estimate before medication adjustment and red bar plots indicate estimate after
50 medication adjustment.

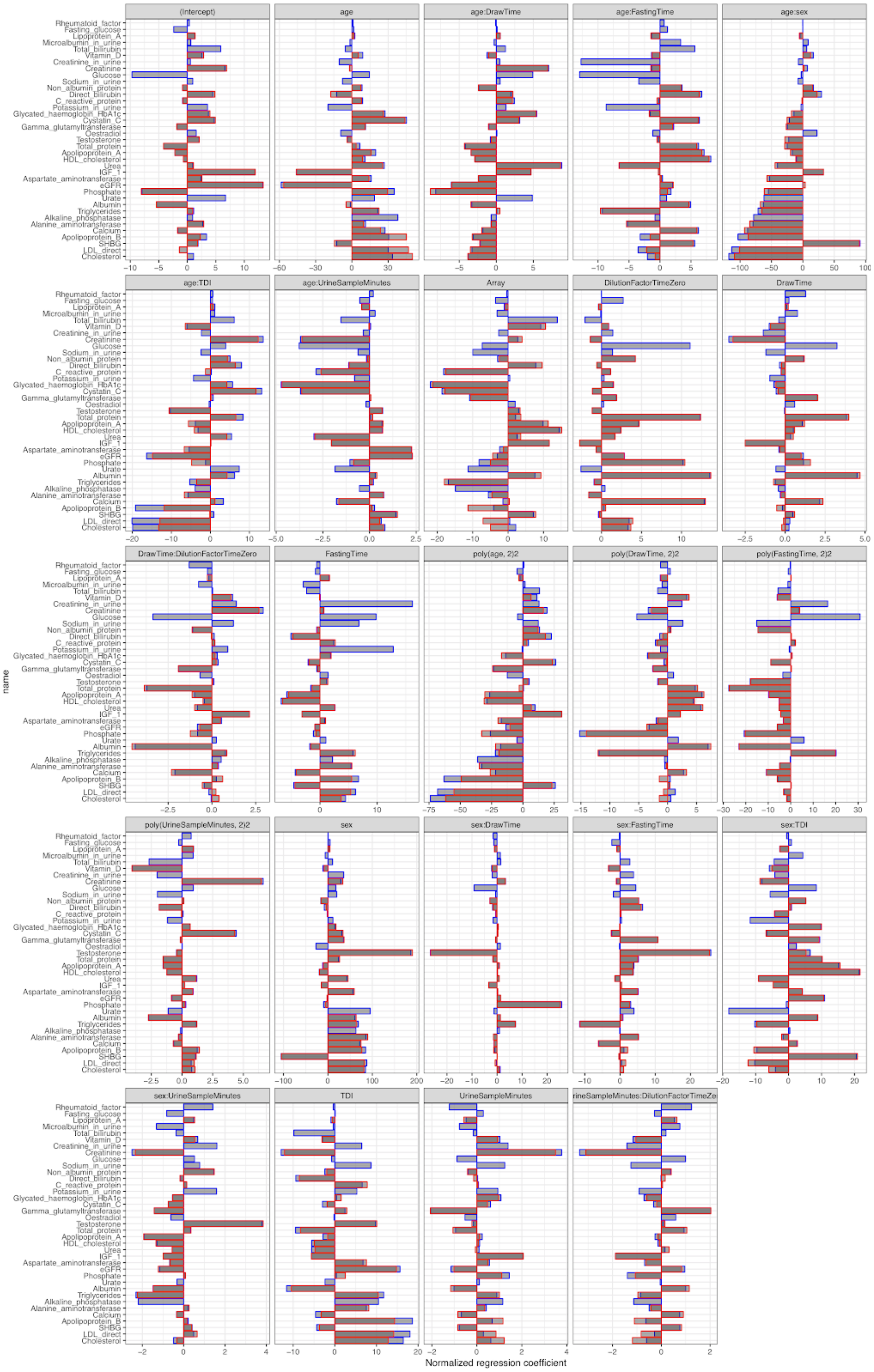
(Intercept)
age
age:DrawTime
age:FastingTime
age:sex
age:TDI
age:UrineSampleMinutes
Array
DilutionFactorTimeZero
DrawTime
DrawTime:DilutionFactorTimeZero
FastingTime
poly(age, 2)2
poly(DrawTime, 2)2
poly(FastingTime, 2)2
poly(UrineSampleMinutes, 2)2
sex
sex:DrawTime
sex:FastingTime
sex:TDI
sex:UrineSampleMinutes
TDI
UrineSampleMinutes
rineSampleMinutes:DilutionFactorTimeZe
name
Rheumatoid_factor
Fasting_glucose
Lipoprotein_A
Microalbumin_in_urine
Total_bilirubin
Vitamin_D
Creatinine_in_urine
Creatinine
Glucose
Sodium_in_urine
Non_albumin_protein
Direct_bilirubin
C_reactive_protein
Potassium_in_urine
Glycated_haemoglobin_HbA1c
Cystatin_C
Gamma_glutamyltransferase
Oestradiol
Testosterone
Total_protein
Apolipoprotein_A
HDL_cholesterol
Urea
IGF_1
Aspartate_aminotransferase
eGFR
Phosphate
Urate
Albumin
Triglycerides
Alkaline_phosphatase
Alanine_aminotransferase
Calcium
Apolipoprotein_B
SHBG
LDL_direct
Cholesterol
Normalized regression coefficient

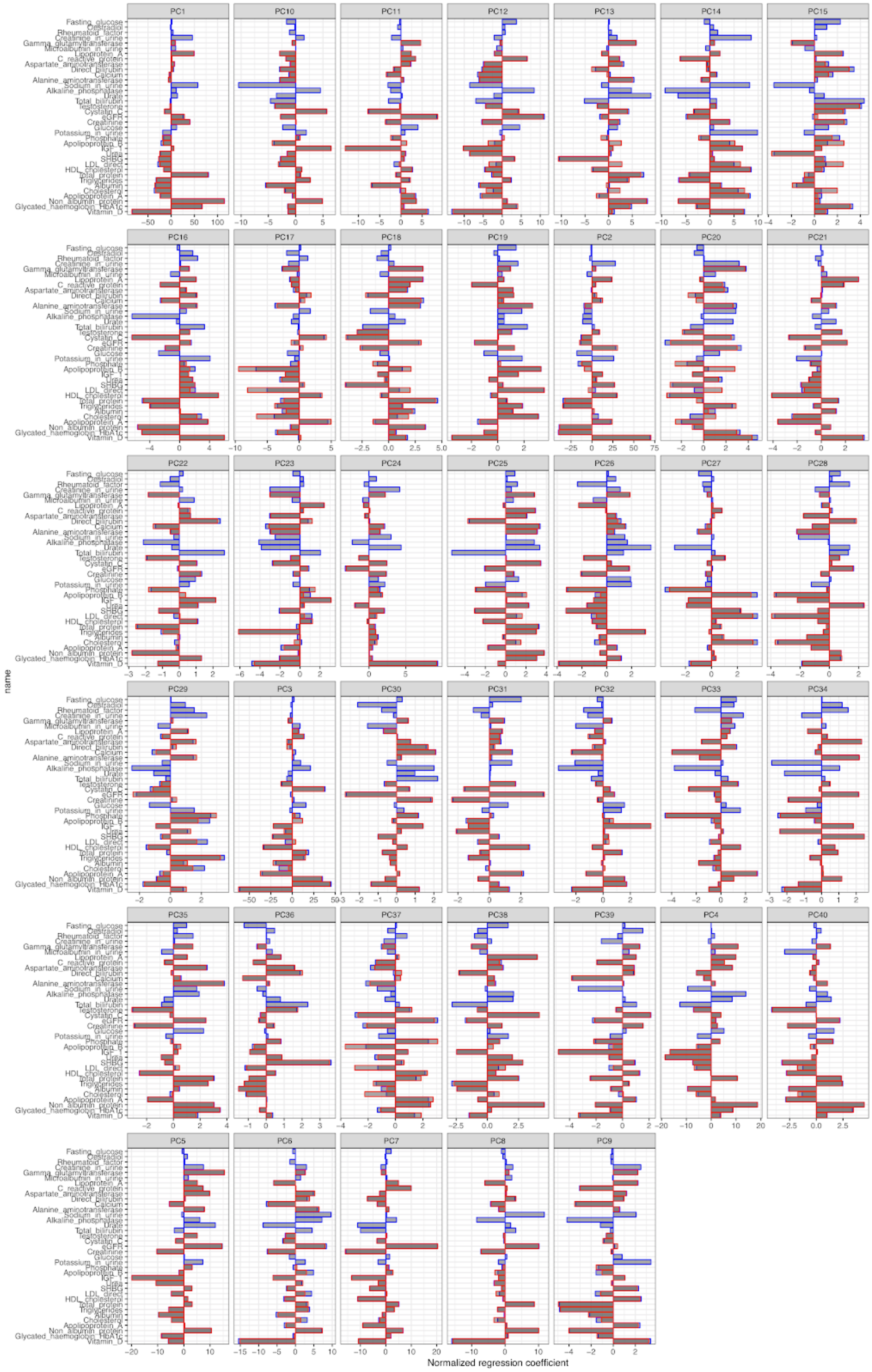
PC1
PC10
PC11
PC12
PC13
PC14
PC15
PC16
PC17
PC18
PC19
PC2
PC20
PC21
PC22
PC23
PC24
PC25
PC26
PC27
PC28
PC29
PC3
PC30
PC31
PC32
PC33
PC34
PC35
PC36
PC37
PC38
PC39
PC4
PC40
PC5
PC6
PC7
PC8
PC9
name
Fasting_glucose
Oestradiol
Rheumatoid_factor
Creatinine_in_urine
Gamma_glutamyltransferase
Microalbumin_in_urine
Lipoprotein_A
C_reactive_protein
Aspartate_aminotransferase
Direct_bilirubin
Calcium
Alanine_aminotransferase
Sodium_in_urine
Alkaline_phosphatase
Urate
Total_bilirubin
Testosterone
Cystatin_C
eGFR
Creatinine
Glucose
Potassium_in_urine
Phosphate
Apolipoprotein_B
IGF_1
Urea
SHBG
LDL_direct
HDL_cholesterol
Total_protein
Triglycerides
Albumin
Cholesterol
Apolipoprotein_A
Non_albumin_protein
Glycated_haemoglobin_HbA1c
Vitamin_D
Normalized regression coefficient

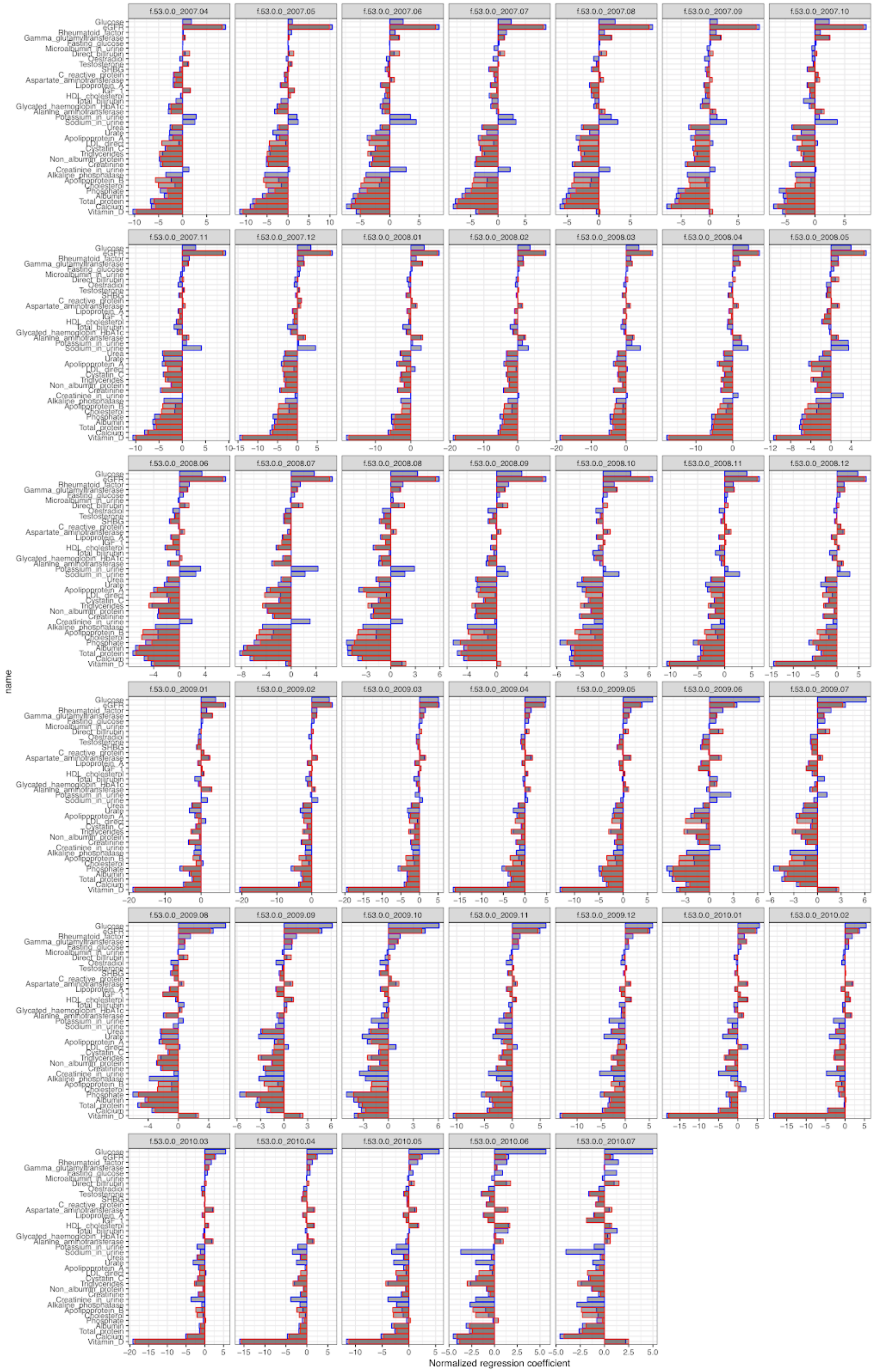
f.53.0.0_2007.04
f.53.0.0_2007.05
f.53.0.0_2007.06
f.53.0.0_2007.07
f.53.0.0_2007.08
f.53.0.0_2007.09
f.53.0.0_2007.10
f.53.0.0_2007.11
f.53.0.0_2007.12
f.53.0.0_2008.01
f.53.0.0_2008.02
f.53.0.0_2008.03
f.53.0.0_2008.04
f.53.0.0_2008.05
f.53.0.0_2008.06
f.53.0.0_2008.07
f.53.0.0_2008.08
f.53.0.0_2008.09
f.53.0.0_2008.10
f.53.0.0_2008.11
f.53.0.0_2008.12
f.53.0.0_2009.01
f.53.0.0_2009.02
f.53.0.0_2009.03
f.53.0.0_2009.04
f.53.0.0_2009.05
f.53.0.0_2009.06
f.53.0.0_2009.07
f.53.0.0_2009.08
f.53.0.0_2009.09
f.53.0.0_2009.10
f.53.0.0_2009.11
f.53.0.0_2009.12
f.53.0.0_2010.01
f.53.0.0_2010.02
f.53.0.0_2010.03
f.53.0.0_2010.04
f.53.0.0_2010.05
f.53.0.0_2010.06
f.53.0.0_2010.07
name
Glucose
eGFR
Rheumatoid_factor
Gamma_glutamyltransferase
Fasting_glucose
Microalbumin_in_urine
Direct_bilirubin
Oestradiol
Testosterone
SHBG
C_reactive_protein
Aspartate_aminotransferase
Lipoprotein_A
IGF_1
HDL_cholesterol
Total_bilirubin
Glycated_haemoglobin_HbA1c
Alanine_aminotransferase
Potassium_in_urine
Sodium_in_urine
Urea
Urate
Apolipoprotein_A
LDL_direct
Cystatin_C
Triglycerides
Non_albumin_protein
Creatinine
Creatinine_in_urine
Alkaline_phosphatase
Apolipoprotein_B
Cholesterol
Phosphate
Albumin
Total_protein
Calcium
Vitamin_D
Normalized regression coefficient

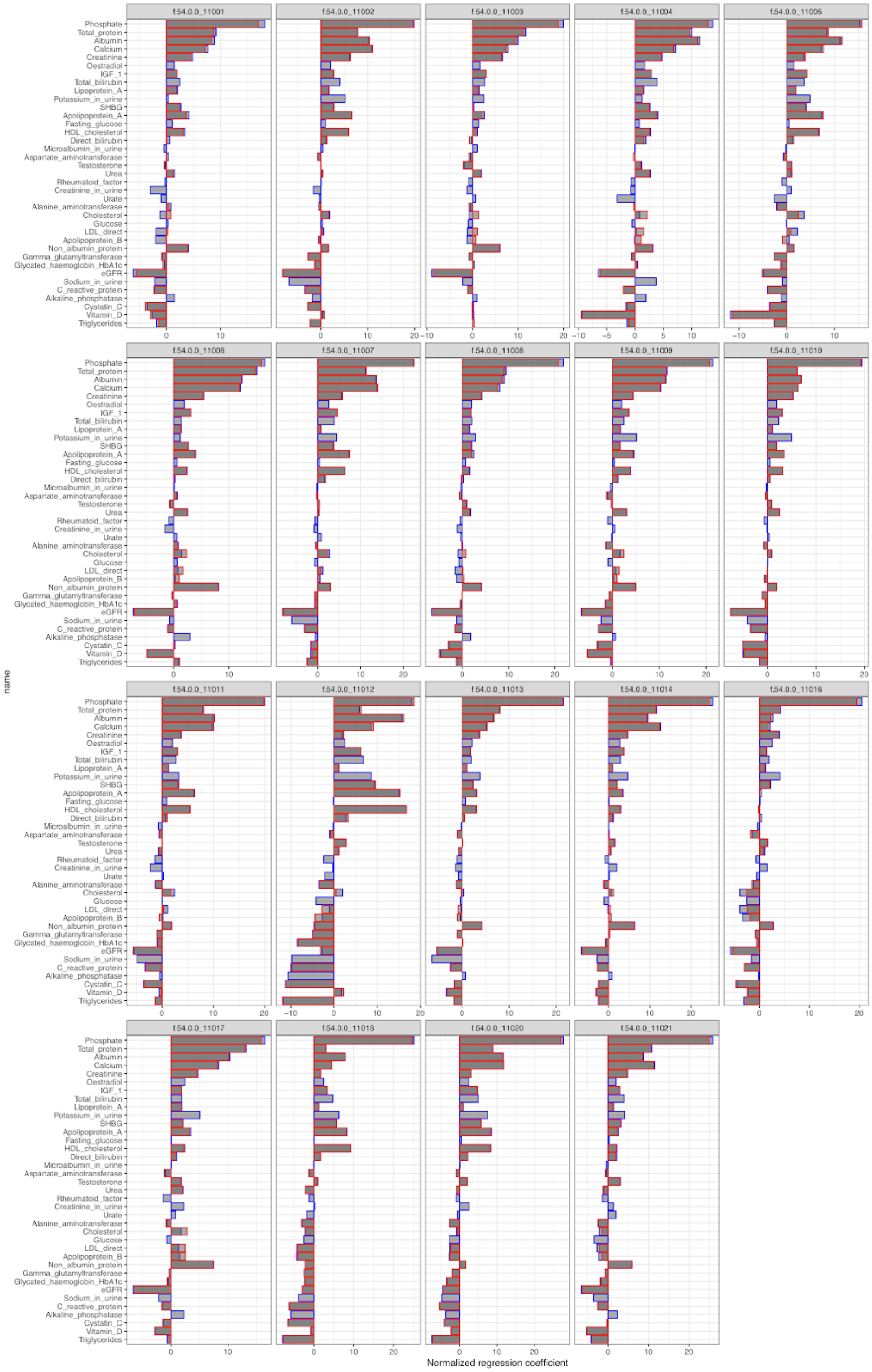


**Supplementary Figure 2B. Normalized regression coefficient for the 37 raw laboratory phenotypes across the covariates.** (x-axis) Normalized regression coefficient for 23 covariates in a linear model for the 37 raw laboratory phenotypes including Fasting glucose defined if fasting time between 8 and 24 hours according to Data Field 74 in UK Biobank Data Showcase (y-axis). Blue bar plots indicate estimate before medication adjustment and red bar plots indicate estimate after medication adjustment.

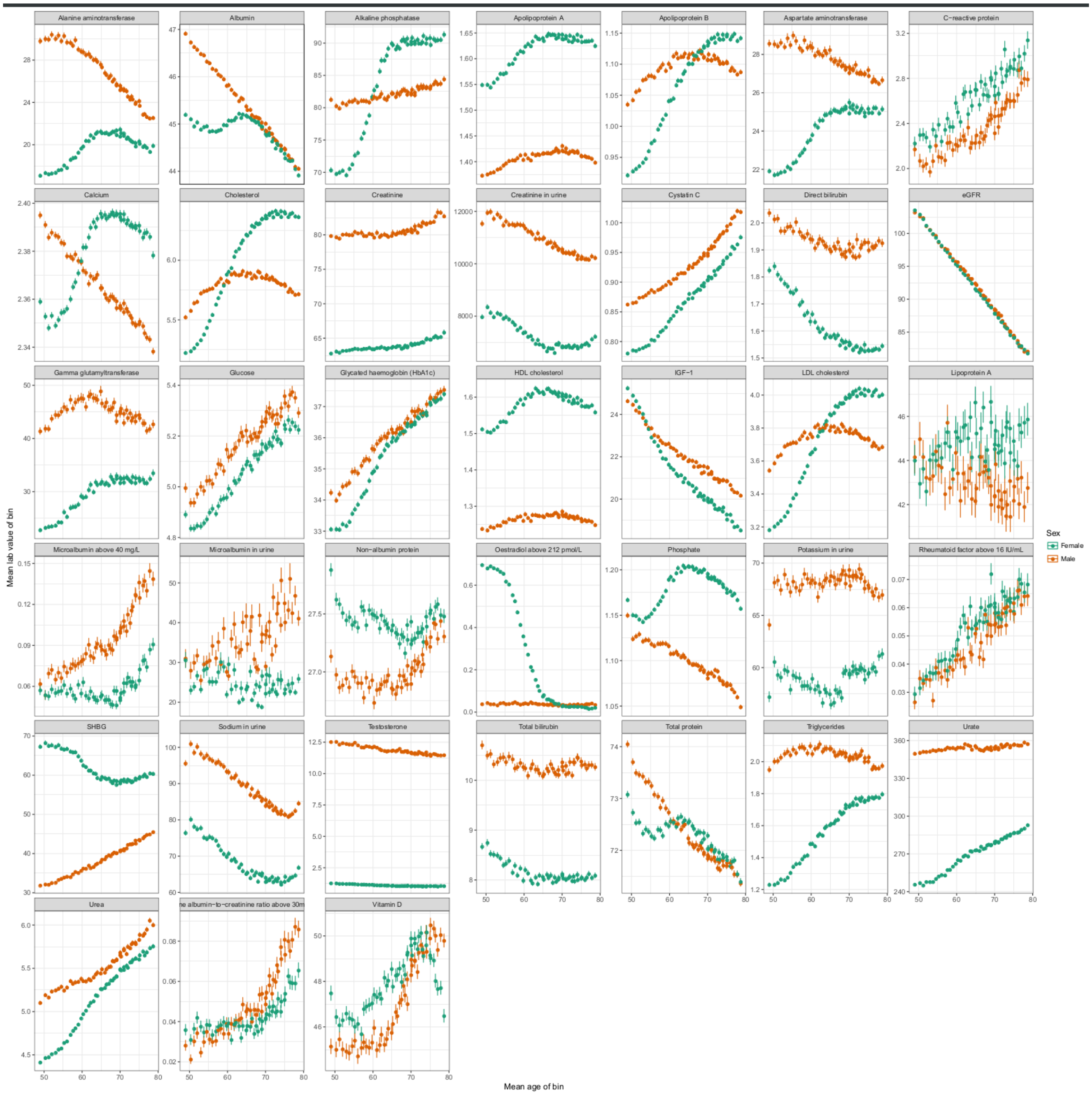


**Supplementary Figure 2C. Phenotype distributions of all laboratory tests by age and sex.** (x-axis) Age of individuals within a pentacontile were averaged. (y-axis) The corresponding average value +/- 1 SD of each laboratory test measurement for all individuals with available data in the study. Color indicates the reported sex of the individuals (orange = male, turquoise = female).

**Supplementary Table 2. Description of 38 measured and derived lab phenotypes.** Lab phenotype name, abbreviation, units of measurement, the UK Biobank field ID, Global Biobank Engine phenotype ID, whether the phenotype is defined as binary (B) or quantitative (Q), whether the phenotype is adjusted for statin (Y) or not (N), whether the phenotype is adjusted for covariates (Y) or not (N), and total number of unrelated individuals across the self-identified White British, self-identified non-British White, African, East Asian, South Asian population subset in UK Biobank, the number of loci identified from GWAS (the number of independent loci, the number of imputed variants on 1000 genome phase 3 MAF > 1% variants, number of protein-altering variants, number of PTVs, the number of HLA alleles with posterior probability >= 0.8, the number of single CNVs, and the number of rare aggregate CNVs), and GBE URL.

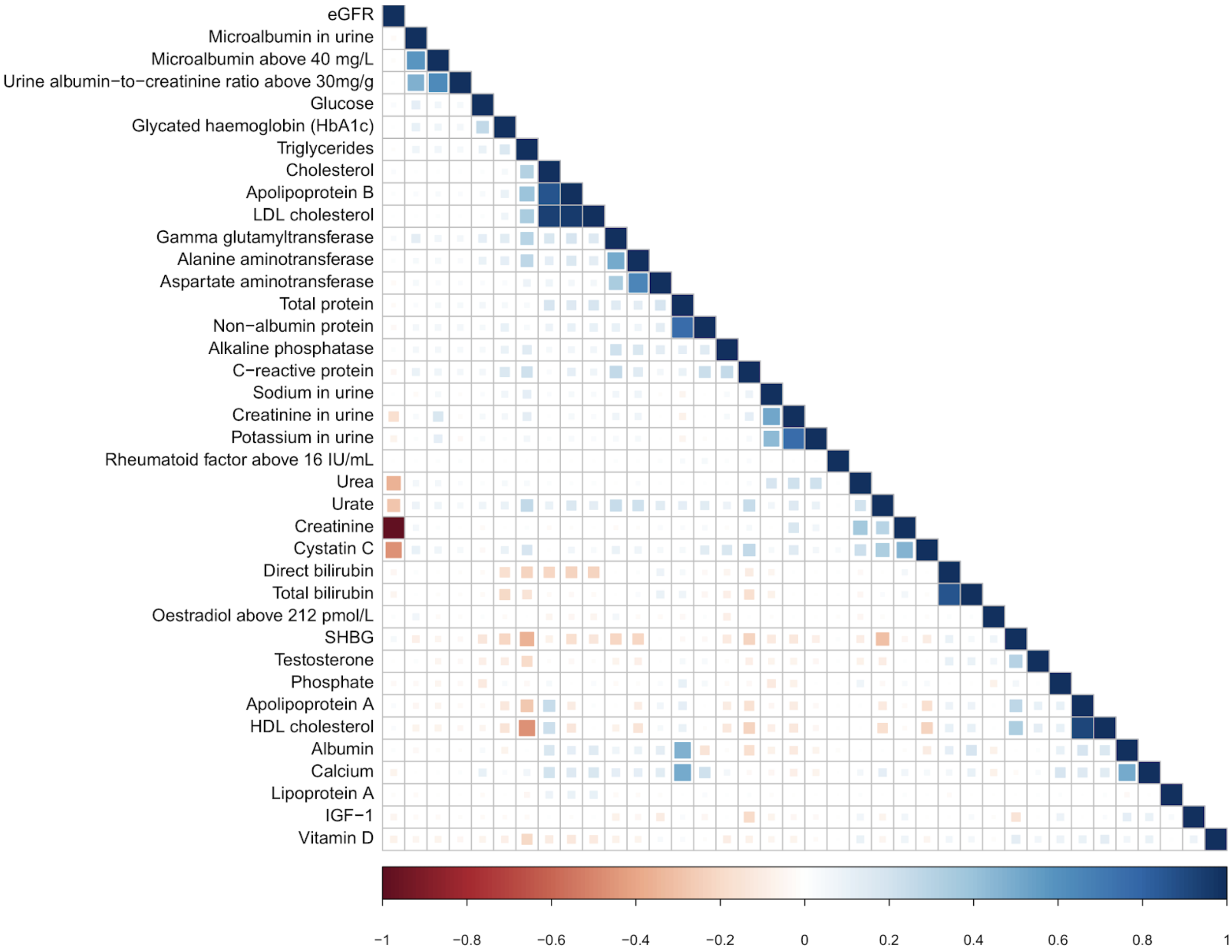


**Supplementary Figure 3. Phenotype correlation among the 38 lab phenotypes.** -1 (red) to 1 (blue) correlation of phenotypes (cell size indicates correlation). Only cells with $p < 0.001$ are shown. Results are consistent with previous work, and captures known associations between both testosterone and SHBG with uric acid (urate) levels[2].

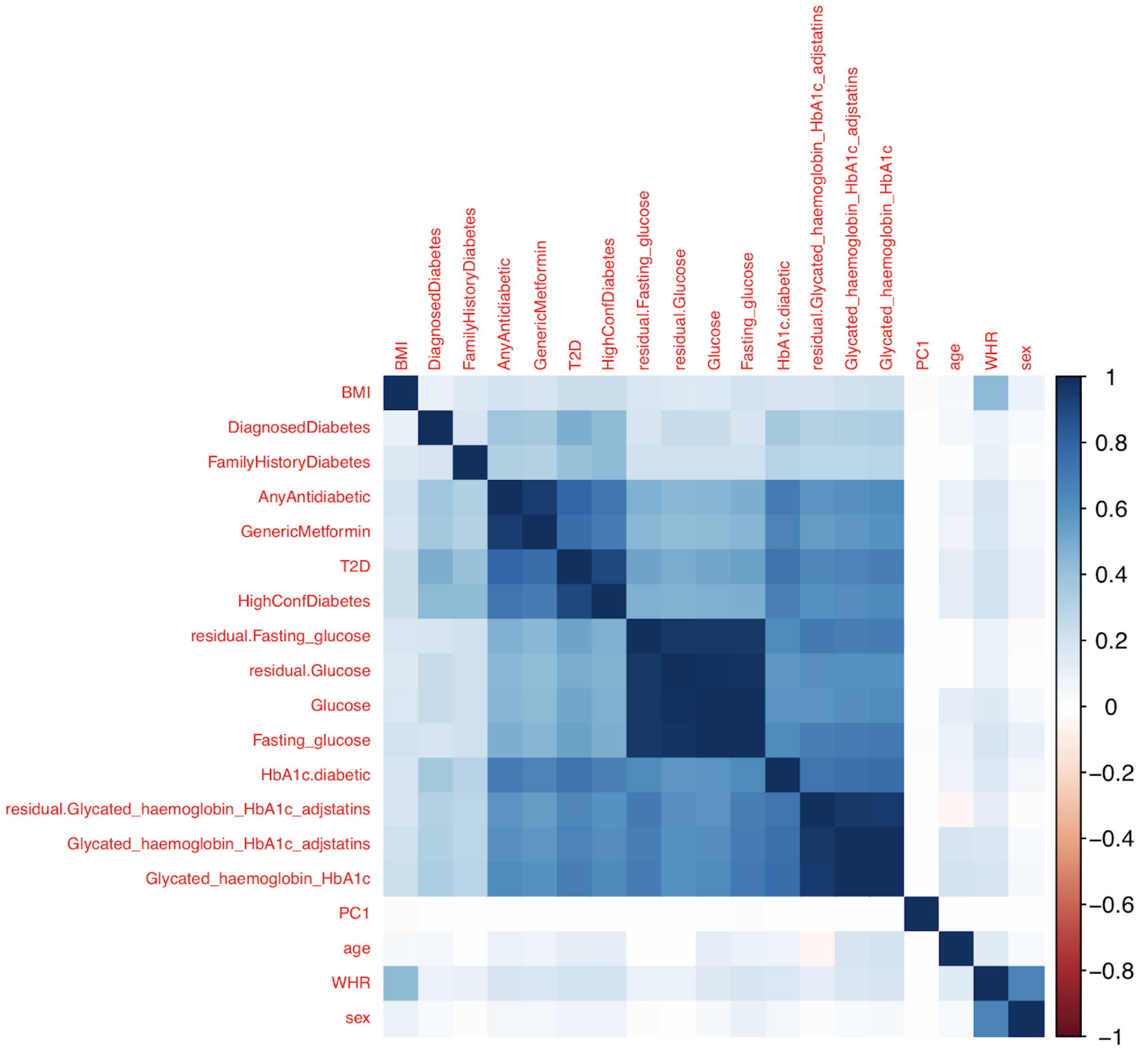


**Supplementary Figure 4. Correlogram of different diabetes- and diabetes-related traits.** The similarity of type 2 diabetes (following Eastwood et al), high confidence diabetes (examining all available timepoints for an individual and using self-report and ICD codes), and prescription of metformin or any oral antidiabetic are compared to the laboratory test measurements of HbA1c and glucose. HbA1c was adjusted for statins (see Methods) and residualized (see Methods), while glucose was subset to individuals with a fasting time between 8 and 24 hours (see Methods) to ensure effects were not driven by fasting. Diagnosed diabetes was defined by the UK Biobank during the nurse interview, and family history was defined as having at least one self reported mother, father, or sibling (non-adopted) with diabetes. Table of correlations presented below (Supplementary Table 3).

**Supplementary Table 3. Diabetes correlation estimates.** For each trait pair, the spearman correlation of the traits across all unrelated White Briitsh individuals for which the traits were defined is presented. AnyAntidiabetic is defined as any non-insulin drug from the oral antidiabetics and metformin codes presented in Eastwood et al; T2D is the definition of type 2 diabetes presented in Eastwood et al; fasting glucose is the glucose measurement for the individuals with a self-reported fasting time between 8 and 24 hours; HighConfDiabetes is a combination of self report and ICD codes presented in (DeBoever et al. 2018); GenericMetformin is just using Metformin and its generic forms; FamilyHistoryDiabetes is defined as 0 or 1 depending on whether the individual has self reported a father, mother, or sibling with diabetes; and HbA1c.diabetic is defined as a binary indicator of the individual having a measured HbA1c greater than 48.

### Genetics of laboratory tests

#### Comparison of effect sizes with published studies

**Supplementary Figure 5. Comparison of estimated effect sizes between UK Biobank and previous GWAS.** (x-axis) UK Biobank estimated effect size. (y-axis) Comparative study estimated effect size. All variants associated p < 1e-6 in either study are shown.

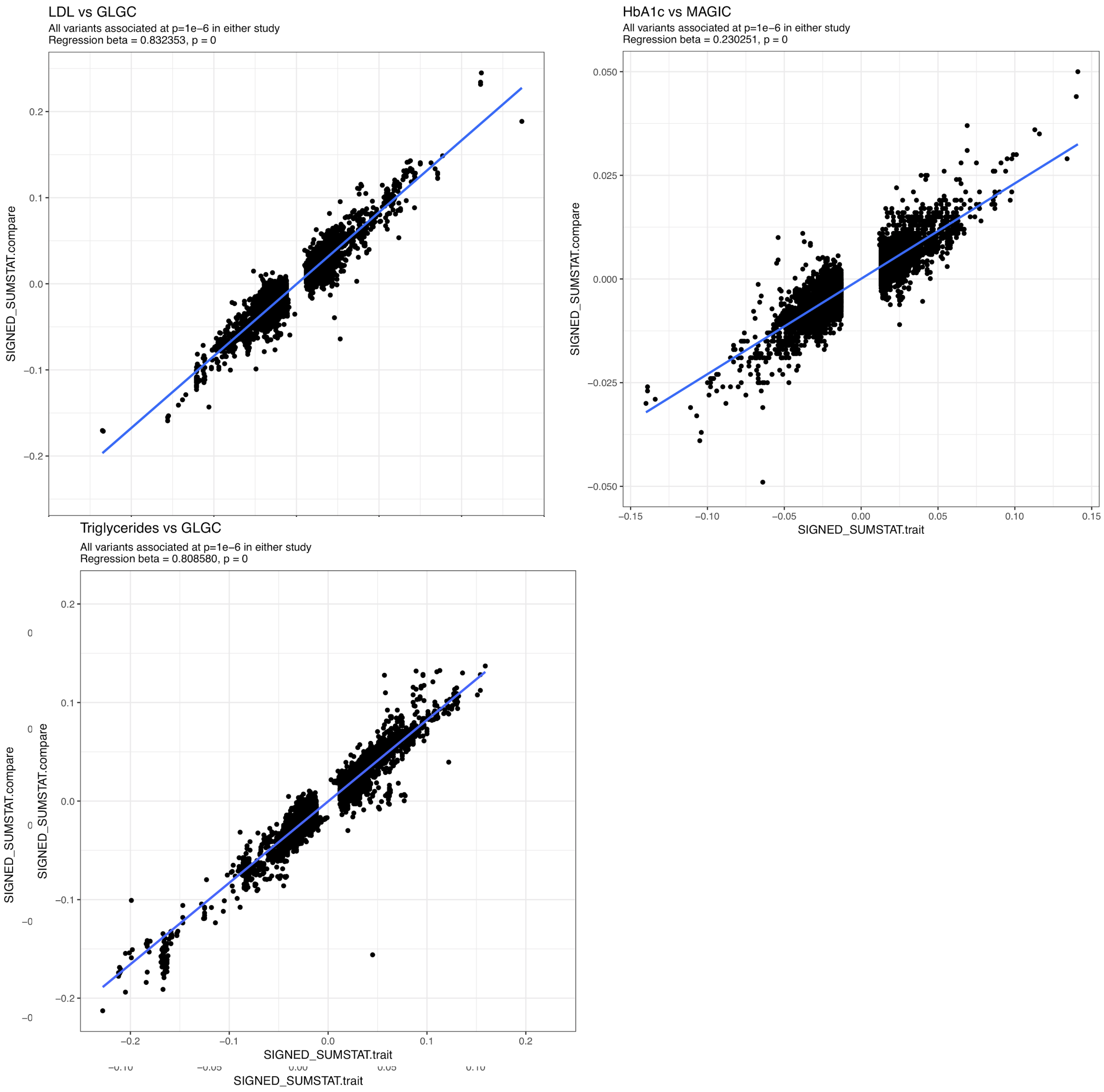

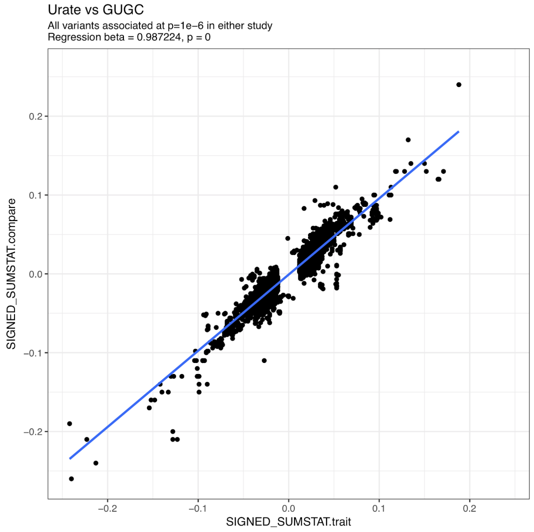


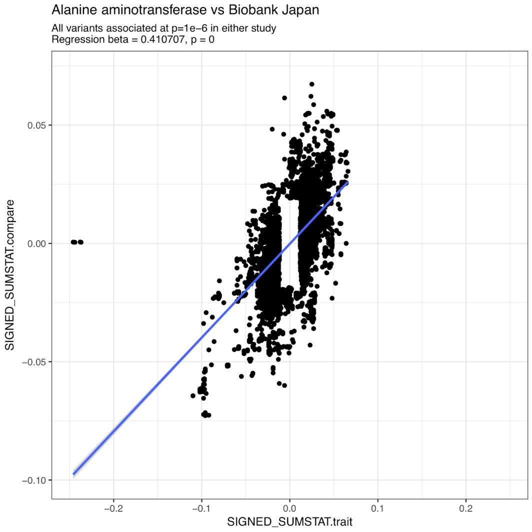


**Supplementary Table 4. Comparison of estimated effect sizes between UK Biobank and other studies.** For each comparison, spearman correlations of beta estimates between the exiting study and UK Biobank laboratory measures are compared. Previous studies are taken from a number of existing papers. Comparisons were made to variants present in both studies with both the set of independent hits with $p < 5e-8$ ascertained from our data (Methods, “hits”) as well as the set of all variants with $p < 1e-6$ in either study (“subthreshold”). For each variant set, the overall correlation, as well as the beta and standard error from a linear regression, are reported.

### Biomarker associated variants prioritize therapeutic targets

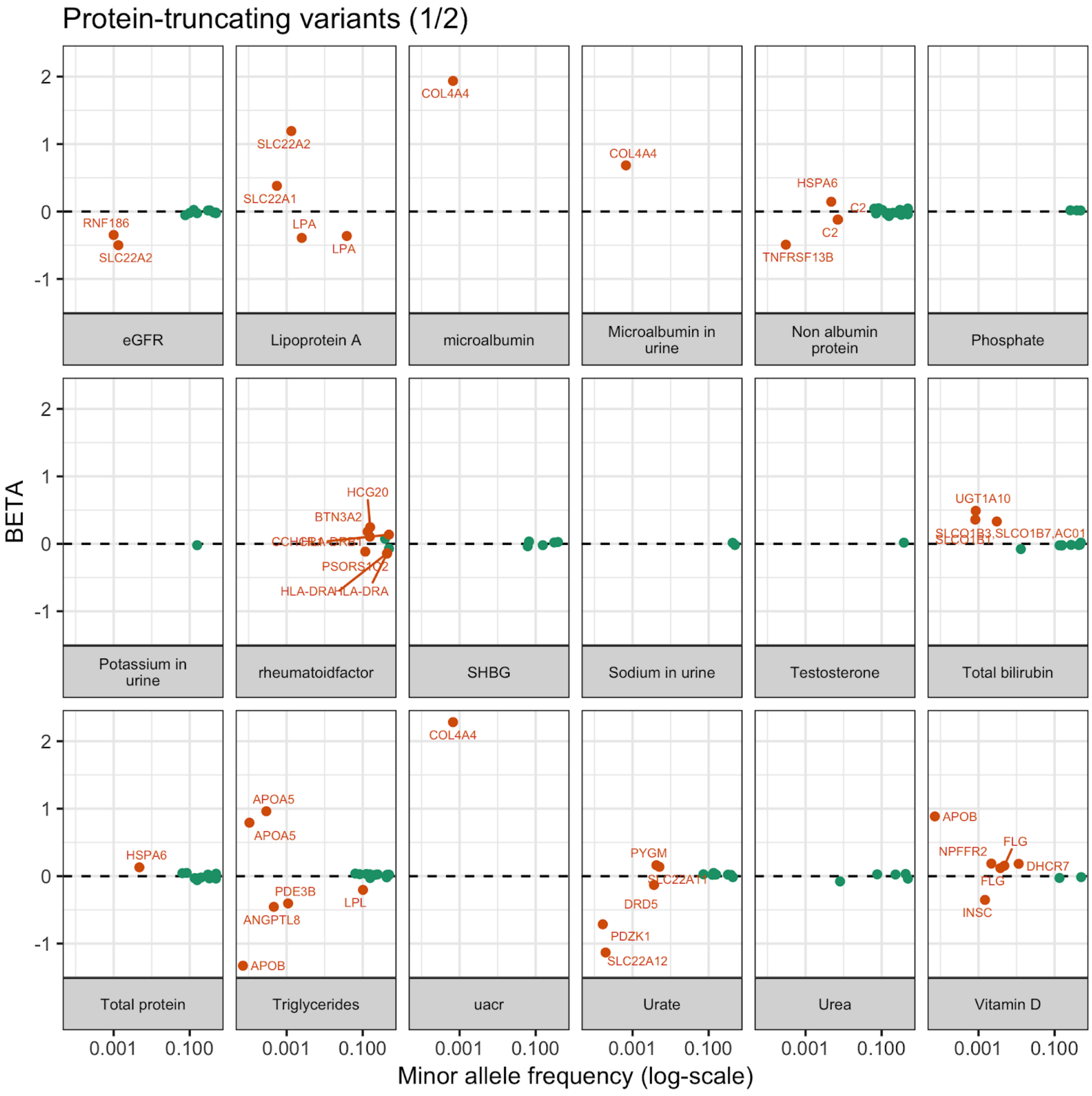

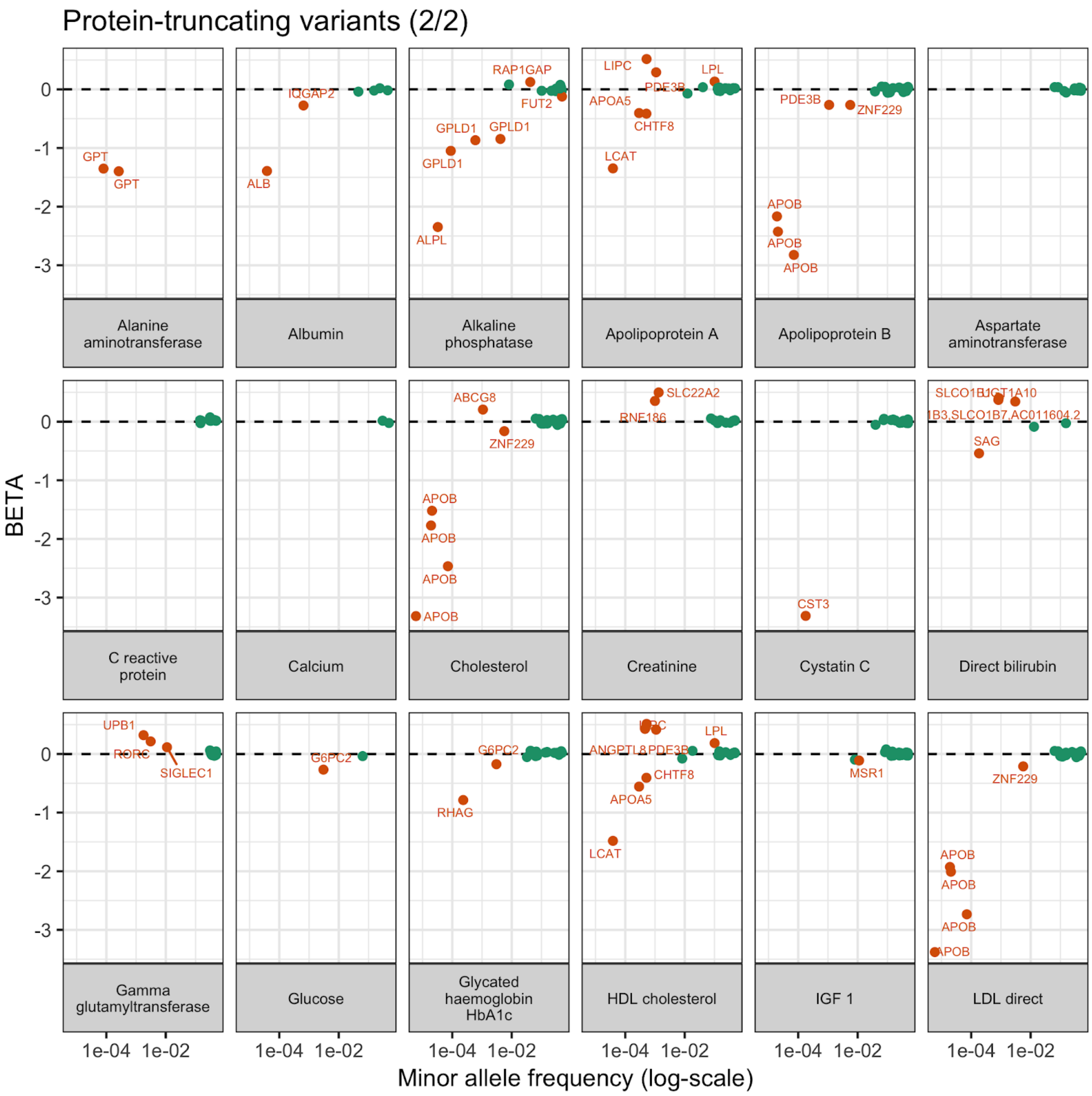


100 **Supplementary Figure 6. Cascade plots for predicted protein-truncating variants across lab phenotypes.** (x-axis) Minor allele frequency of genetic variant associated to phenotype (p < 1e-7) and (y-axis) BETA univariate regression coefficient estimate. Orange and labelled data points include genes with PTVs whose estimated effect size (BETA) is greater than or equal to .1 or less than or equal to -.1 standard deviation (SD). Two phenotypes (Creatinine in urine and oestradiol) did not have PTV associations with p < 1e-7 and excluded from the plot.

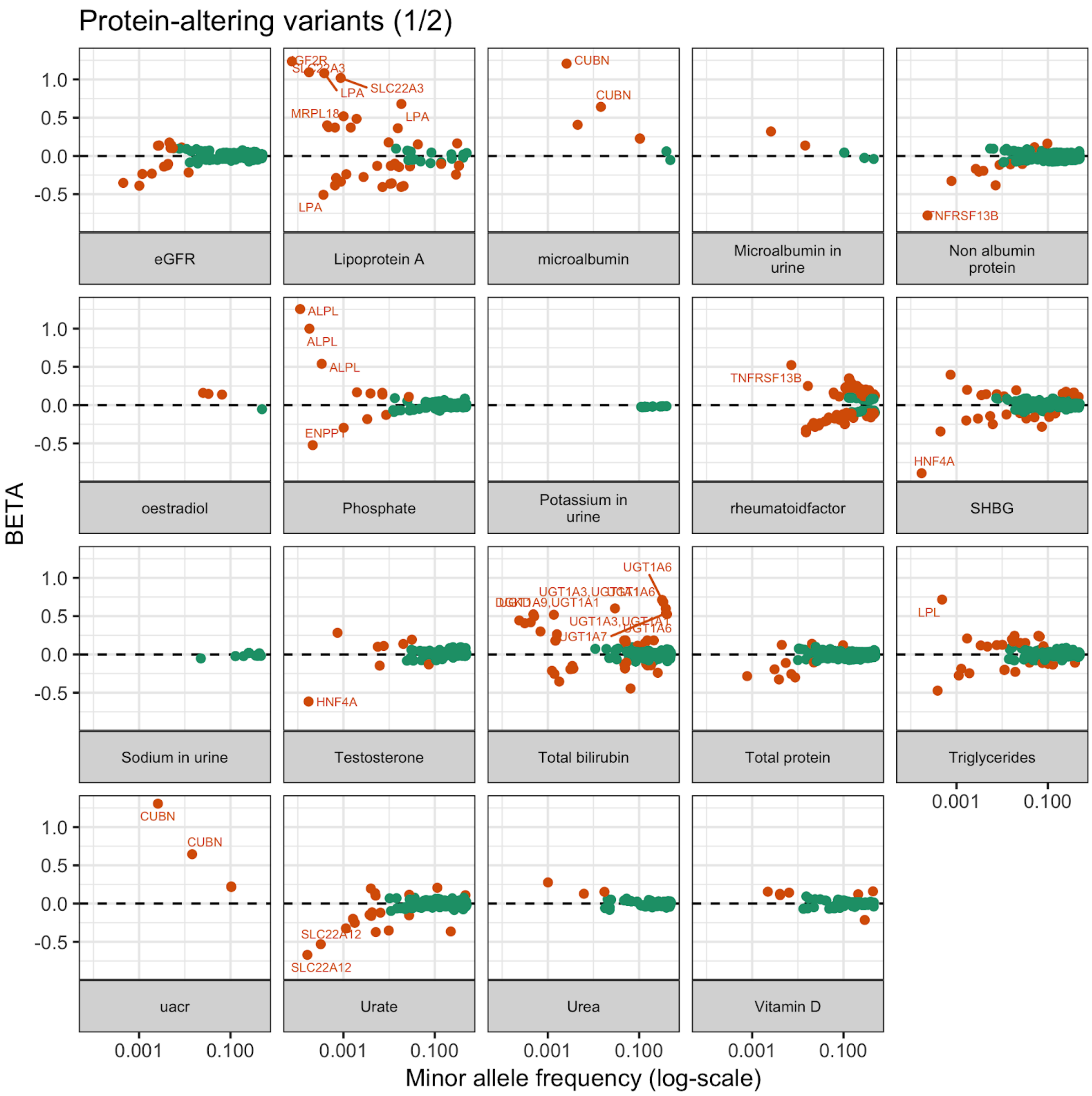

Protein-altering variants (1/2)
BETA
Minor allele frequency (log-scale)
1.0
0.5
0.0
-0.5
0.001
0.100
eGFR
Lipoprotein A
microalbumin
Microalbumin in urine
Non albumin protein
oestradiol
Phosphate
Potassium in urine
rheumatoidfactor
SHBG
Sodium in urine
Testosterone
Total bilirubin
Total protein
Triglycerides
uacr
Urate
Urea
Vitamin D
SLC22A3
LPA
MRPL18
CUBN
TNFRSF13B
ALPL
ENPP1
HNF4A
UGT1A6
UGT1A7
LPL
SLC22A12

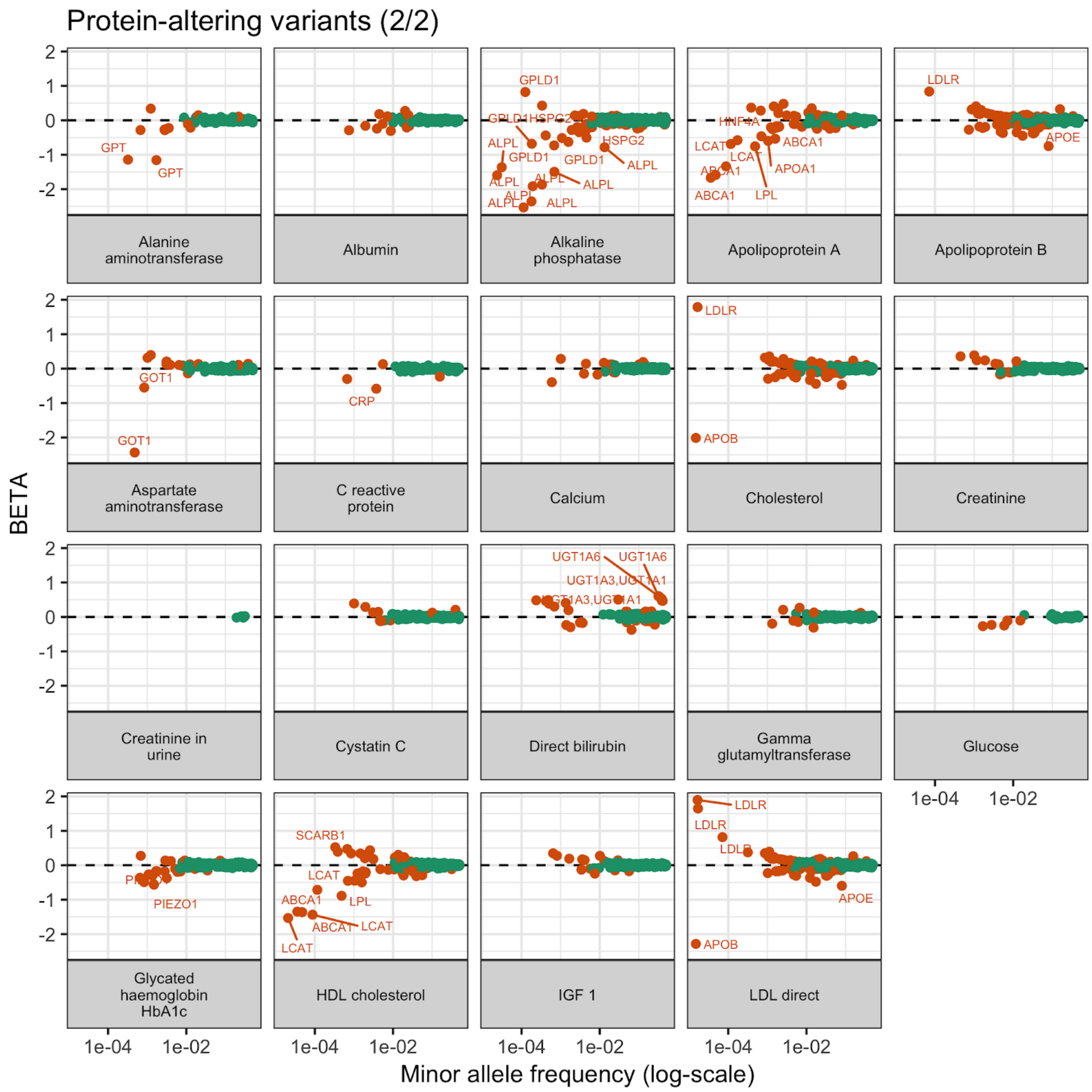


**Supplementary Figure 7. Cascade plots for predicted protein-altering variants across lab phenotypes.** (x-axis) Minor allele frequency of genetic variant associated to phenotype ($p < 1e-7$) and (y-axis) BETA univariate regression coefficient estimate. Orange and labelled data points include genes with protein-altering variants whose estimated effect size (BETA) is greater than or equal to .1 or less than or equal to -.1.

Non-coding variants (1/3)

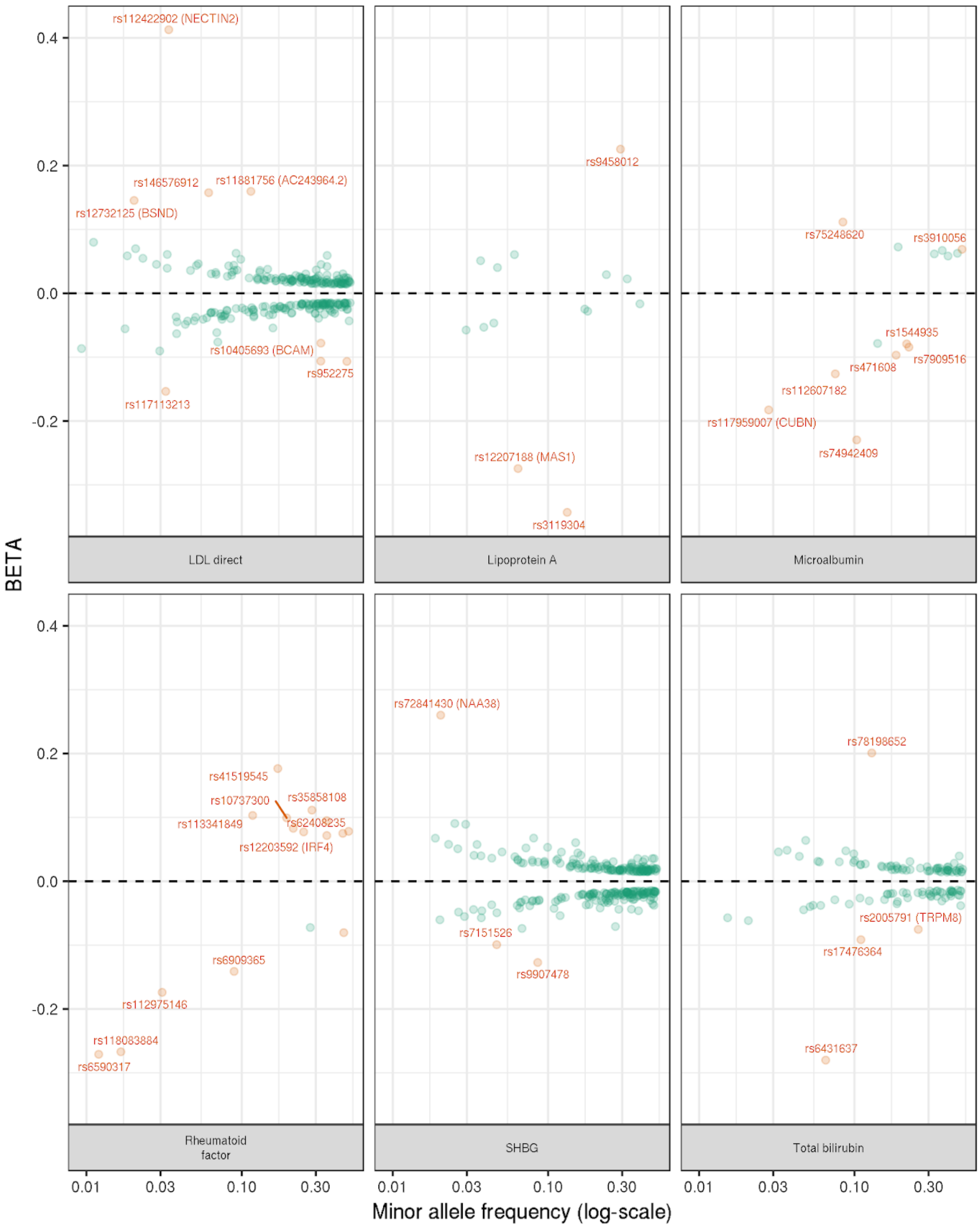

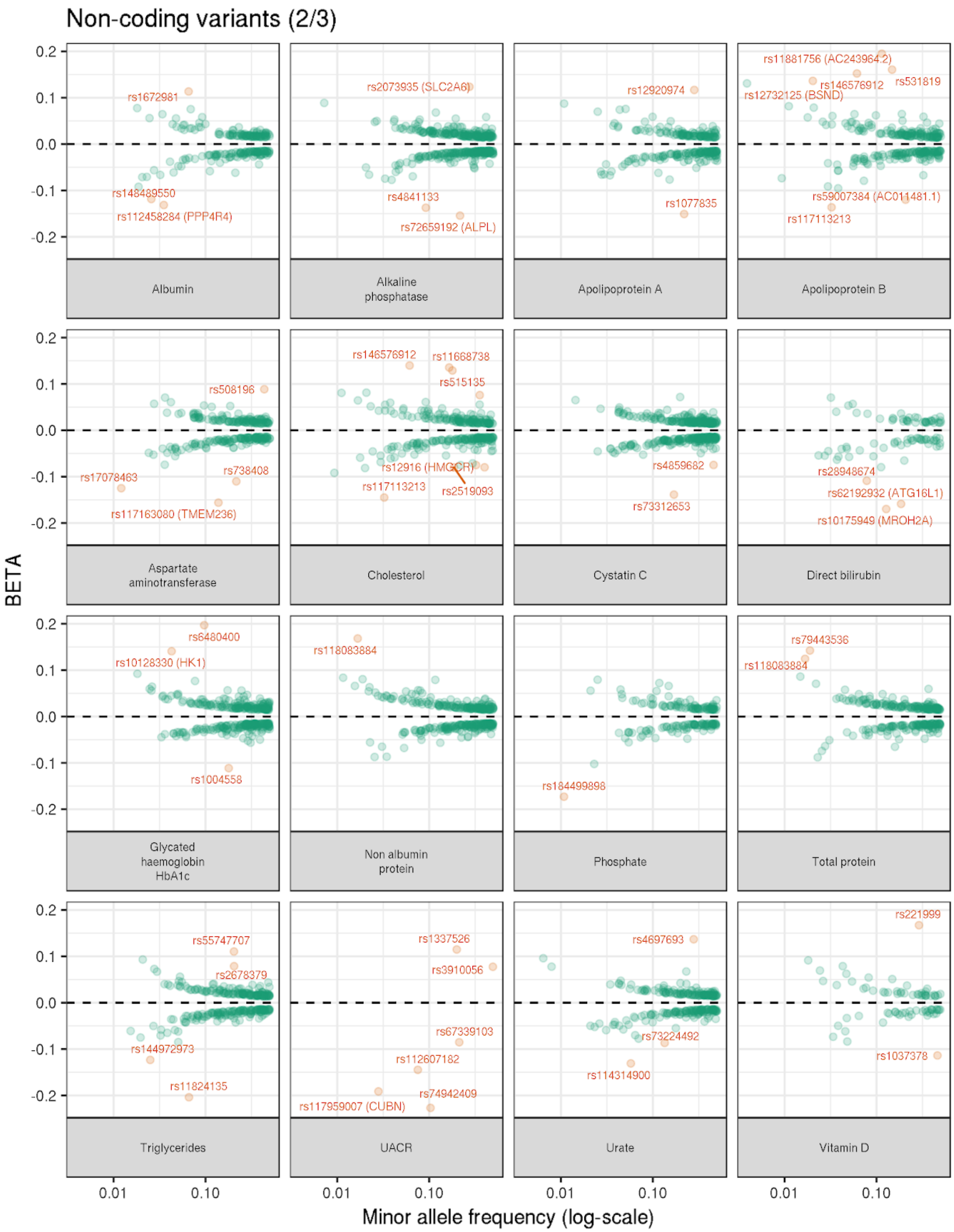
Non-coding variants (2/3)
BETA
Minor allele frequency (log-scale)
0.2
0.1
0.0
-0.1
-0.2
0.01
0.10
Albumin
rs1672981
rs148489550
rs112458284 (PPP4R4)
Alkaline phosphatase
rs2073935 (SLC2A6)
rs4841133
rs72659192 (ALPL)
Apolipoprotein A
rs12920974
rs1077835
Apolipoprotein B
rs11881756 (AC243964.2)
rs146576912
rs531819
rs12732125 (BSND)
rs59007384 (AC011481.1)
rs117113213
Aspartate aminotransferase
rs508196
rs17078463
rs738408
rs117163080 (TMEM236)
Cholesterol
rs146576912
rs11668738
rs515135
rs12916 (HMGCR)
rs117113213
rs2519093
Cystatin C
rs4859682
rs73312653
Direct bilirubin
rs28948674
rs62192932 (ATG16L1)
rs10175949 (MROH2A)
Glycated haemoglobin HbA1c
rs6480400
rs10128330 (HK1)
rs1004558
Non albumin protein
rs118083884
Phosphate
rs184499898
Total protein
rs79443536
rs118083884
Triglycerides
rs55747707
rs2678379
rs144972973
rs11824135
UACR
rs1337526
rs3910056
rs67339103
rs112607182
rs117959007 (CUBN)
rs74942409
Urate
rs4697693
rs73224492
rs114314900
Vitamin D
rs221999
rs1037378

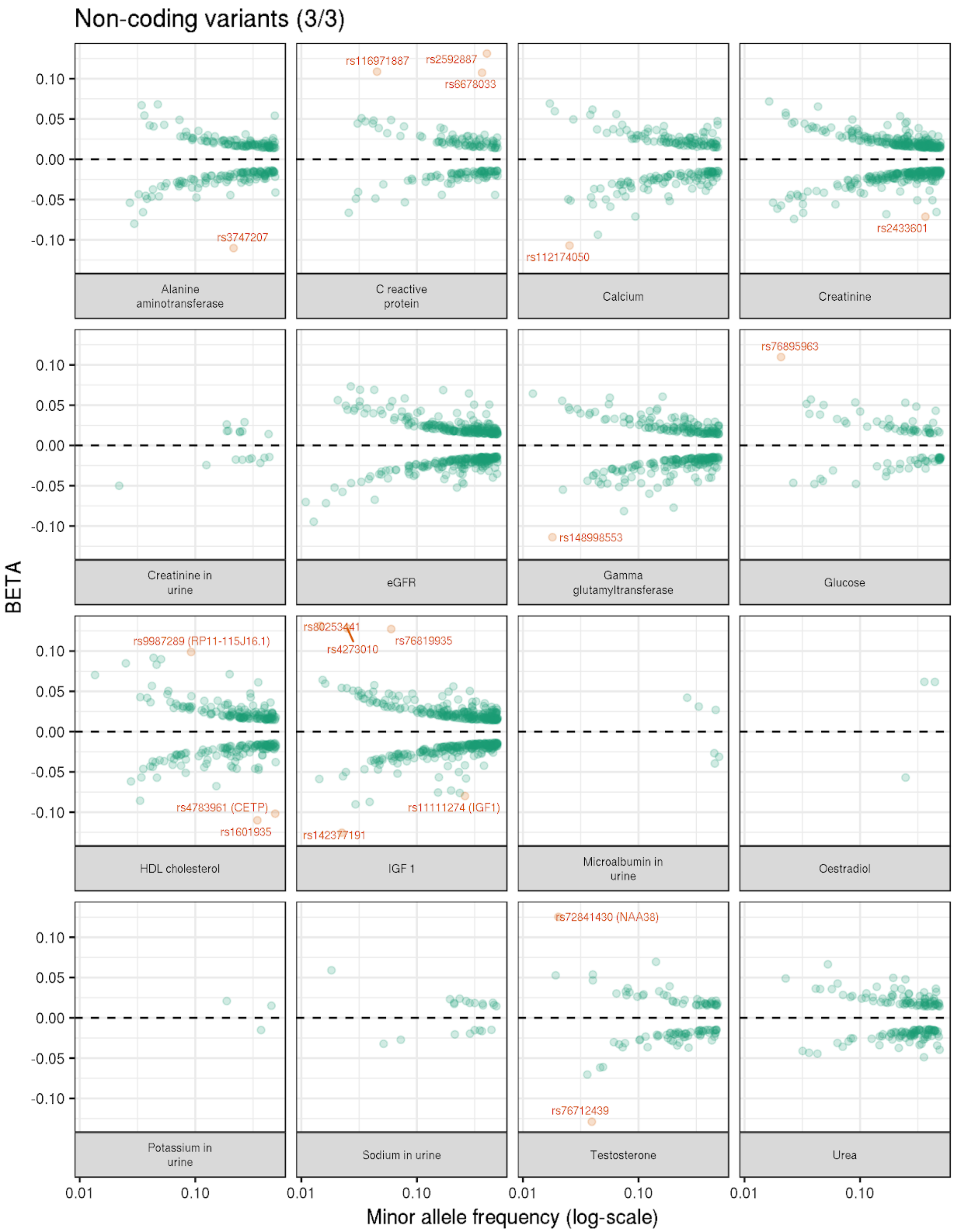


**Supplementary Figure 8. Cascade plots for non-coding variants across lab phenotypes.** (x-axis) Minor allele frequency of non-coding variants characterized on the imputed 1000 Genomes Phase I variant associated to phenotype ($p < 5e-8$) and (y-axis) BETA univariate regression coefficient estimate. Orange and labelled data points include non-coding variants whose estimated effect size (BETA) is an outlier, I.e. absolute value of estimated effect size deviates from the standard deviation range estimated from linear fit between log minor allele frequency and absolute value of estimated effect size (outlier, see methods for more details). The gene symbols are shown for splicing variants.

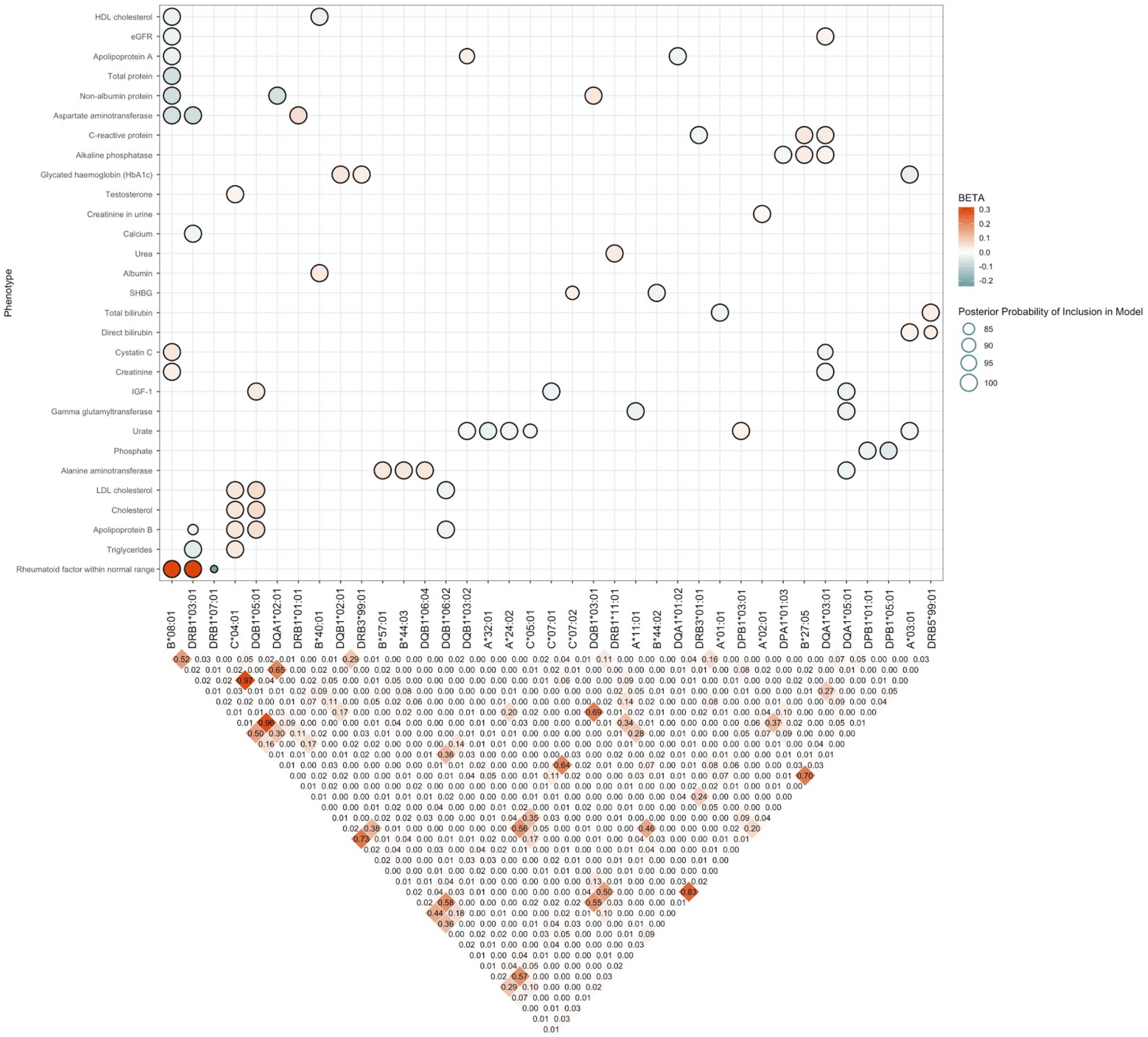


**Supplementary Figure 9. Posterior effect sizes, probabilities of Bayesian Model Averaging model inclusion, and linkage disequilibrium for HLA alleles on 29 different laboratory test phenotypes.** *y*-axis indicates phenotype, and *x*-axis indicates allele. Above - the size of each dot corresponds to the posterior probability that the HLA allele is included as a variable across all plausible models as deemed by BIC measures from BMA, and the color of each dot corresponds to the size and direction of the effect of the allele on the phenotype as found by PLINK. Only the top 10 significant PLINK hits per phenotype were considered for the analysis. Below - LD measures (as determined and visualized by the *gaston* package) across HLA allelotypes; the measures displayed are $R^2$ values.

**Supplementary Table 5. Association results for protein-truncating variants across the 38 lab phenotypes (p < 1e-7).** Protein-truncating variant annotation (variant, ID) and its association to the lab phenotype (trait). Effect size allele (A1), estimated effect size (BETA), standard error (SE), p-value of association (P), minor allele frequency (MAF), predicted protein-truncating or protein-altering variant (Csq), predicted major consequence (Consequence), the HGVS protein sequence name (HGVSp), Gene

Symbol (Gene Symbol), Ensembl Gene ID (Gene), whether the variant is outside of MHC region (is_outside_of_MHC), whether the variant is LD independent based on LD pruning (ld_indep), absolute value of estimated effect size is greater than or equal to .1 (outlier), additional manual comments from authors (Comments).

**Supplementary Table 6. Association results for protein-altering variants across the 38 lab phenotypes (p < 1e-7).** Protein-altering variant annotation (variant, ID) and its association to the lab phenotype (trait). Effect size allele (A1), estimated effect size (BETA), standard error (SE), p-value of association (P), minor allele frequency (MAF), predicted protein-truncating or protein-altering variant (Csq), predicted major consequence (Consequence), the HGVS protein sequence name (HGVSp), Gene Symbol (Gene Symbol), Ensembl Gene ID (Gene), absolute value of estimated effect size is greater than or equal to .1 (outlier), additional manual comments from authors (Comments).

**Supplementary Table 7. Association results for non-coding variants across the 38 lab phenotypes (p < 5e-8).** The non-coding variants characterized on the imputed 1000 Genomes Phase I variants (ID, variant), their positions in centimorgans, (CM) and its association to the lab phenotype (trait). Effect size allele (A1), estimated effect size (BETA), standard error (SE), p-value of association (P), minor allele frequency (MAF), whether the variant is outside of MHC region (is_outside_of_MHC), gene symbol (Gene Symbol), and absolute value of estimated effect size deviates from the standard deviation range estimated from linear fit between log minor allele frequency and absolute value of estimated effect size (outlier, see methods for more details), additional manual comments from authors (Comments).

**Supplementary Table 8. (a) HLA alleles found to be associated to the 38 lab phenotypes via both PLINK association tests and Bayesian Model Averaging (BMA). (b) Other, non-lab phenotypes significantly associated (via PLINK and BMA) to the 37 alleles that had significant results in (a).** Tables enumerate associations' BETA, SE, T/Z STAT values (depending on the type of test), P values from PLINK , and the same P values that have been Benjamini-Yekutieli adjusted (BY_ADJ_P). The tables also contain probabilities of model inclusion from BMA. The tables only enumerate those associations that were found to have both a PLINK association p-value <= 0.05/10000 and a BMA posterior probability >= 0.8.

#### CNVs influencing lab phenotypes

**Supplementary Table 9. Copy number variation associated to the 38 lab phenotypes.** Bonferroni p < 0.05/10000. Columns in the provided data file correspond to the phenotype, chromosome and centroid position of each CNV tested, CNV ID (formatted as chrom:bp1-bp2_del/dup (del denoted by - and dup by +), reference copy number (always N), alternate CNV (always denoted by +), tested "allele" (usually +), genotype model (ADD is additive), N, estimated beta/log odds ratio, standard error of estimate, t/z-statistic, and p-value.

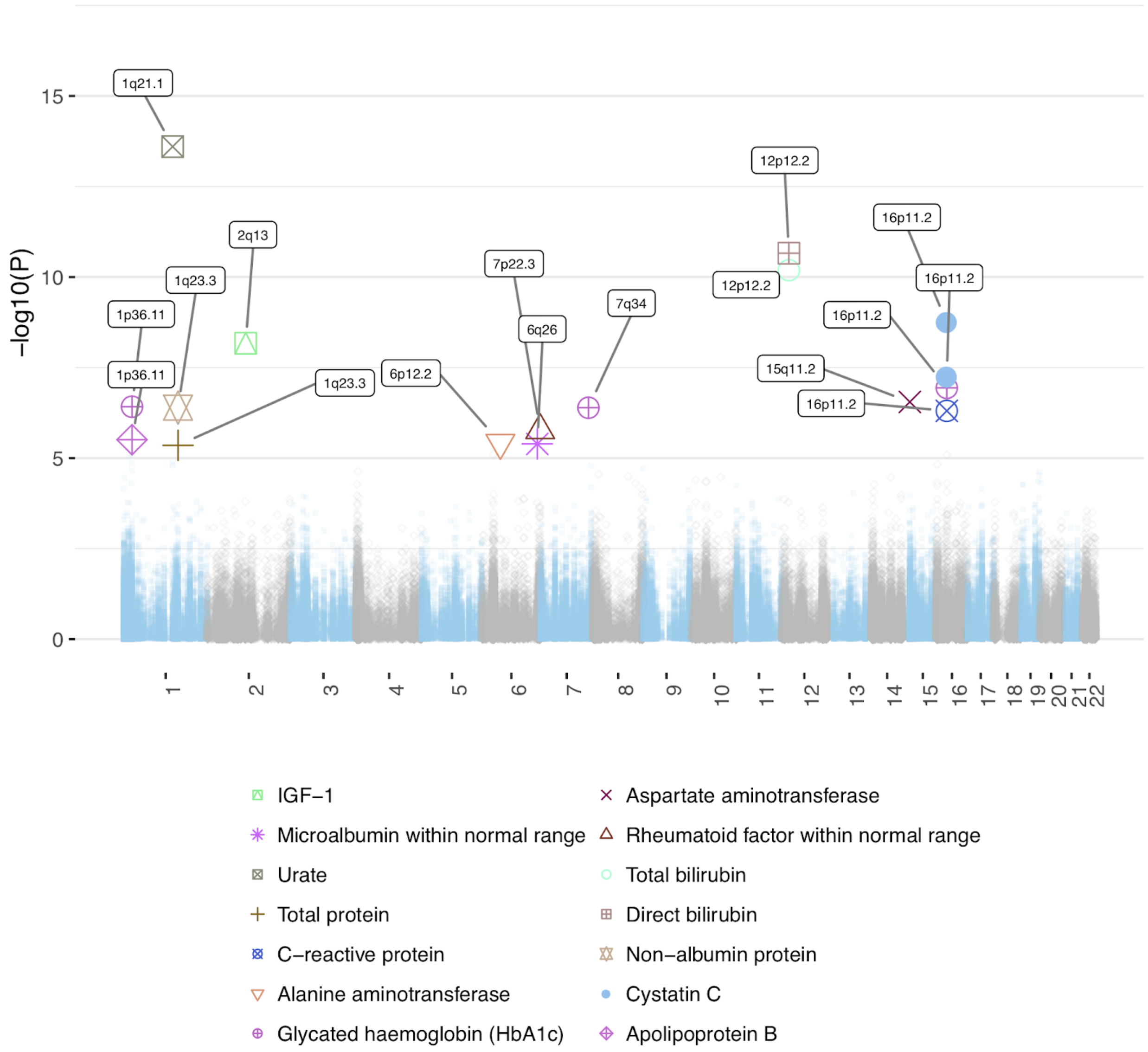


**Supplementary Figure 10A. CNV association analysis across the 38 laboratory tests.** X-axis Genomic coordinate and -log10(P) for single CNV association. CNV laboratory test associations are highlighted when p < .05/10000, with cytogenic band labelled.

**Supplementary Table 10. Rare variant CNV test.** Bonferroni p < .01/25000. Columns in the provided data file correspond to the phenotype, chromosome and centroid position of each gene tested, gene name, reference copy number (always N), burden of CNV (always denoted by +), tested “allele” (usually +), genotype model (ADD is additive), N, estimated beta/log odds ratio, standard error of estimate, t/z-statistic, and p-value.

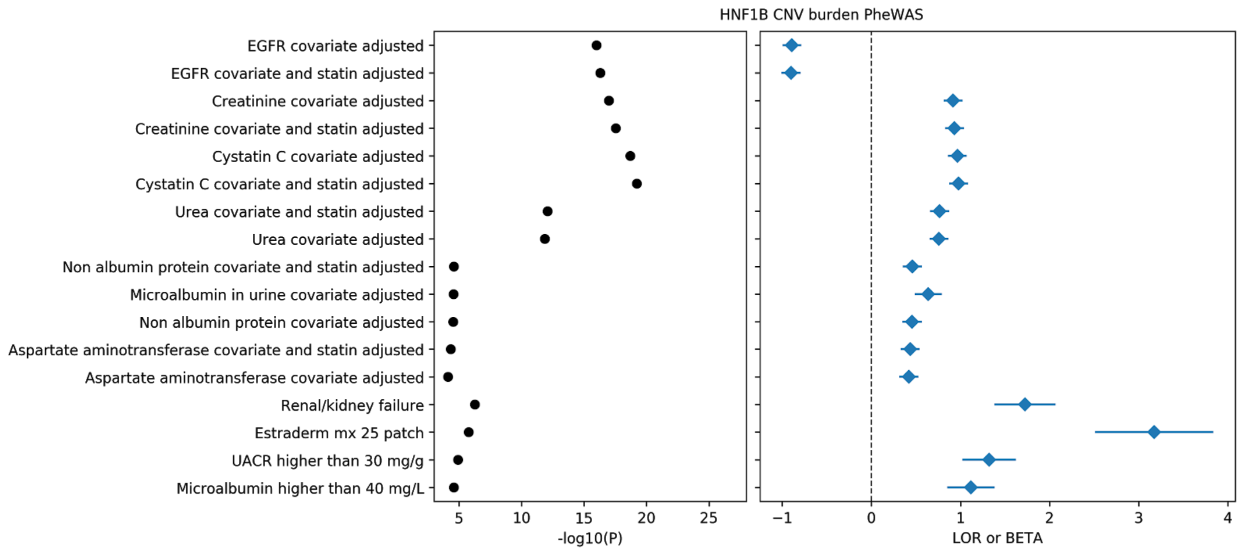


**Supplementary Figure 10B. PheWAS of rare CNVs affecting *HNF1B*.** X-axis log-odds ratio and -log10(P) for each trait having association with *HNF1B* CNVs at p < 1e-4. Associations for all traits run as in previous analysis[3].

#### Global and local heritability of biomarkers

**Supplementary Table 11A. Total SNP heritability.** Total heritability estimates of the 38 laboratory tests with and without adjustment for statins. Estimates are generated using a local SNP heritability model, HESS (Shi et al. 2016).

**Supplementary Table 11B. Cumulative heritability.** Percent heritability explained by the top 1% of SNPs for each of the 38 lab phenotypes.

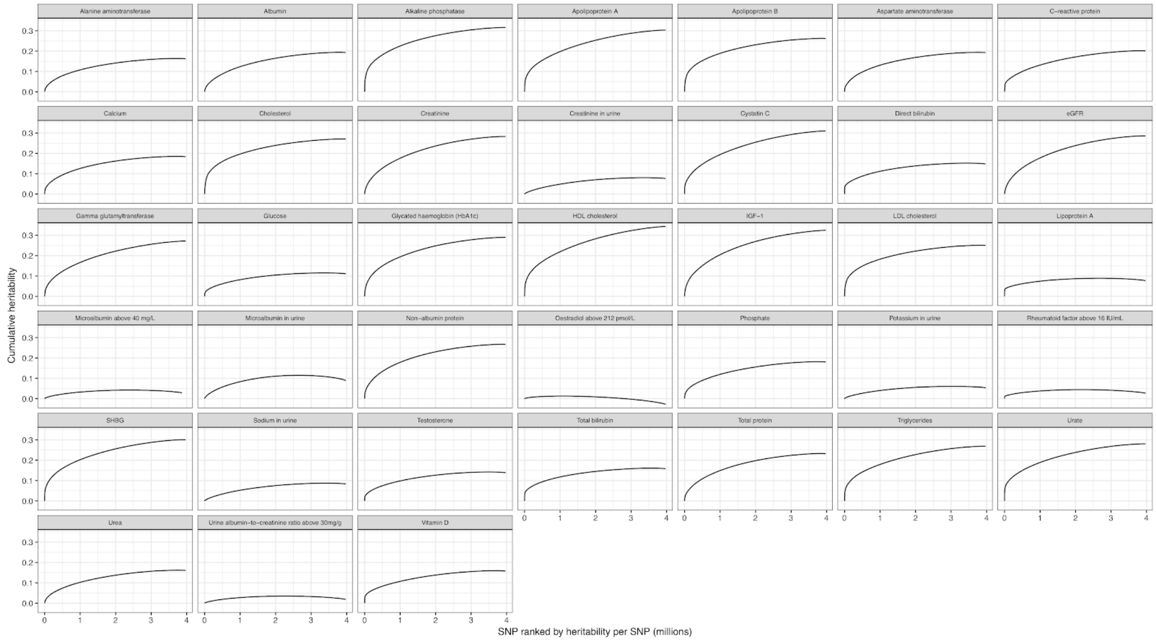


**Supplementary Figure 11. Cumulative heritability.** x-axis SNP ranked by heritability per SNP (millions) and its corresponding cumulative heritability (y-axis) across the 38 lab phenotypes. Lab phenotype label shown in the title of the subplots.

### Cell type decomposition of genetic effects

**Supplementary Table 11C. Cell type enrichments of SNP heritability.** For each of the 10 cell groupings presented previously[4], estimates of the mean and standard deviation of chromatin enrichment for the given cell group were generated using LD Score regression, adjusted for the 53 baseline annotations and the aggregate of each Roadmap annotation mark across all cell types[5].

**Supplementary Table 11D. Single cell heritability enrichments.** Marker genes from single cell RNA-seq datasets from human liver[6], mouse adult kidney[7], human fetal kidney[8], human adult kidney[9], and human adult pancreas[10] were tested for enrichment using S-LDSC[5] with the 53 baseline annotations, Roadmap average annotations, and 10 cell group annotations as controls. Each marker gene set was extended by 100Kb and SNPs the region were annotated as being associated with the cell type.

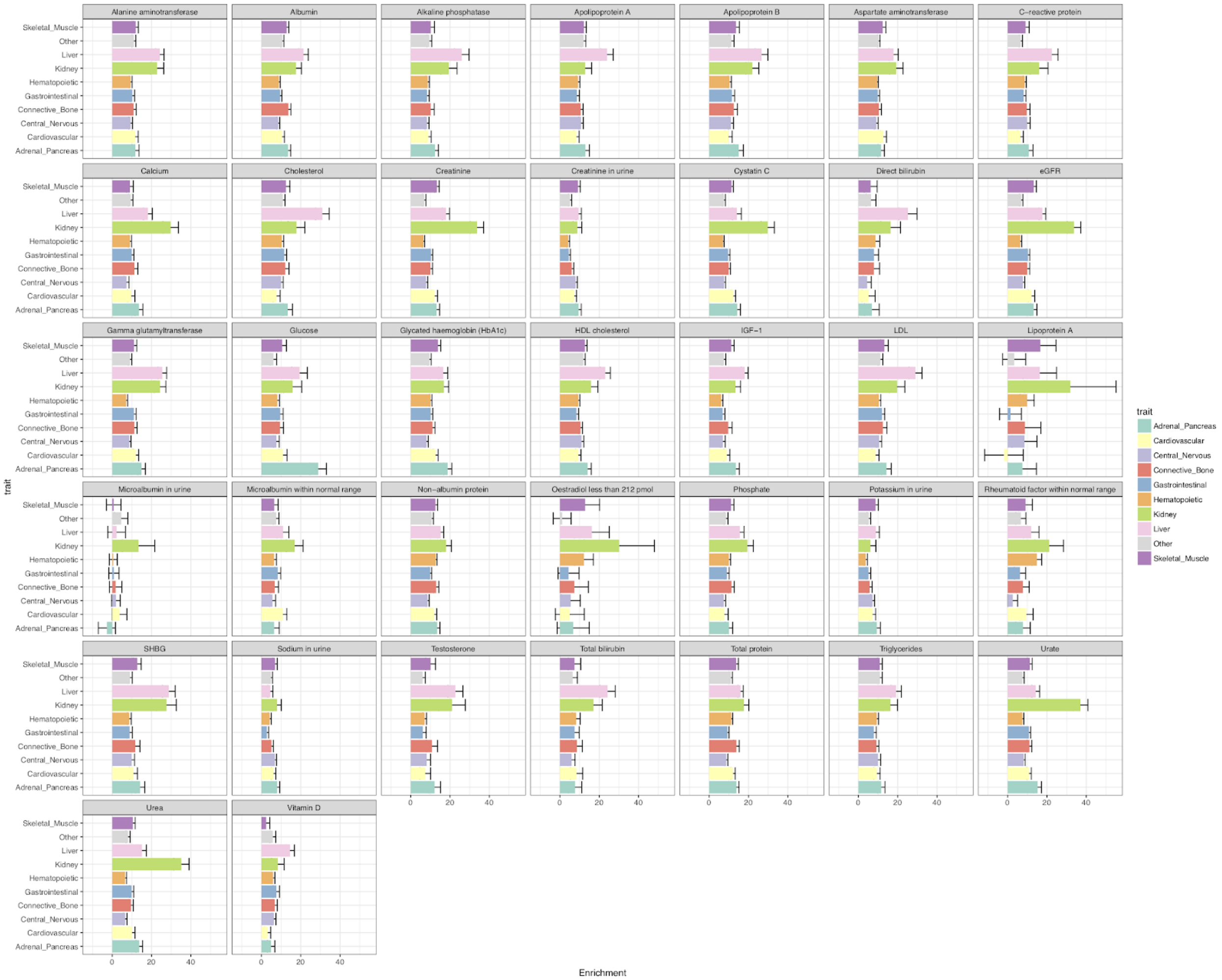

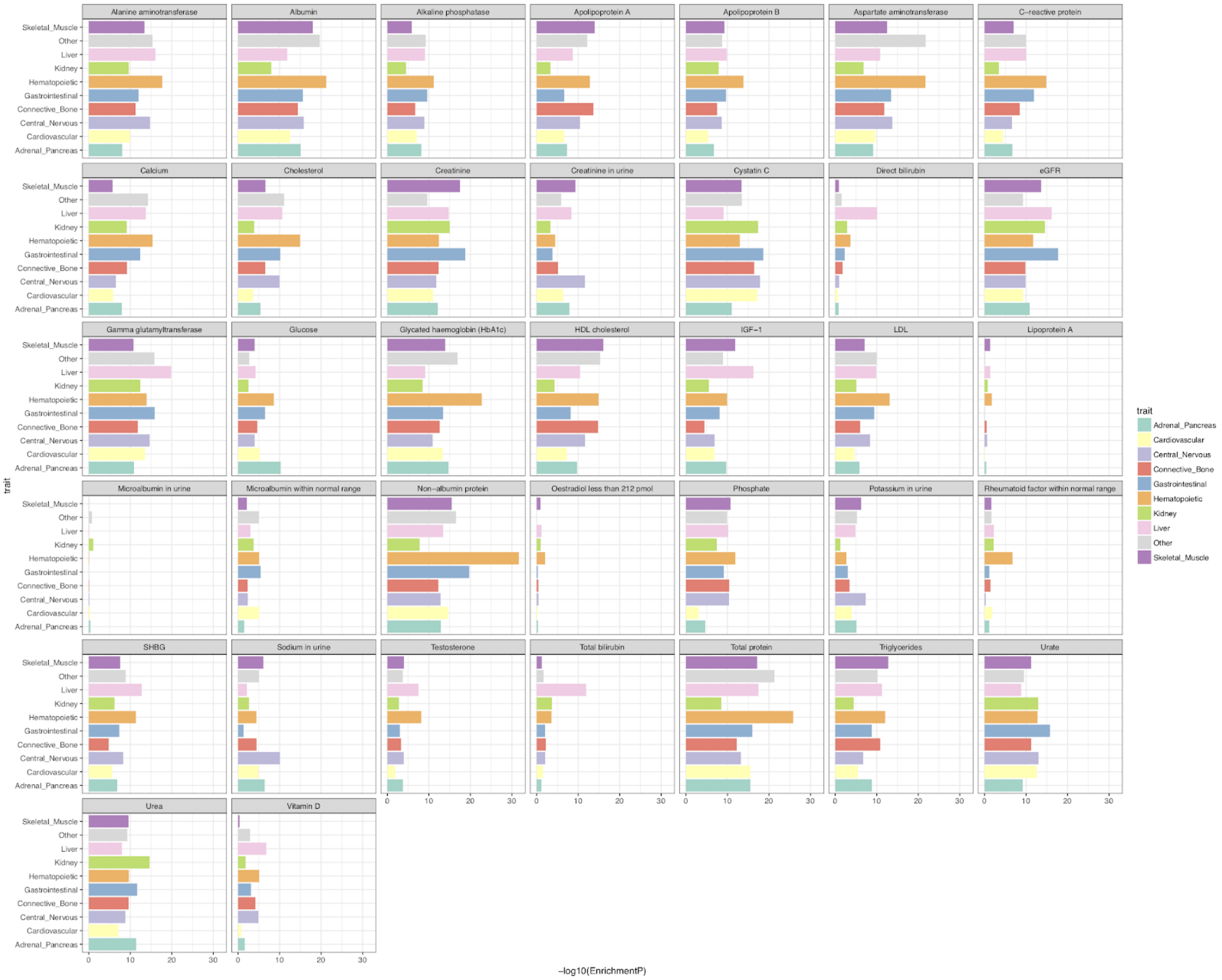


**Supplementary Figure 12. Grouped cell type heritability enrichments across ten tissues.** (x-axis, top) Fold enrichment with SE for each lab phenotype across 10 tissues (y-axis). (x-axis, bottom) -log10(P) value of enrichment or each lab phenotype across 10 tissues (y-axis).

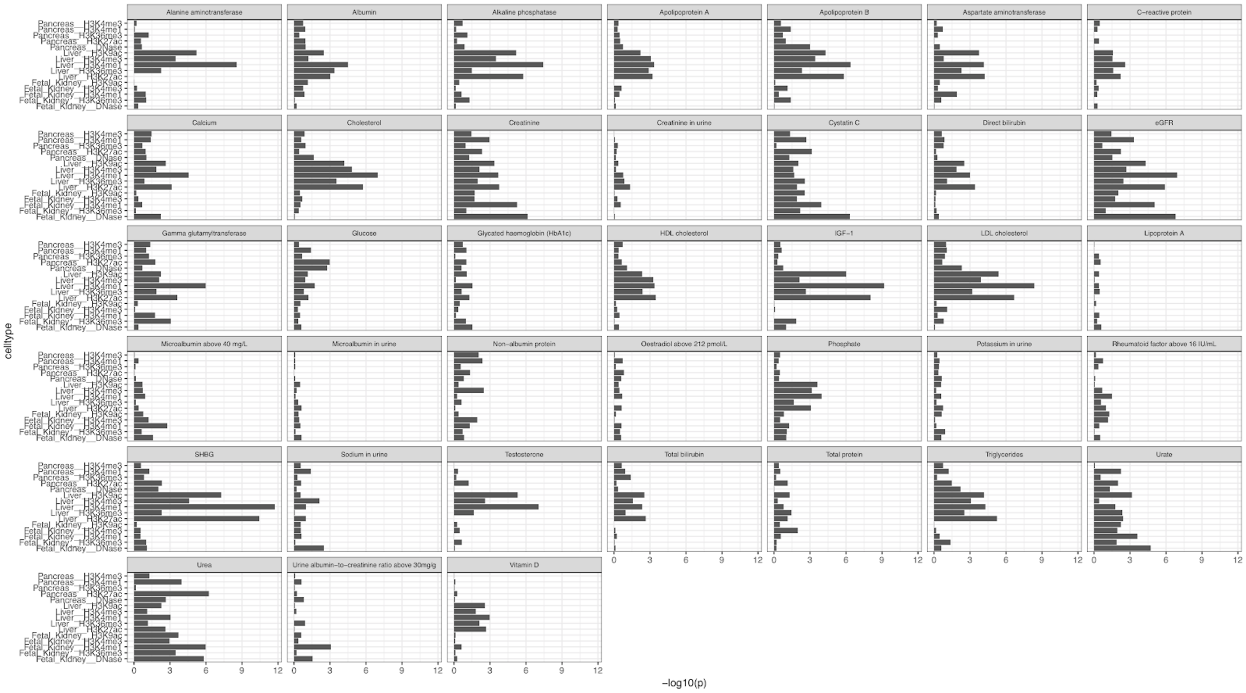


**Supplementary Figure 13. Individual annotations for pancreas, liver, and kidney ChIP-seq experiments.** -log10(P) (x-axis) for cell type heritability enrichment across pancreas, liver, and kidney ChIP-seq experiments (y-axis).

### Targeted phenome-wide association study

**Supplementary Table 12. Association results for the targeted phenome-wide association study (p < 1e-5).** The variant and their ID (Variant, Variant_ID) and its association to disease outcomes (Phenotype) with the corresponding Global Biobank Engine phenotype ID (GBE_ID). The -log10 p-value of association (log10P), estimated effect size (log odds ratio, LOR), standard error of effect size estimate (SE), Gene Symbol (Gene_symbol), predicted protein-truncating or protein-altering variant (Csq), predicted major consequence (Consequence), whether the variant is outside of MHC region (is_outside_of_MHC), whether the variant is LD independent based on LD pruning (ld_indep), and the URLs for the corresponding pages on Global Biobank Engine (GBE_variant_page and GBE_phenotype_page).

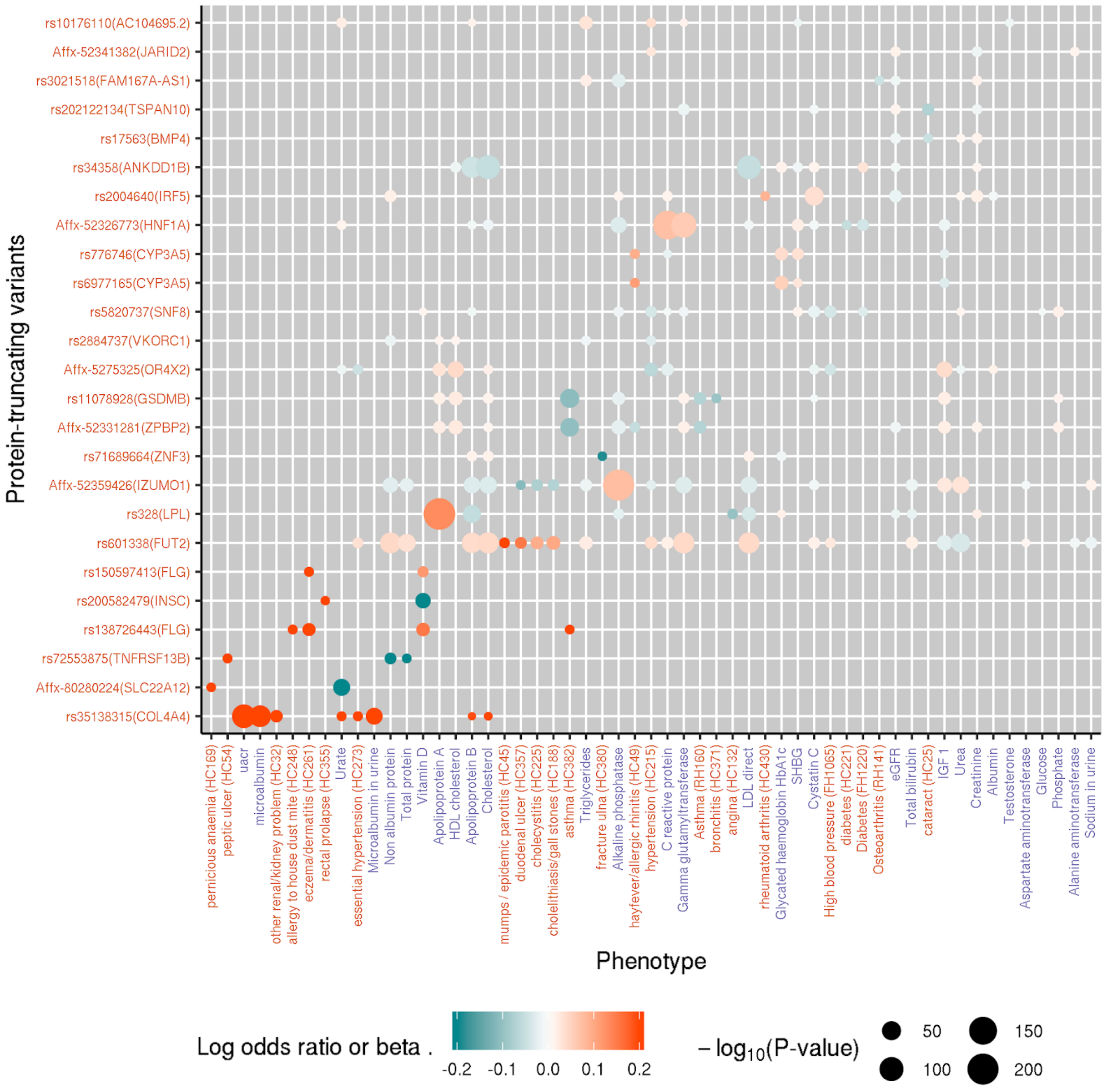


**Supplementary Figure 14. Phenome-wide associations across 25 protein-truncating variants and laboratory measurements and 24 disease outcomes in the UK Biobank.** Targeted phenome-wide association analysis was performed for PTVs outside of the human MHC region that showed significant genome-wide associations (p < 1e-7) with at least one of the laboratory measurement traits. The log odds ratio of the significant PheWAS associations (p < 1e-5) are shown across phenotypes (x-axis) and PTVs (y-axis). The 46 significant (p < 1e-5) associations across 25 variants and 24 disease outcomes are shown as well as the associations with laboratory measurements. The color of phenotype names indicate binary disease outcomes or family history (red) or laboratory measurements (purple). The color for log odds ratio or beta = 0.2 is used for the associations with > 0.2 log odds ratio or beta.

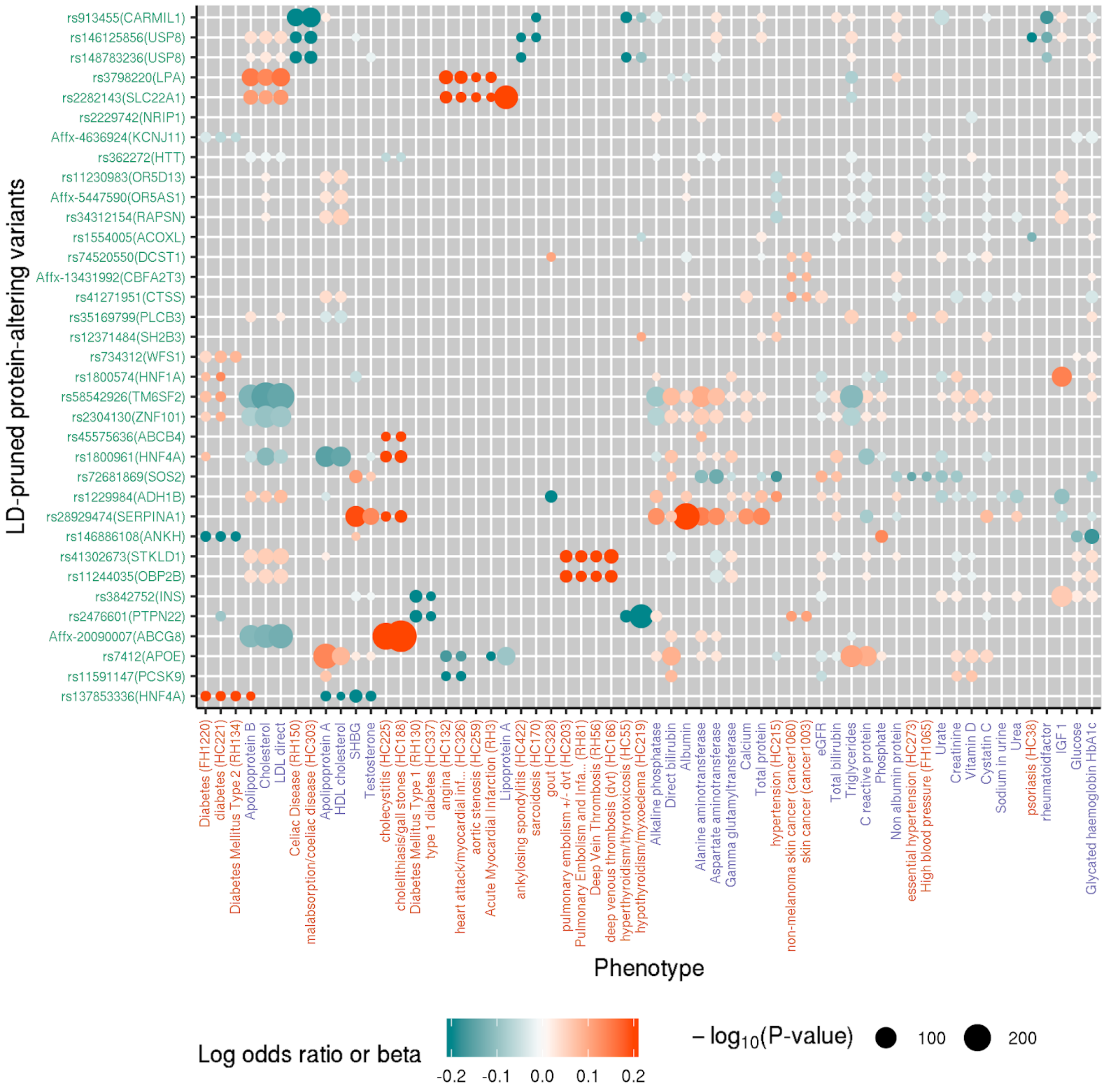


**Supplementary Figure 15. Phenome-wide associations across 35 LD-independent protein-altering variants and 28 disease outcomes in the UK Biobank.** Targeted phenome-wide association analysis was performed for protein-altering variants outside of the human MHC region that showed significant genome-wide associations ($p < 1e-7$) with at least one of the laboratory measurement traits. The log odds ratio of the significant PheWAS associations ($p < 1e-5$) are shown across phenotypes (x-axis) and protein-altering variants (y-axis). Out of 172 significant ($p < 1e-5$) associations across 80 LD-independent protein-altering variants and 75 disease outcomes, 35 variants and 28 disease outcomes with maximal number of significant associations are chosen for visualization. The associations for those variant-phenotype pairs are shown as well as the associations across laboratory measurement phenotypes. The color of phenotype names indicate binary disease outcomes or family history (red) and laboratory measurements (purple). The color for log odds ratio or beta = 0.2 is used for the associations with > 0.2 log odds ratio or beta.

### Correlation of genetic effects between laboratory tests, diseases, and medically relevant phenotypes

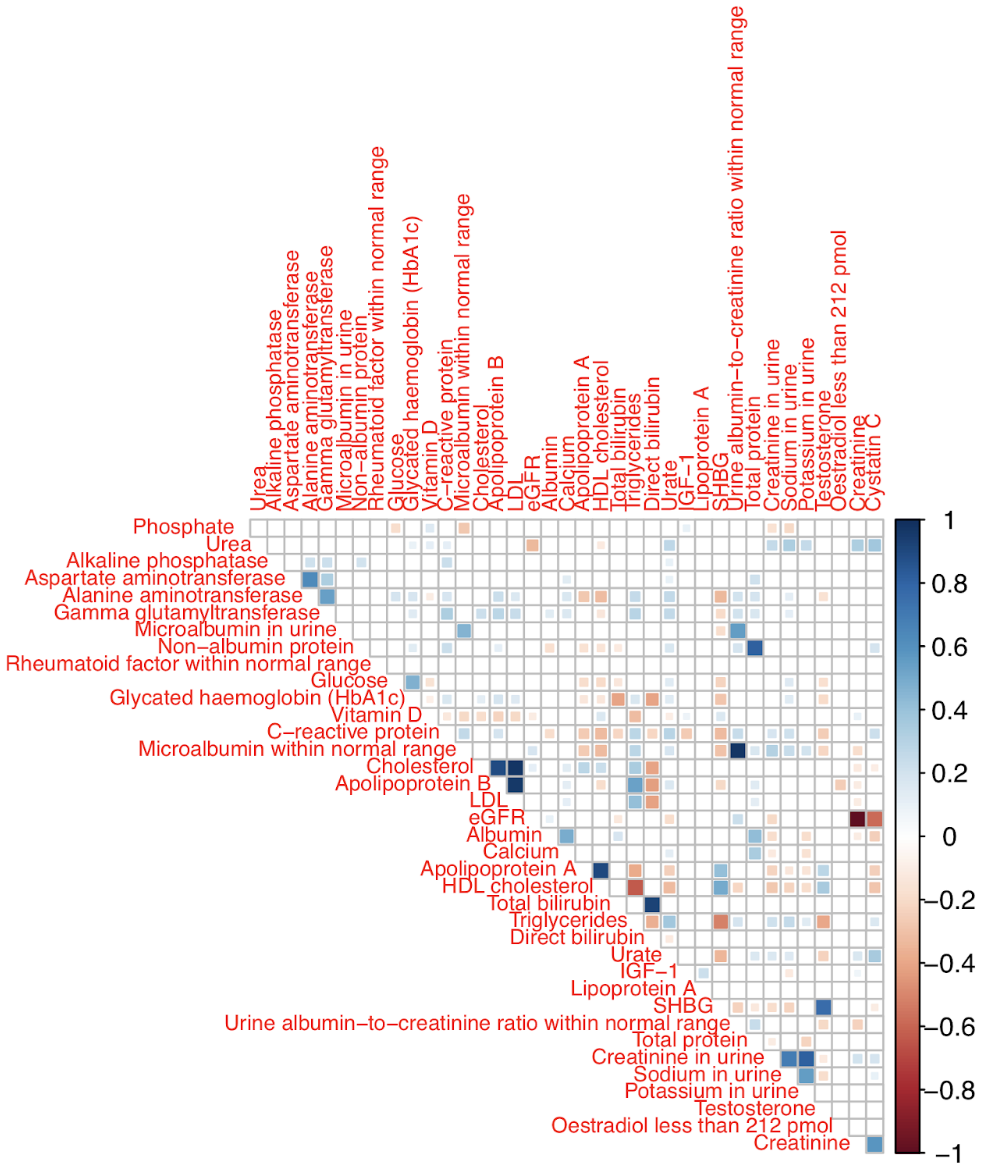


**Supplementary Figure 16. Correlation of genetic effects between laboratory tests.** -1 (red) to 1 (blue) scale of correlation of genetic effects estimated using LD-score regression.

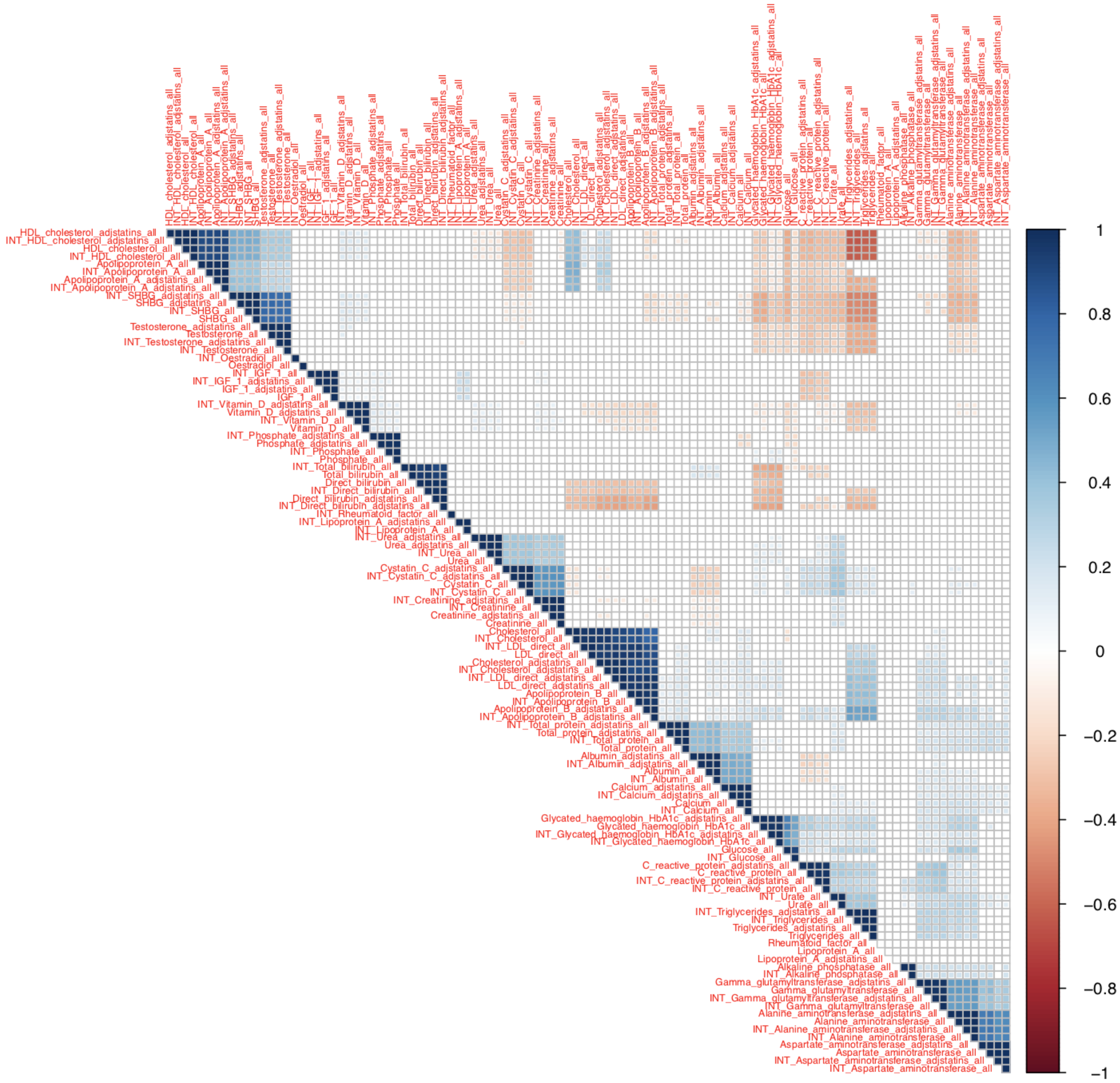


**Supplementary Figure 17. Correlation of genetic effects between laboratory tests with normalization (“INT”), and with lipid-lowering therapy adjustment (“adjstatins”) and without.** -1 (red) to 1 (blue) scale of correlation of genetic effects estimated using LD-score regression.

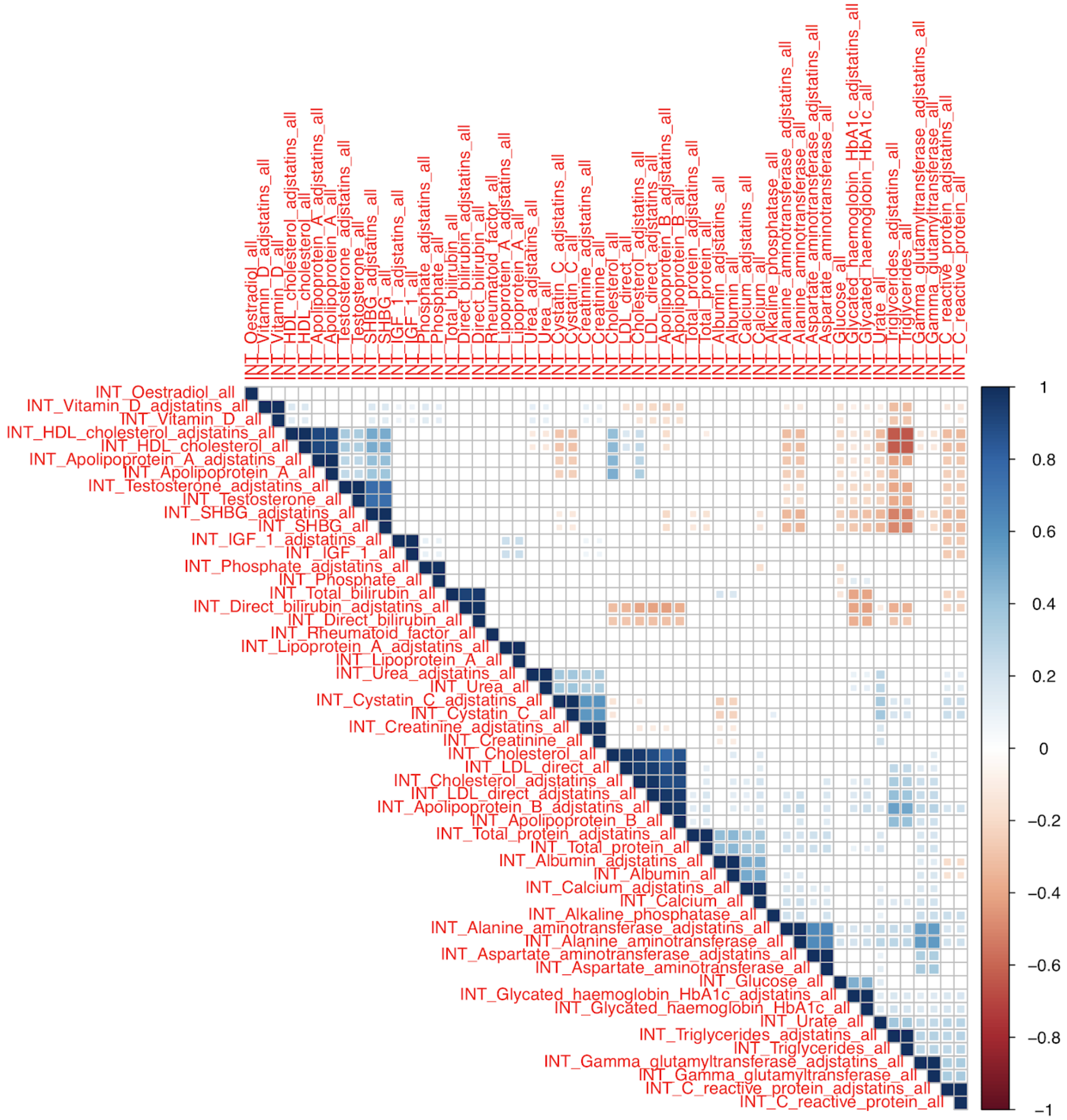

**Supplementary Figure 18. Correlation of genetic effects between normalized ("INT") lab phenotypes with lipid-lowering therapy adjustment ("adjstatins") and without.** -1 (red) to 1 (blue) scale of correlation of genetic effects estimated using LD-score regression.

**Supplementary Table 13. Disease and medically relevant phenotypes used for genetic correlation analysis.**

### Causal inference

**Supplementary Table 14. Causal inference results using MR-Egger and LCV.** Each row represents a significant exposure-outcome pair by either MR-Egger or LCV (FDR 10%). The edge type marks if the causal link was found by MR-Egger only, LCV only, or both. Estimated causal effects are presented for all pairs.

#### Mendelian randomization

Mendelian randomization methods enable estimation of causal effects between an exposure X and an outcome Y. Given a set of genetic instruments of X (i.e., direct causes of X that are not affected by confounders), the causal effect of X on Y can be extracted by analyzing their associations with both X and Y. Most methods are based on linear models and start with a 2D plot of the association summary statistics. A meta-analysis is then used to estimate if there is a significant correlation between the effects, which then translates into a line whose slope reveals the causal effect. MR-Egger is a powerful method that uses Egger regression for the meta-analysis[11]. Egger regression was developed originally for correcting publication bias in meta-analyses, but the problem is analogous to adjusting bias from pleiotropy in the MR setting. Thus, Egger-regression provides a way to both estimate and adjust for biases in the 2D plot that originate from pleiotropic effects (under the assumption that the association of each genetic instrument with the exposure is independent of the pleiotropic effect of the variant).

#### Latent causal variables

LCV is a recent method that makes use of the MR graphical model to evaluate if an observed genetic correlation can be attributed to a causal relationship[12]. LCV is based on a 2D analysis of summary statistics as in MR methods, with two notable differences. First, it uses a latent variable to model the mediation of genetic correlation between two traits. This allows for the estimation of the full or partial proportion of genetic causal relation between two traits. Second, it takes as input all summary statistics and does not require a set of independent instruments. On the other hand,unlike MR methods, LCV does not address reverse causality, and it does not estimate causal effect sizes.

### Polygenic prediction of biomarkers within and across populations

**Supplementary Table 15. Predictive performance of lab phenotypes from genetic data within and across populations.** The laboratory phenotype (Phenotype), whether the phenotype is binary (bin) or quantitative (qt), evaluated population (population), the increments of predictive performance (AUC for binary traits and R for quantitative traits) from covariate-only model to the model with both covariates and genotypes (delta_R_or_AUC), predictive performance measures of the model with genotype and covariates (Genotype_and_covariates), the model with covariates (Covariates_only), and the model with genotypes (Genotype_only), and their trans-populational comparison with respect to self-identified White British population shown in percent (Relative_to_WB_delta_R_or_AUC, Relative_to_WB_Genotype_and_covariates, Relative_to_WB_Covariates_only, and Relative_to_WB_Genotype_only).

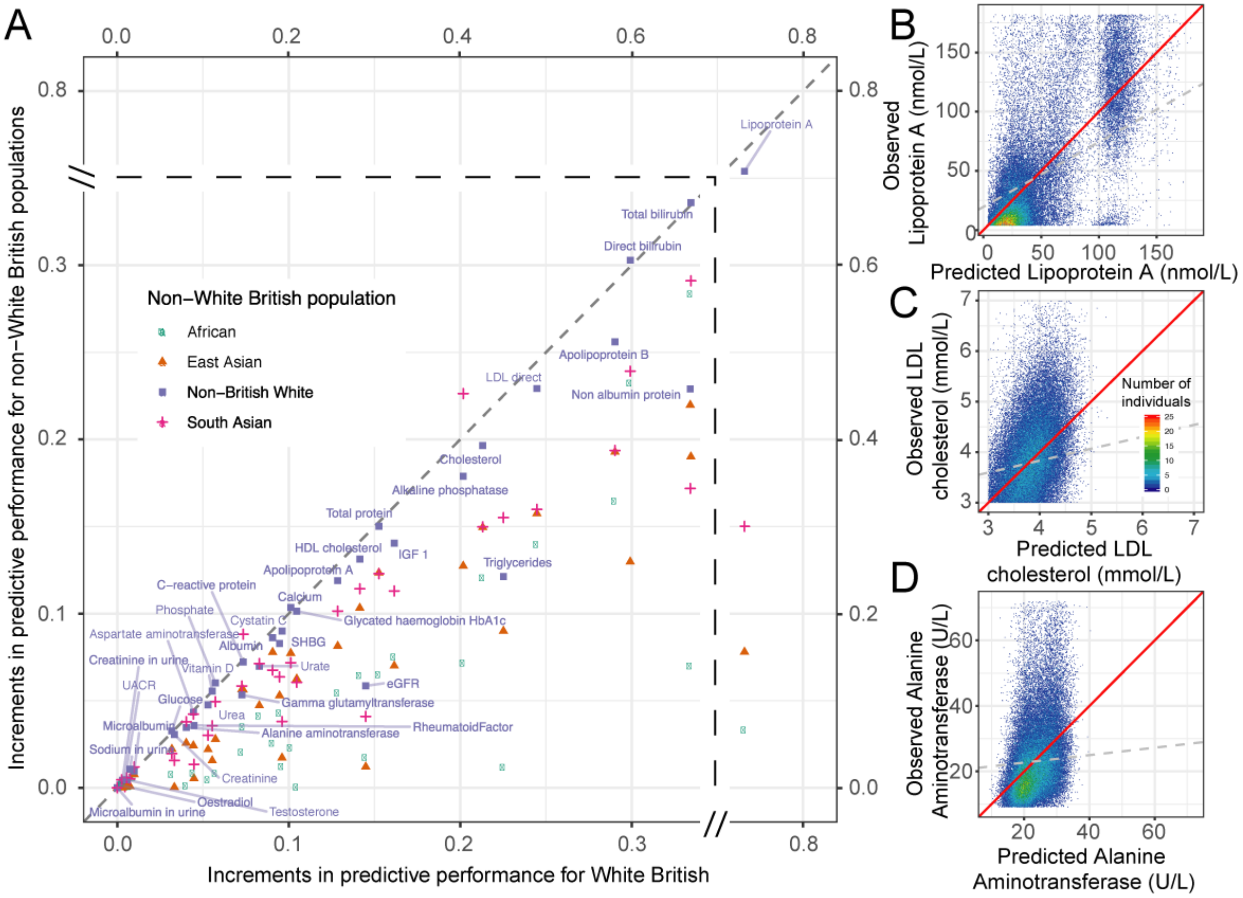


**Supplementary Figure 19A-D. Lab phenotype prediction from genetic data within and across populations. (A)** Increments in predictive performance with genetic data (change in correlation, R, or ROC-AUC) for self-identified White British (x-axis) and other ethnic groups (y-axis) are shown across the 38 lab phenotypes. **(B-D)** Predicted vs. observed phenotypes comparison for individuals in the test sets for Lipoprotein A **(B)**, LDL **(C)**, and alanine aminotransferase **(D)**. The red diagonal line indicates x=y, whereas the gray dashed line shows the linear regression fit between observed and predicted phenotypes.

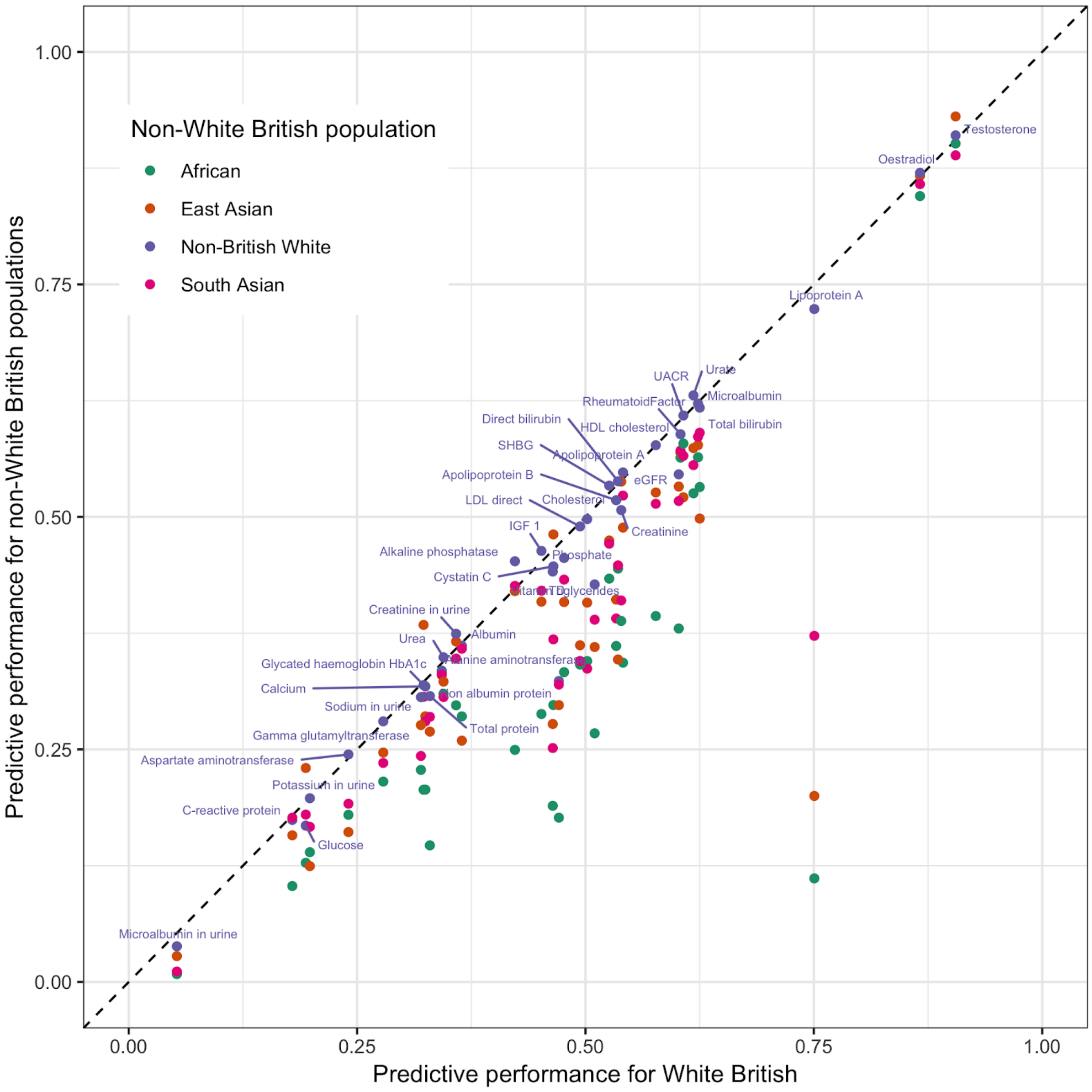


**Supplementary Figure 19E. Lab phenotype prediction from genetic data within and across populations.** The predictive performance with both genetic data and covariates (correlation, R) for self-identified White British (x-axis) and other ethnic groups (y-axis) are shown across the 38 lab phenotypes.

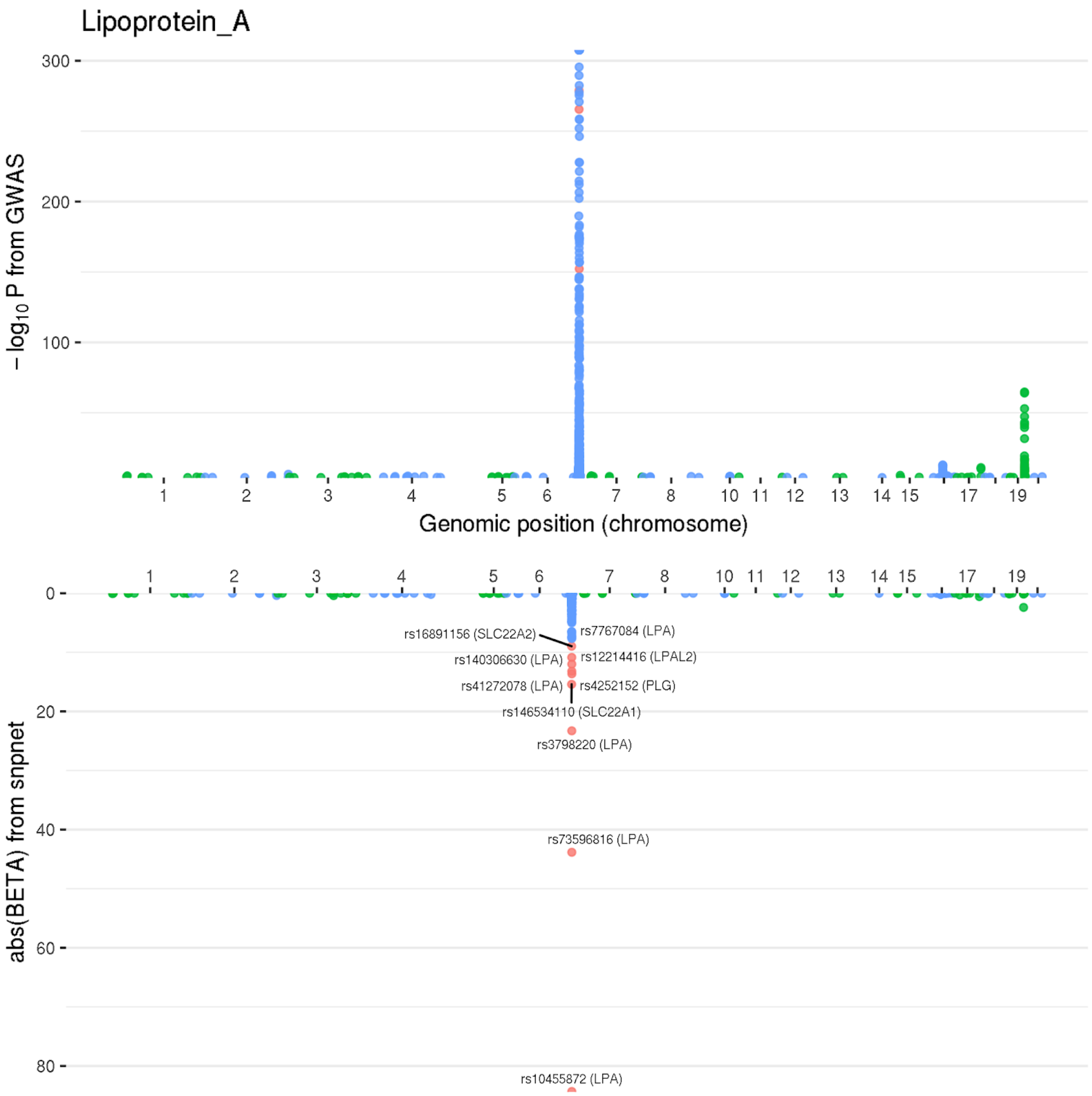


**Supplementary Figure 20A. “Lake” plots of GWAS p-value and the magnitude of effect size estimates from snpnet for Lipoprotein A.** (x-axis) Genomic coordinates for (top panel) -log10(P) from GWAS and (bottom panel) absolute value of estimated
260 effect size using snpnet (abs(BETA) from snpnet).

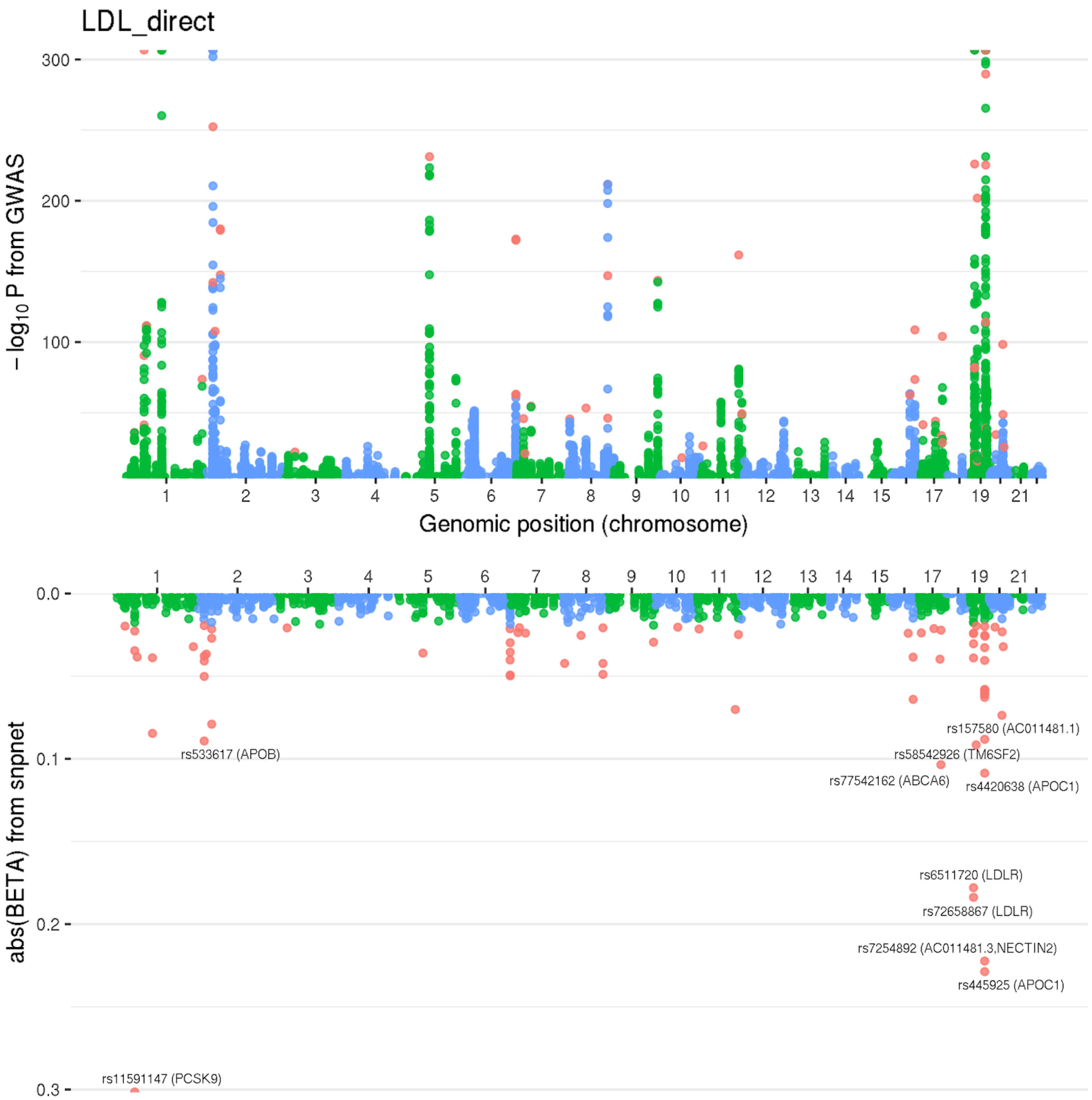


**Supplementary Figure 20B. "Lake" plots of GWAS p-value and the magnitude of effect size estimates from snpnet for LDL.** (x-axis) Genomic coordinates for (top panel) -log10(P) from GWAS and (bottom panel) absolute value of estimated effect size using snpnet (abs(BETA) from snpnet).

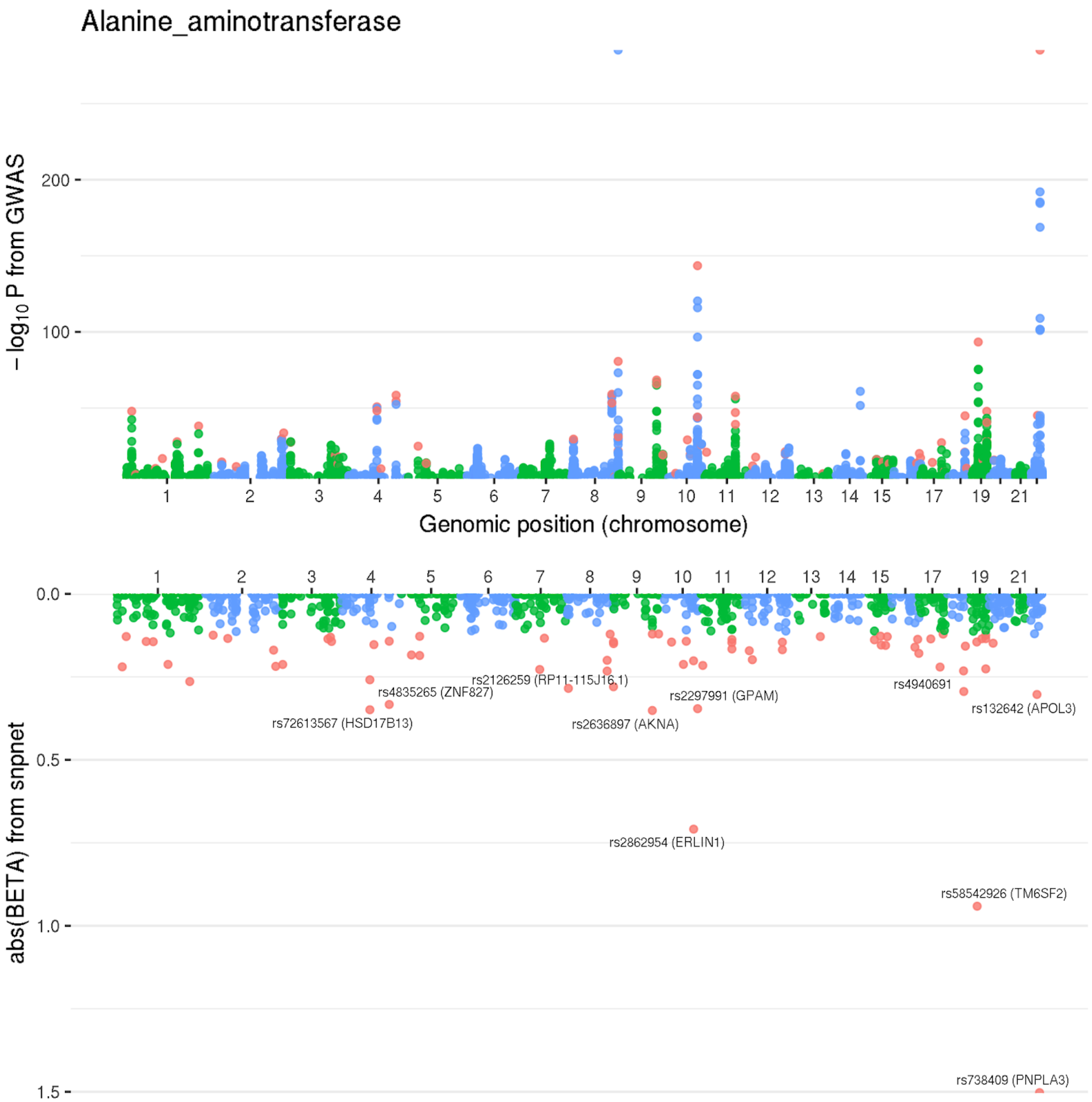


265 **Supplementary Figure 20C. "Lake" plots of GWAS p-value and the magnitude of effect size estimates from snpnet for Alanine Aminotransferase.** (x-axis) Genomic coordinates for (top panel) -log10(P) from GWAS and (bottom panel) absolute value of estimated effect size using snpnet (abs(BETA) from snpnet).

**Supplementary Table 16. Population-specific bias in polygenic prediction of the 38 lab phenotypes**. The rank of the increments in predictive performance comparing the PRS model with both genotype and covariates and covariate alone across 5 population groups are summarized. The sum across population for a given rank varies due to the ties in the ranks.

### Multiple regression with PRSs for laboratory tests improves prediction of traits and diseases

**Supplementary Table 17. Predictive power of multiple regression of laboratory tests.** Each trait is treated independently an a regression model (linear or logistic, determined by outcome) is used. McFadden's adjusted R^2 (for binary outcomes) and Adjusted R^2 (for continuous outcomes) are presented for models which contain just covariates or covariates with the traits of interest. All regressions were run with age, sex, genotyping array, 40 principal components of the genotyping matrix, age squared, townsend deprivation index, and age-sex interaction. Type 2 diabetes additionally had covariates of BMI and Waist to Hip ratio and interactions of each with age and sex, and liver fat percentage has covariates of alcohol and interactions with age and sex.

**Supplementary Table 18. F test for improved predictive performance of liver fat percentage.** F statistics for comparison of the explained variance under the covariate only model versus the trait PRS and combination of all laboratory test PRSs, as well as comparisons of each of these with the combined model with all PRSs**.** We observed a consistent and significant improvement across all model comparisons.

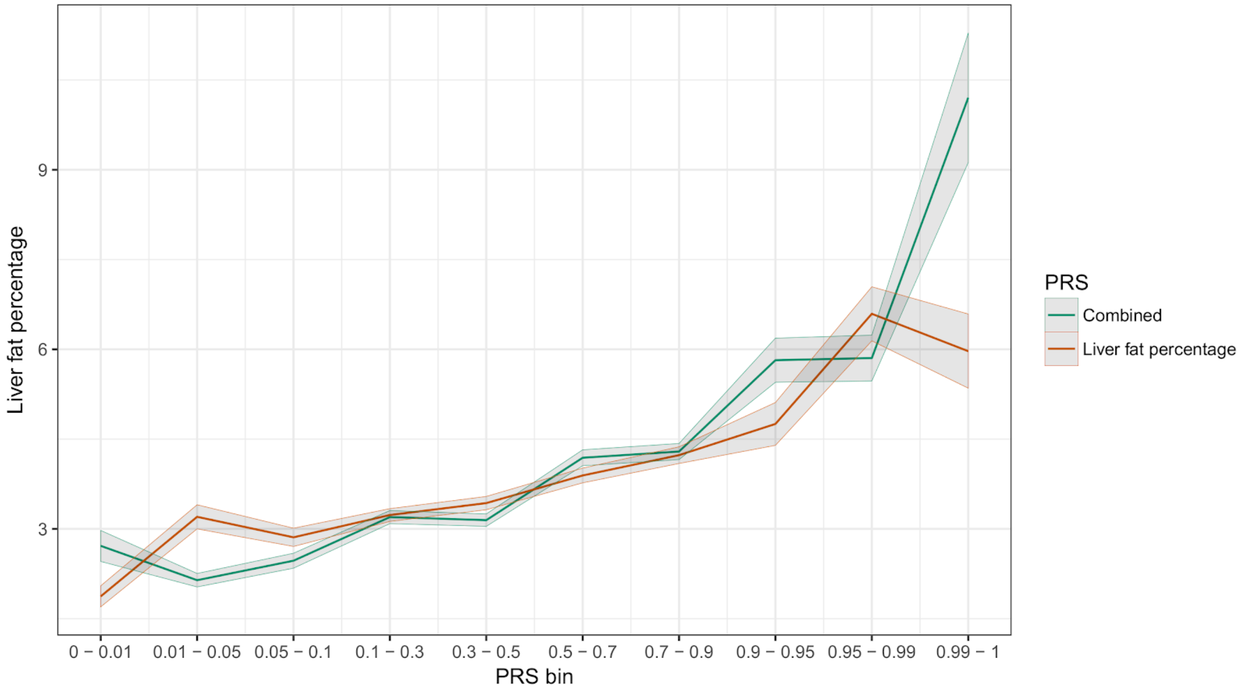


**Supplementary Figure 21a. Average liver fat percentages at different polygenic score quantiles.** Individual estimates were predicted from existing polygenic scores and the bins of self-identified White British individuals in both the snpnet PRS for liver fat percentage and the multi-PRS are shown. Estimates are averages of the actual percentage of liver fat, determined by MRI.

**Supplementary Table 19. Regression coefficients for prediction of liver fat percentage.** Regression coefficient terms and their standard errors estimated from individual liver fat percentage. All terms included in the full regression model are present in the table.

**Supplementary Table 20. F test for improved predictive performance of other traits.** F statistics for comparison of the explained variance under the covariate only model versus the trait PRS and combination of all laboratory test PRSs, as well as comparisons of each of these with the combined model with all PRSs, for each of kidney and liver cancer, and acute and all myocardial infarction. All results suggest significant improvement of laboratory tests versus trait PRSs alone.

**Supplementary Table 21. Prevalence estimates within quantiles of polygenic scores of the multi-PRS.** Each quantile is estimated with the number of individuals and corresponding cases in the population indicated.

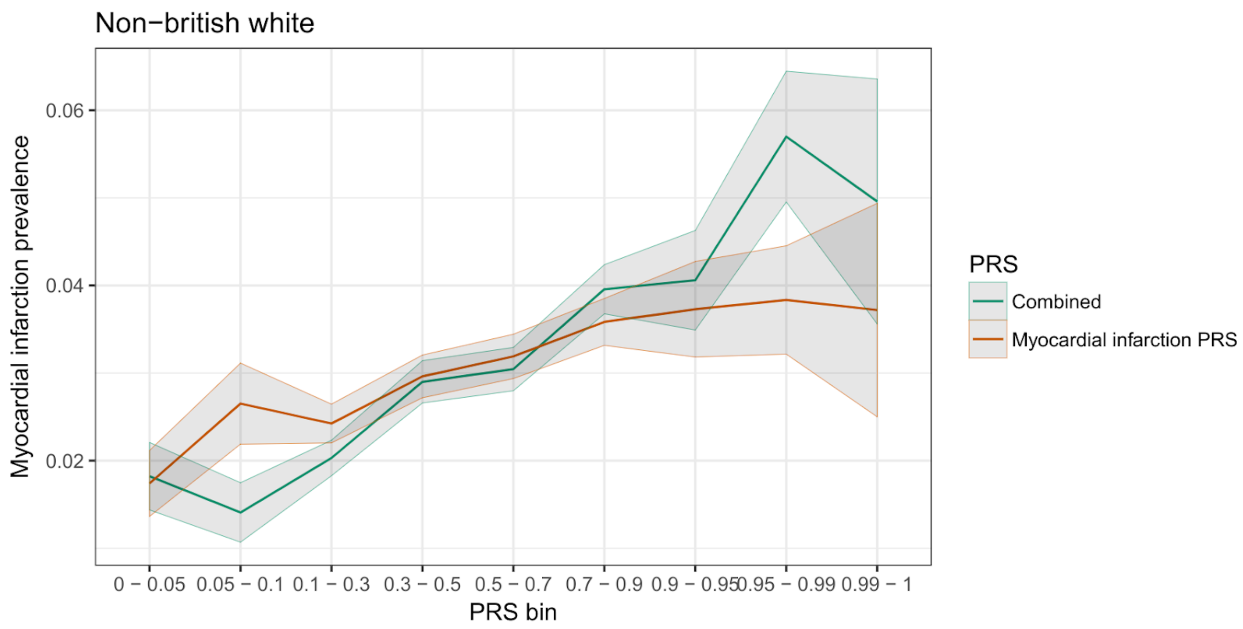


**Supplementary Figure 21b. Prevalence of myocardial infarction in quantiles of the self-identified non-British White individuals in UK Biobank.** Individual estimates were predicted from existing polygenic scores and the bins of individuals in both
295 the snpnet PRS for myocardial infarction and the multi-PRS are shown. Quantiles were chosen to ensure an adequate number of cases in each quantile bin.

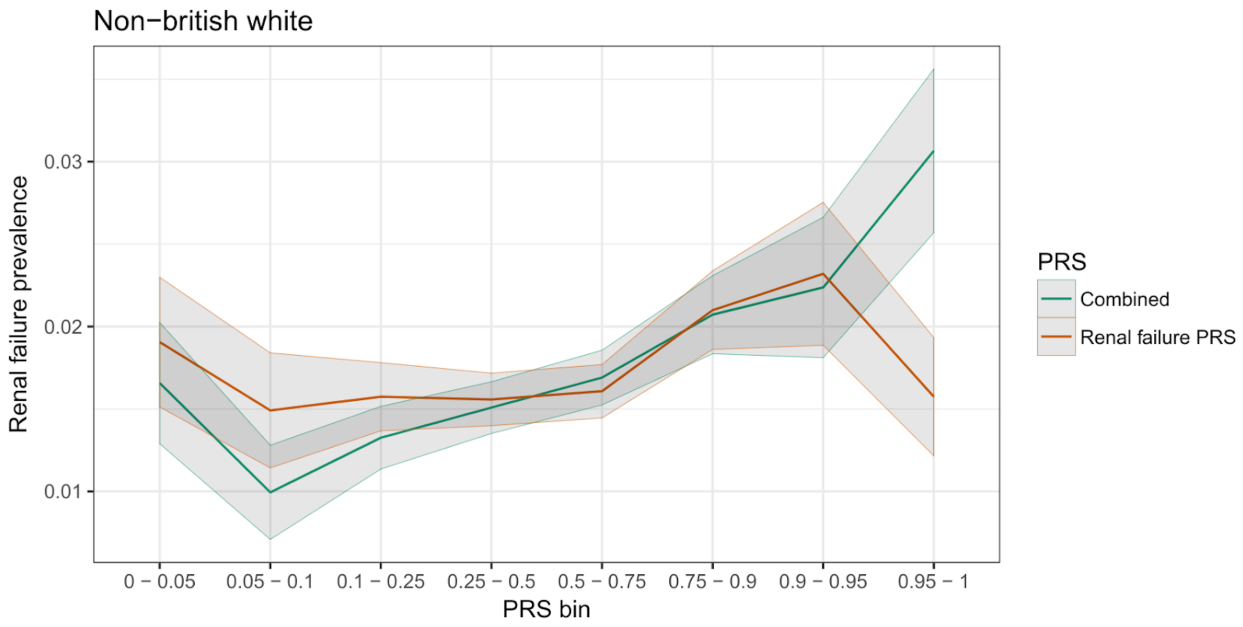


**Supplementary Figure 22a. Prevalence of renal failure in quantiles of the self-identified non-British White individuals in UK Biobank.** Individual estimates were predicted from existing polygenic scores and the bins of individuals in both the snpnet PRS for renal failure and the multi-PRS are shown. Quantiles were chosen to ensure an adequate number of cases in each quantile bin.

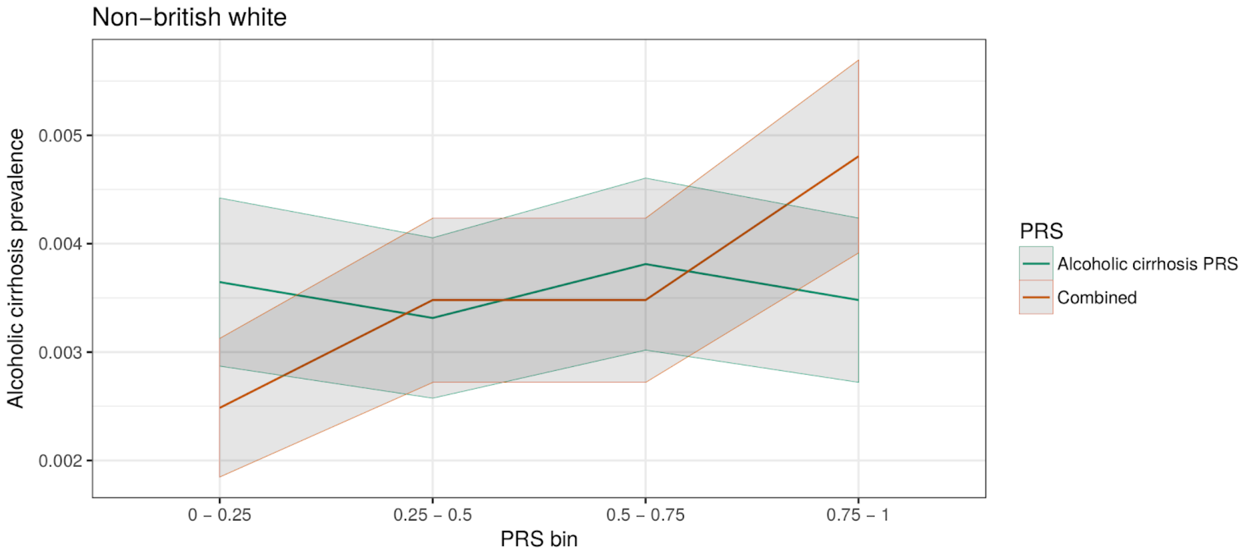


**Supplementary Figure 22b. Prevalence of alcoholic cirrhosis in quantiles of the self-identified non-British White individuals in UK Biobank.** Individual estimates were predicted from existing polygenic scores and the bins of individuals in both the snpnet PRS for alcoholic cirrhosis and the multi-PRS are shown. Quantiles were chosen to ensure an adequate number of cases in each quantile bin.

**Supplementary Table 22. Description of case definition in FinnGen derived from ICD codes and registry data.** All ICD-8, ICD-9, ICD-10, cause of death, and medical reimbursement codes used to identify cases for each trait are shown.

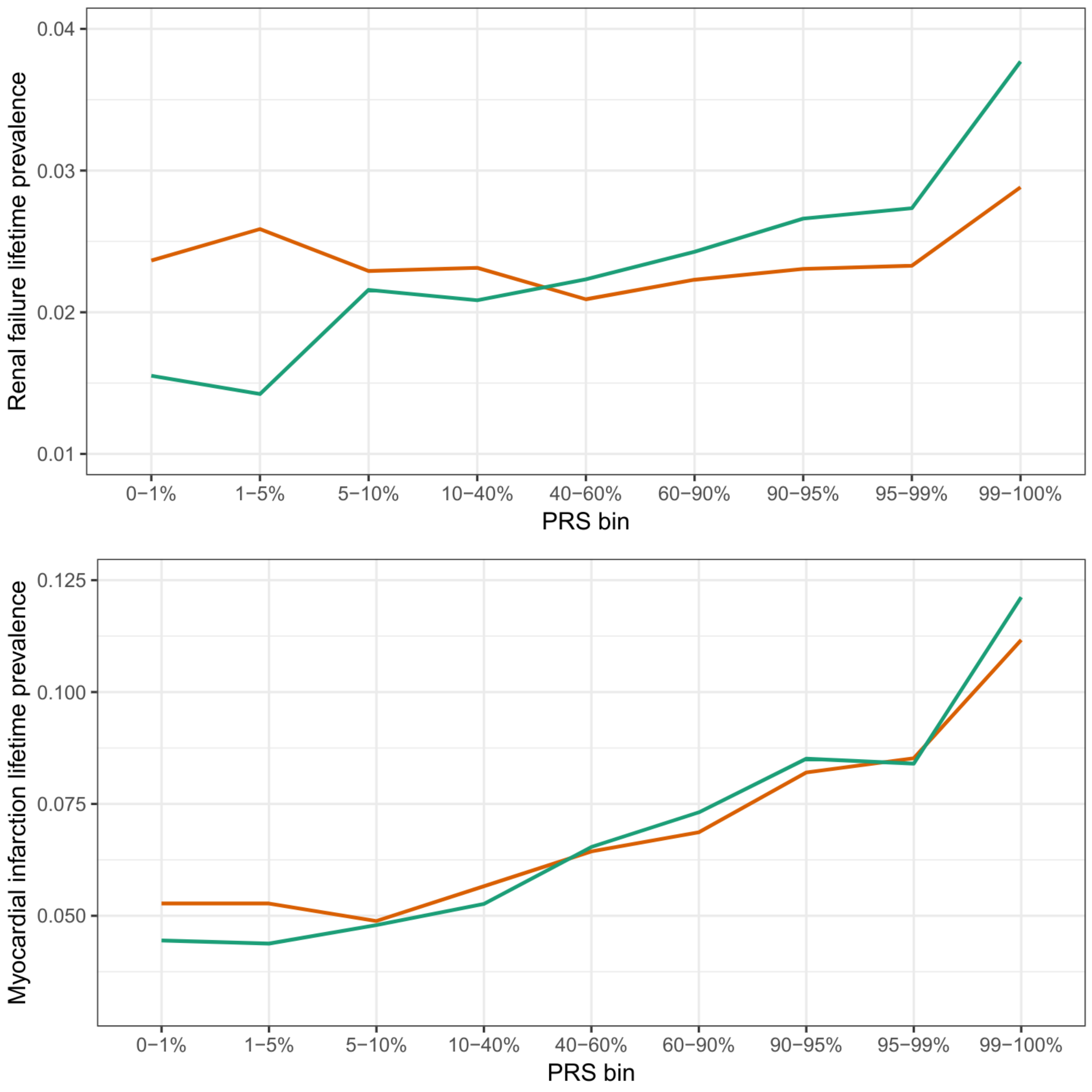


**Supplementary Figure 23. Prevalence of renal failure and myocardial infarction in quantiles of Finnish individuals in FinnGen.** Individual estimates were predicted from existing polygenic scores and the bins of individuals in both the snpnet PRS for renal failure and the multi-PRS are shown.

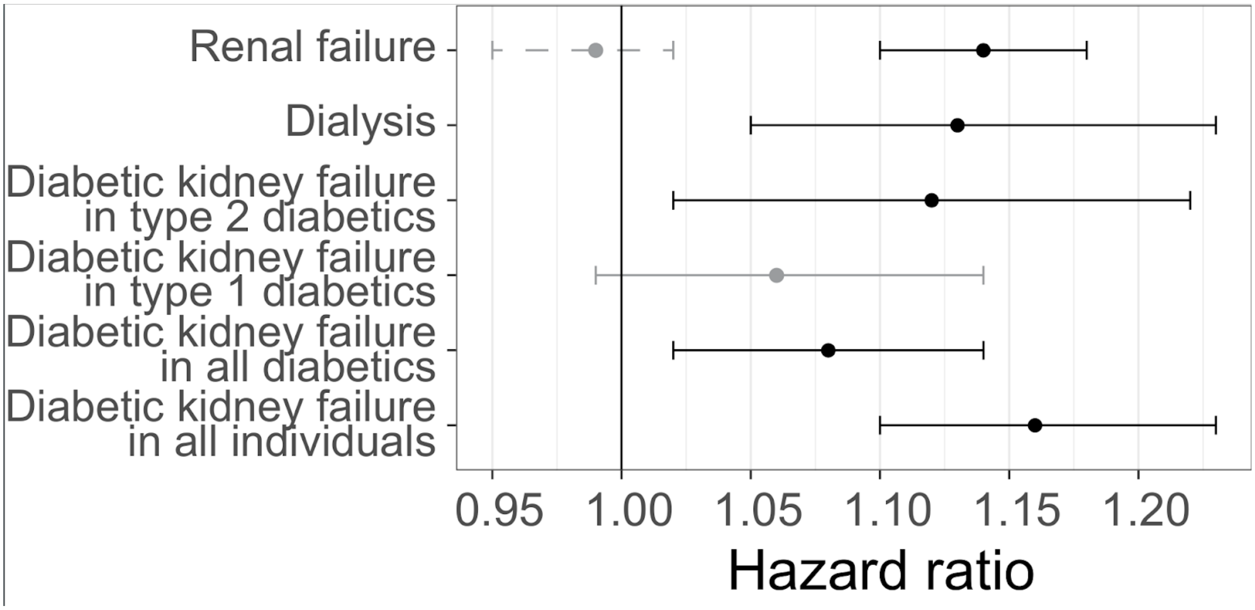


**Supplementary Figure 24. Extended diabetes status predictions of the renal failure multi-PRS.** Hazard ratios for incidence of various outcomes using the renal failure PRS (solid lines) or snpnet PRS for renal failure (dashed line). Error bars represent 95% confidence intervals. Number of individuals with each diagnosis, statistical significance, and covariates described in Supplementary Table 23 and Methods.

**Supplementary Table 23. Estimated predictive ability in incident cases in FinnGen.** Estimated hazard ratios on incident cases of each PRS are presented, with the number of individuals shown developing disease during the follow-up period. Results are presented for both the trait PRS (trained with snpnet) or the combined multi-PRS model; the hazard ratio, standard errors, and overall predictive ability represented as a C-index are shown.

**Supplementary Table 24. Estimated predictive ability in incident cases in FinnGen at different PRS bins.** The number of individuals and cases in each bin, as well as the corresponding hazard ratios, for both the trait PRS and multi-PRS are shown.

#### FinnGen

FinnGen consists of the following people:

Steering Committee

| | |
|---|---|
| Aarno Palotie | University of Helsinki / FIMM |
| Mark Daly | University of Helsinki / FIMM |

Pharma

| | |
|---|---|
| Howard Jacob | Abbvie |
| Athena Matakidou | Astra Zeneca |
| Heiko Runz | Biogen |
| Sally John | Biogen |
| Robert Plenge | Celgene |
| Julie Hunkapiller | Genentech |
| Meg Ehm | GSK |
| Dawn Waterworth | GSK |
| Caroline Fox | Merck |
| Anders Malarstig | Pfizer |
| Kathy Klinger | Sanofi |
| Kathy Call | Sanofi |

UH & Biobanks

| | |
|---|---|
| Tomi Mäkelä | University of Helsinki / FIMM |
| Jaakko Kaprio | University of Helsinki / FIMM |
| Petri Virolainen | Auria BB / Univ. of Turku /VSSHP |
| Kari Pulkki | Auria BB / Univ. of Turku /VSSHP |
| Terhi Kilpi | THL Biobank (BB) / THL |
| Markus Perola | THL Biobank (BB) / THL |
| Jukka Partanen | Finnish Red Cross Blood Service/FHRB |
| Anne Pitkäranta | HUS/Univ Hosp Districts |
| Riitta Kaarteenaho | Borealis BB/Univ. of Oulu/PPSHP |
| Seppo Vainio | Borealis BB/Univ. of Oulu/PPSHP |
| Kimmo Savinainen | Tampere BB/Univ Tampere/PSHP |
| Veli-Matti Kosma | Eastern Finland BB/UEF/PSSHP |
| Urho Kujala | Central Finland BB /UJy/KSSHP |

Other Experts/ Non-Voting Members

| | |
|---|---|
| Outi Tuovila | Business Finland |
| Minna Hendolin | Business Finland |
| Raimo Pakkanen | Business Finland |

Scientific Committee

Pharma

| | |
|---|---|
| Jeff Waring | Abbvie |
| Bridget Riley-Gillis | AbbVie |
| Athena Matakidou | Astra Zeneca |
| Heiko Runz | Biogen |
| Jimmy Liu | Biogen |
| Shameek Biswas | Celgene |
| Julie Hunkapiller | Genentech |
| Dawn Waterworth | GSK |
| Meg Ehm | GSK |
| Josh Hoffman | GSK |
| Dorothee Diogo | Merck |
| Caroline Fox | Merck |
| Anders Malarstig | Pfizer |
| Catherine Marshall | Pfizer |
| Xinli Hu | Pfizer |
| Kathy Call | Sanofi |
| Kathy Klinger | Sanofi |

UH & Biobanks

| | |
|---|---|
| Samuli Ripatti | University of Helsinki / FIMM |
| Johanna Schleutker | Auria BB / Univ. of Turku /VSSHP |
| Markus Perola | THL Biobank (BB) / THL |
| Tiina Wahlfors | Finnish Red Cross Blood Service/FHRB |
| Olli Carpen | HUS/Univ Hosp Districts |
| Johanna Myllyharju | Borealis BB/Univ. of Oulu/PPSHP |
| Johannes Kettunen | Borealis BB/Univ. of Oulu/PPSHP |
| Reijo Laaksonen | Tampere BB/Univ Tampere/PSHP |
| Arto Mannermaa | Eastern Finland BB/UEF/PSSHP |
| Juha Paloneva | Central Finland BB /UJy/KSSHP |
| Urho Kujala | Central Finland BB /UJy/KSSHP |

Other Experts/ Non-Voting Members

| | |
|---|---|
| Outi Tuovila | Business Finland |
| Minna Hendolin | Business Finland |
| Raimo Pakkanen | Business Finland |

Clinical Groups

Neurology Group

| | |
|---|---|
| Hilkka Soininen | LEAD: Kuopio |
| Valtteri Julkunen | Kuopio |
| Anne Remes | Oulu |
| Reetta Kälviäinen | Kuopio |
| Mikko Hiltunen | Kuopio |
| Jukka Peltola | Tampere |
| Pentti Tienari | Helsinki |
| Juha Rinne | Turku |
| Adam Ziemann | AbbVie |
| Jeffrey Waring | AbbVie |
| Sahar Esmaeeli | AbbVie |
| Nizar Smaoui | AbbVie |
| Anne Lehtonen | AbbVie |
| Susan Eaton | Biogen |
| Heiko Runz | Biogen |
| Sanni Lahdenperä | Biogen |
| Janet van Adelsberg | Celgene |
| Shameek Biswas | Celgene |
| John Michon | Genentech |
| Geoff Kerchner | Genentech |
| Julie Hunkapiller | Genentech |
| Natalie Bowers | Genentech |
| Edmond Teng | Genentech |
| John Eicher | Merck |
| Vinay Mehta | Merck |
| Padhraig Gormley | Merck |
| Kari Linden | Pfizer |
| Christopher Whelan | Pfizer |
| Fanli Xu | GSK |
| David Pulford | GSK |

Gastroenterology Group

| | |
|---|---|
| Martti Färkkilä | LEAD:Helsinki |
| Sampsa Pikkarainen | HUS |

| | |
|---|---|
| Airi Jussila | Tampere |
| Timo Blomster | Oulu |
| Mikko Kiviniemi | Kuopio |
| Markku Voutilainen | Turku |
| Bob Georgantas | AbbVie |
| Graham Heap | AbbVie |
| Jeffrey Waring | AbbVie |
| Nizar Smaoui | AbbVie |
| Fedik Rahimov | AbbVie |
| Anne Lehtonen | AbbVie |
| Keith Usiskin | Celgene |
| Tim Lu | Genentech |
| Natalie Bowers | Genentech |
| Danny Oh | Genentech |
| John Michon | Genentech |
| Vinay Mehta | Merck |
| Dermot Reilly | Merck |
| Kirsi Kalpala | Pfizer |
| Melissa Miller | Pfizer |
| Xinli Hu | Pfizer |
| Linda McCarthy | GSK |

Rheumatology Group

| | |
|---|---|
| Kari Eklund | LEAD:Helsinki |
| Antti Palomäki | Turku |
| Pia Isomaki | Tampere |
| Laura Pirilä | Turku |
| Oili Kaipiainen-Seppänen | Kuopio |
| Johanna Huhtakangas | Oulu |
| Bob Georgantas | AbbVie |
| Jeffrey Waring | AbbVie |
| Fedik Rahimov | AbbVie |
| Apinya Lertratanakul | AbbVie |
| Nizar Smaoui | AbbVie |
| Anne Lehtonen | AbbVie |
| David Close | AstraZeneca |
| Marla Hochfeld | Celgene |
| Natalie Bowers | Genentech |
| John Michon | Genentech |
| Dorothee Diogo | Merck |
| Vinay Mehta | Merck |
| Kirsi Kalpala | Pfizer |

| | |
|---|---|
| Nan Bing | Pfizer |
| Xinli Hu | Pfizer |
| Jorge Esparza Gordillo | GSK |
| Nina Mars | University of Helsinki / FIMM |

Pulmonology Group

| | |
|---|---|
| Tarja Laitinen | LEAD:Tampere |
| Margit Pelkonen | Kuopio |
| Paula Kauppi | Helsinki |
| Hannu Kankaanranta | Tampere |
| Terttu Harju | Oulu |
| Nizar Smaoui | AbbVie |
| David Close | AstraZeneca |
| Steven Greenberg | Celgene |
| Hubert Chen | Genentech |
| Natalie Bowers | Genentech |
| John Michon | Genentech |
| Vinay Mehta | Merck |
| Jo Betts | GSK |
| Soumitra Ghosh | GSK |

Cardiometabolic Diseases Group

| | |
|---|---|
| Veikko Salomaa | Lead: THL |
| Teemu Niiranen | THL |
| Markus Juonala | Turku |
| Kaj Metsärinne | Turku |
| Mika Kähönen | Tampere |
| Juhani Junttila | Oulu |
| Markku Laakso | Kuopio |
| Jussi Pihlajamäki | Kuopio |
| Juha Sinisalo | Helsinki |
| Marja-Riitta Taskinen | Helsinki |
| Tiinamaija Tuomi | Helsinki |
| Jari Laukkanen | Keski-Suomen Keskussairaala |
| Ben Challis | AstraZeneca |
| Keith Usiskin | Celgene |
| Andrew Peterson | Genentech |
| Julie Hunkapiller | Genentech |
| Natalie Bowers | Genentech |
| John Michon | Genentech |
| Dorothee Diogo | Merck |
| Dermot Reilly | Merck |

| | |
|---|---|
| Audrey Chu | Merck |
| Vinay Mehta | Merck |
| Jaakko Parkkinen | Pfizer |
| Melissa Miller | Pfizer |
| Anthony Muslin | Sanofi |
| Dawn Waterworth | GSK |

Oncology Group

| | |
|---|---|
| Heikki Joensuu | Lead: Helsinki |
| Tuomo Meretoja | Helsinki |
| Olli Carpen | Helsinki |
| Lauri Aaltonen | Helsinki |
| Annika Auranen | Tampere |
| Peeter Karihtala | Oulu |
| Saila Kauppila | Oulu |
| Päivi Auvinen | Kuopio |
| Klaus Elenius | Turku |
| Relja Popovic | AbbVie |
| Jeffrey Waring | AbbVie |
| Bridget Riley-Gillis | AbbVie |
| Anne Lehtonen | AbbVie |
| Athena Matakidou | AstraZeneca |
| Jennifer Schutzman | Genentech |
| Julie Hunkapiller | Genentech |
| Natalie Bowers | Genentech |
| John Michon | Genentech |
| Vinay Mehta | Merck |
| Andrey Loboda | Merck |
| Aparna Chhibber | Merck |
| Heli Lehtonen | Pfizer |
| Stefan McDonough | Pfizer |
| Marika Crohns | Sanofi |
| Diptee Kulkarni | GSK |

Opthalmology Group

| | |
|---|---|
| Kai Kaarniranta | Lead: Kuopio |
| Joni Turunen | HUS/ Secretary |
| Terhi Ollila | HUS |
| Sanna Seitsonen | HUS |
| Hannu Uusitalo | Tampere |
| Vesa Aaltonen | Turku |
| Hannele Uusitalo-Järvinen | PSHP |
| Marja Luodonpää | Oulu |

| | |
|---|---|
| Nina Hautala | Oulu |
| Heiko Runz | Biogen |
| Erich Strauss | Genentech |
| Natalie Bowers | Genentech |
| Hao Chen | Genentech |
| John Michon | Genentech |
| Anna Podgornaia | Merck |
| Vinay Mehta | Merck |
| Dorothee Diogo | Merck |
| Joshua Hoffman | GSK |

Dermatology Group

| | |
|---|---|
| Kaisa Tasanen | Oulu |
| Laura Huilaja | Oulu |
| Katariina Hannula-Jouppi | HUS |
| Teea Salmi | Tampere |
| Sirkku Peltonen | Turku |
| Leena Koulu | Turku |
| Ilkka Harvima | Kuopio |
| Kirsi Kalpala | Pfizer |
| Ying Wu | Pfizer |
| David Choy | Genentech |
| John Michon | Genentech |
| Nizar Smaoui | AbbVie |
| Fedik Rahimov | AbbVie |
| Anne Lehtonen | AbbVie |
| Dawn Waterworth | GSK |

FinnGen Teams

Administration Team

| | |
|---|---|
| Anu Jalanko | University of Helsinki / FIMM |
| Risto Kajanne | University of Helsinki / FIMM |
| Ulrike Lyhs | University of Helsinki / FIMM |

Communication

| | |
|---|---|
| Mari Kaunisto | University of Helsinki / FIMM |

Analysis Team

| | |
|---|---|
| Justin Wade Davis | Abbvie |
| Bridget Riley-Gillis | Abbvie |
| Danjuma Quarless | Abbvie |
| Slavé Petrovski | Astra Zeneca |
| Jimmy Liu | Biogen |
| Chia-Yen Chen | Biogen |
| Paola Bronson | Biogen |
| Robert Yang | Celgene |
| Joseph Maranville | Celgene |
| Shameek Biswas | Celgene |
| Diana Chang | Genentech |
| Julie Hunkapiller | Genentech |
| Tushar Bhangale | Genentech |
| Natalie Bowers | Genentech |
| Dorothee Diogo | Merck |
| Emily Holzinger | Merck |
| Padhraig Gormley | Merck |
| Xulong Wang | Merck |
| Xing Chen | Pfizer |
| Åsa Hedman | Pfizer |
| Joshua Hoffman | GSK |
| Clarence Wang | Sanofi |
| Ethan Xu | Sanofi |
| Franck Auge | Sanofi |
| Clement Chatelain | Sanofi |
| Mitja Kurki | University of Helsinki / FIMM/ Broad Institute |
| Samuli Ripatti | University of Helsinki / FIMM |
| Mark Daly | University of Helsinki / FIMM |
| Juha Karjalainen | University of Helsinki / FIMM/ Broad Institute |
| Aki Havulinna | University of Helsinki / FIMM |
| Anu Jalanko | University of Helsinki / FIMM |
| Kimmo Palin | University of Helsinki |
| Priit Palta | University of Helsinki / FIMM |
| Pietro della Briotta Parolo | University of Helsinki / FIMM |
| Wei Zhou | Broad Institute |
| Susanna Lemmelä | University of Helsinki / FIMM |
| Manuel Rivas | University of Stanford |
| Jarmo Harju | University of Helsinki / FIMM |
| Aarno Palotie | University of Helsinki / FIMM |
| Arto Lehisto | University of Helsinki / FIMM |
| Andrea Ganna | University of Helsinki / FIMM |
| Vincent Llorens | University of Helsinki / FIMM |
| Antti Karlsson | Auria BB / Univ. of Turku /VSSHP |

| | |
|---|---|
| Kati Kristiansson | THL BB / THL |
| Mikko Arvas | Finnish Red Cross Blood Service BB /FHRB |
| Kati Hyvärinen | Finnish Red Cross Blood Service BB /FHRB |
| Jarmo Ritari | Finnish Red Cross Blood Service BB /FHRB |
| Tiina Wahlfors | Finnish Red Cross Blood Service BB /FHRB |
| Miika Koskinen | Helsinki BB/HUS/Univ Hosp Districts |
| Olli Carpen | Helsinki BB/HUS/Univ Hosp Districts |
| Johannes Kettunen | Borealis BB/Univ. of Oulu/PPSHP |
| Katri Pylkäs | Borealis BB/Univ. of Oulu/PPSHP |
| Marita Kalaoja | Borealis BB/Univ. of Oulu/PPSHP |
| Minna Karjalainen | Borealis BB/Univ. of Oulu/PPSHP |
| Tuomo Mantere | Borealis BB/Univ. of Oulu/PPSHP |
| Eeva Kangasniemi | Tampere BB/Univ Tampere/PSHP |
| Sami Heikkinen | Eastern Finland BB/UEF/PSSHP |
| Arto Mannermaa | Eastern Finland BB/UEF/PSSHP |
| Eija Laakkonen | Central Finland BB /UJy/KSSHP |
| Juha Kononen | Central Finland BB /UJy/KSSHP |

Sample Collection Coordination

| | |
|---|---|
| Anu Loukola | Helsinki BB/HUS/Univ Hosp Districts |

Sample Logistics

| | |
|---|---|
| Päivi Laiho | THL BB / THL |
| Tuuli Sistonen | THL BB / THL |
| Essi Kaiharju | THL BB / THL |
| Markku Laukkanen | THL BB / THL |
| Elina Järvensivu | THL BB / THL |
| Sini Lähteenmäki | THL BB / THL |
| Lotta Männikkö | THL BB / THL |
| Regis Wong | THL BB / THL |

Registry Data Operations

| | |
|---|---|
| Kati Kristiansson | THL BB / THL |
| Hannele Mattsson | THL BB / THL |
| Susanna Lemmelä | University of Helsinki / FIMM |
| Tero Hiekkalinna | THL BB / THL |
| Manuel González Jiménez | THL BB / THL |

Genotyping

| | |
|---|---|
| Kati Donner | University of Helsinki / FIMM |

Sequencing Informatics

| | |
|---|---|
| Priit Palta | University of Helsinki / FIMM |
| Kalle Pärn | University of Helsinki / FIMM |
| Javier Nunez-Fontarnau | University of Helsinki / FIMM |

Data Management and IT Infrastructure

| | |
|---|---|
| Jarmo Harju | University of Helsinki / FIMM |
| Elina Kilpeläinen | University of Helsinki / FIMM |
| Timo P. Sipilä | University of Helsinki / FIMM |
| Georg Brein | University of Helsinki / FIMM |
| Alexander Dada | University of Helsinki / FIMM |
| Ghazal Awaisa | University of Helsinki / FIMM |
| Anastasia Shcherban | University of Helsinki / FIMM |
| Tuomas Sipilä | University of Helsinki / FIMM |

Clinical Endpoint Development

| | |
|---|---|
| Hannele Laivuori | University of Helsinki / FIMM |
| Aki Havulinna | University of Helsinki / FIMM |
| Susanna Lemmelä | University of Helsinki / FIMM |
| Tuomo Kiiskinen | University of Helsinki / FIMM |

Trajectory Team

| | |
|---|---|
| Tarja Laitinen | Tampere University Hospital |
| Harri Siirtola | University of Tampere |
| Javier Gracia Tabuenca | University of Tampere |

Biobank Directors

| | |
|---|---|
| Lila Kallio | Auria Biobank |
| Sirpa Soini | THL Biobank |
| Jukka Partanen | Blood Service Biobank |
| Kimmo Pitkänen | Helsinki Biobank |
| Seppo Vainio | Northern Finland Biobank Borealis |
| Kimmo Savinainen | Tampere Biobank |
| Veli-Matti Kosma | Biobank of Eastern Finland |
| Teijo Kuopio | Central Finland Biobank |

### Supplementary Acknowledgments

We thank the International Genomics of Alzheimer's Project (IGAP) for providing summary results data for these analyses. The investigators within IGAP contributed to the design and implementation of IGAP and/or provided data but did not participate in analysis or writing of this report. IGAP was made possible by the generous participation of the control subjects, the patients, and their families. The i–Select chips was funded by the French National Foundation on Alzheimer's disease and related disorders. EADI was supported by the LABEX (laboratory of excellence program investment for the future) DISTALZ grant, Inserm, Institut Pasteur de Lille, Université de Lille 2 and the Lille University Hospital. GERAD was supported by the Medical Research Council (Grant n° 503480), Alzheimer's Research UK (Grant n° 503176), the Wellcome Trust (Grant n° 082604/2/07/Z) and German Federal Ministry of Education and Research (BMBF): Competence Network Dementia (CND) grant n° 01GI0102, 01GI0711, 01GI0420. CHARGE was partly supported by the NIH/NIA grant R01 AG033193 and the NIA AG081220 and AGES contract N01–AG–12100, the NHLBI grant R01 HL105756, the Icelandic Heart Association, and the Erasmus Medical Center and Erasmus University. ADGC was supported by the NIH/NIA grants: U01 AG032984, U24 AG021886, U01 AG016976, and the Alzheimer's Association grant ADGC–10–196728.

Data on glycaemic traits have been contributed by MAGIC investigators and have been downloaded from www.magicinvestigators.org.

Data on coronary artery disease / myocardial infarction have been contributed by CARDIoGRAMplusC4D investigators and have been downloaded from www.CARDIOGRAMPLUSC4D.ORG'. For the Exome chip study please acknowledge the source of the data as follows: 'Data on coronary artery disease / myocardial infarction have been contributed by the Myocardial Infarction Genetics and CARDIoGRAM Exome investigators and have been downloaded from www.CARDIOGRAMPLUSC4D.ORG'. For the Chromosome X-CAD please acknowledge the source of the data as follows: ‘Data on the X chromosomal analysis of coronary artery disease / myocardial infarction have been contributed by the referenced authors.’ For the UK Biobank meta-analysis please acknowledge the source of the data as follows: 'Data on coronary artery disease / myocardial infarction have been contributed by the CARDIoGRAMplusC4D and UK Biobank CardioMetabolic Consortium CHD working group who used the UK Biobank Resource (application number 9922). Data have been downloaded from www.CARDIOGRAMPLUSC4D.ORG.

We used summary statistics generated by Ben Neale’s lab in UK Biobank, downloaded from https://docs.google.com/spreadsheets/d/1kvPoupSzsSFBNSztMzl04xMoSC3Kcx3CrjVf4yBmESU/edit?ts=5b5f17db#gid=227859291 and https://docs.google.com/spreadsheets/d/1b3oGI2lUt57BcuHttWaZotQcI0-mBRPyZihz87Ms_No/edit#gid=275725118.

We acknowledge the participants and researchers in the Biobank Japan Project.
